# Most biodiversity indicators add little beyond natural land cover

**DOI:** 10.64898/2026.11.10.765737

**Authors:** Taku Kadoya

**Affiliations:** Biodiversity Division, National Institute for Environmental Studies, 16-2 Onogawa, Tsukuba, Ibaraki 305-8506, Japan

## Abstract

Land-use biodiversity indicators increasingly inform progress on the Kunming-Montreal Global Biodiversity Framework and corporate nature-related disclosure, yet few have been tested against independent field data. In standardized bird surveys on three continents, most of seven leading indicators distinguished sites within ecoregions no better than the share of natural land cover, for both species richness and composition. Because many rely on IUCN range maps, which restrict commercial use, I estimated land-cover effects on both from commercially usable GBIF records. Only the composition estimate outperformed natural-cover share for composition in all three regions, the richness estimate matched the IUCN richness map in two, and neither predicted the other quantity. Cropland retained about half of forest bird species but four-fifths of grassland, savanna and shrubland species, whereas its relative richness hardly depended on the reference. Indicators should report richness and composition together, be tested in independent surveys and state a justified reference.

---

Decisions on where to protect, restore or convert land increasingly draw on global indicators of land-use impact on biodiversity, used to monitor the Kunming-Montreal Global Biodiversity Framework and to inform nature-related corporate disclosure^1^. Unless these indicators track what land use does to the species living there, targets monitored with them can be met on paper but not in the field. Seven indicators are especially prominent: two versions of the Biodiversity Intactness Index, from the Natural History Museum (BII (NHM))^2,3^ and from Impact Observatory (BII (IO))^4^, the Biodiversity Habitat Index (BHI)^5^, Mean Species Abundance (MSA)^6^, the Potentially Disappeared Fraction (PDF)^7^, the LIFE extinction-impact metric^8^ and the Species Threat Abatement and Restoration metric (STAR)^9^. They capture different facets of biodiversity, are distributed as maps or as coefficients for each land-cover type, and draw on expert-drawn ranges, habitat classifications and meta-analyses of local studies. A separate product, the IUCN richness map for birds and mammals^10^, stacks each species’ Area of Habitat (AoH)^11^, an expert-drawn range refined by habitat and elevation. Few studies have tested, however, whether these products align with independent field surveys^3^. Many also depend on IUCN range maps, whose terms restrict commercial use, which limits their use in the corporate disclosure they are meant to serve.

Knowing what an indicator captures matters because richness and composition can tell different stories. Time series show no average decline in local richness alongside faster-than-expected compositional turnover^12,13^, whereas spatial comparisons find lower local richness on human-modified land^14^. Site selection may account for part of this discrepancy^15^. Whatever its cause, compositional change does not track richness change in direction or magnitude^16^. An indicator that captures only one of the two will therefore miss changes in the other. Existing indicators weigh these dimensions differently or blend them with others, such as abundance or extinction risk, and how well each tracks field richness and composition has not been tested against independent data.

Open occurrence records could remove the licensing barrier, but their uneven recording effort makes the effects of land cover hard to compare among places. Here I develop two estimates of the effect of land cover, one on species richness and one on composition, from public GBIF occurrence records licensed under CC0 or CC BY, for birds, mammals, reptiles and amphibians in 5 km cells across the global land surface (Fig. 1a,b). I test these GBIF-based estimates, together with the seven indicators and the IUCN AoH richness map, against standardized bird surveys in three regions that were not used in the estimation (Fig. 1c–e). The baseline is the share of natural land cover in each surveyed area: an indicator that does not outperform it conveys nothing beyond how much natural land remains. Indicators also differ in the natural state against which they compare land use^2,6^, and the State of Nature metrics framework published in 2026 counts semi-natural ecosystems as nature and accepts other justifiable references^17^. I therefore use the estimates to examine globally how the assessment of anthropogenic land depends on the natural reference, and apply them to land-cover change between 2020 and 2024.

**Fig. 1.**
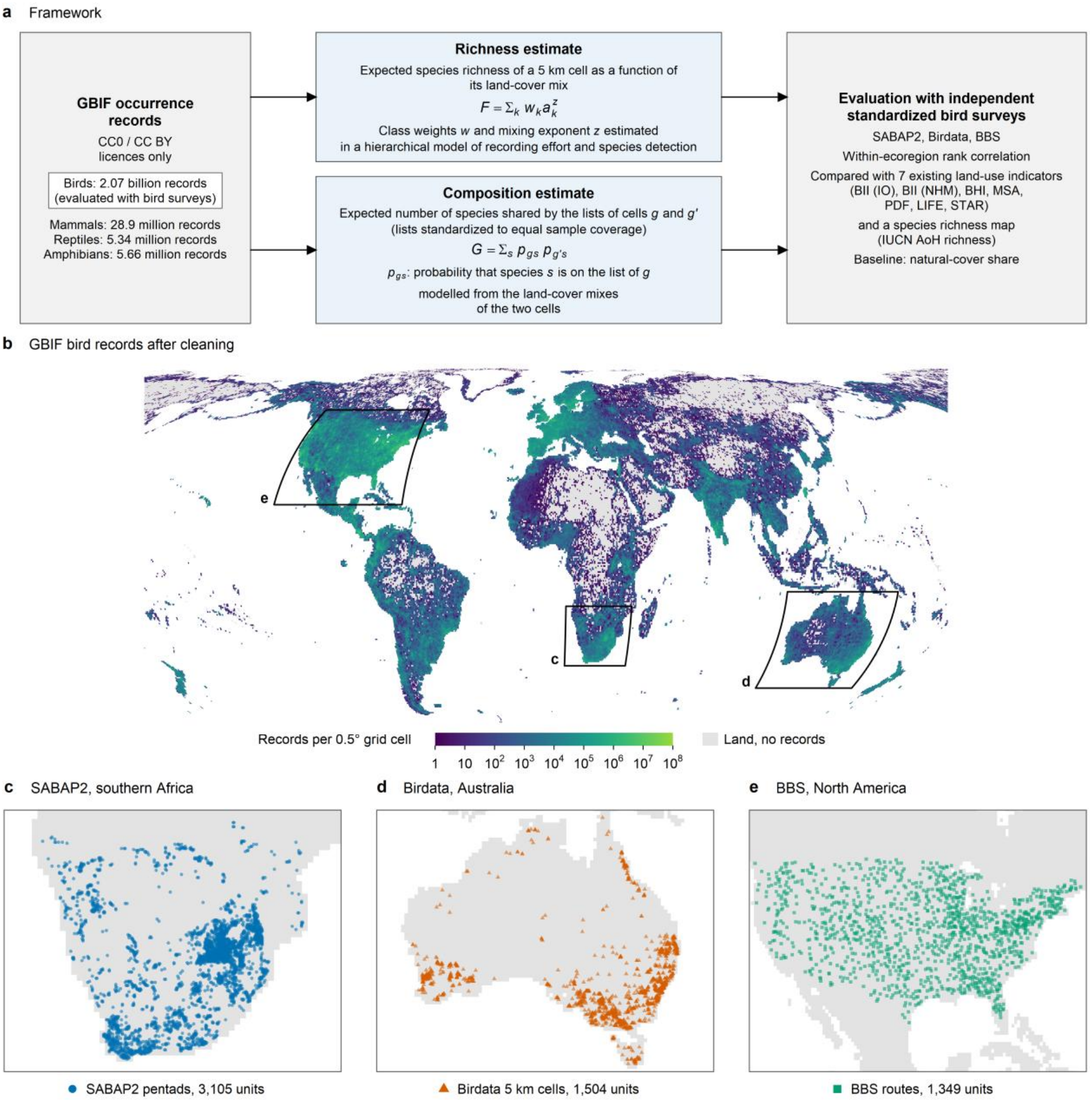
Estimating the effects of land cover from open occurrence records and testing them in independent bird surveys. (a) Records open to commercial use (left) are used to estimate how the land-cover mix of a cell determines its expected richness (top) and the species it shares with another cell (bottom). In the richness estimate, *a*_*k*_ is the area share of land-cover class *k*, the weight *w* is the expected richness of a cell covered entirely by class *k*, and the mixing exponent *z* sets the additional richness gained when classes are mixed. In the composition estimate, *G* is the number of species shared by the lists of two cells, from which compositional similarity and species retention are derived. Both estimates are then tested against standardized bird surveys not used in the estimation (right); birds (inner frame) are the only taxon that could be tested. (b) GBIF bird records after cleaning and before the inclusion criteria (Methods), with the extents of the three surveys; they include atlas records, such as those of SABAP2, that were removed from the estimation inputs. (c)–(e) The survey units used for the test: squares of about 9 km (pentads) in SABAP2, 5 km cells in Birdata and routes in the BBS.

## Results

### Two estimates from public occurrence records

I described land cover with the MODIS 500 m classification, which has 14 natural and three anthropogenic classes (cropland, urban and cropland/natural mosaic), calculated the area share of each class in 5 km cells (hereafter the land-cover mix) and made all estimates within ecoregions^18^.

The richness estimate predicts a cell’s expected richness from its land-cover mix (Fig. 1a, upper box). Its parameters, a weight for each class in each ecoregion (the expected richness of a cell covered entirely by that class) and a mixing exponent (the additional richness gained, beyond the area-weighted mean of the class weights, when classes are mixed), were fitted to GBIF records with a hierarchical model of recording effort and species detection (Methods).

Estimating composition requires species lists that are comparable among cells despite very different recording effort. I therefore built each cell’s list by adding recording days in random order until sample coverage, a measure of how complete the list is^19^, reached the same value in every cell, and then modelled the number of species shared by the lists of two cells in an ecoregion as a function of their land-cover mixes (the sharing model; Fig. 1a, lower box). This model yields species retention (hereafter retention): how much of one cell’s community remains in a cell with a different land-cover mix, relative to a second cell with the same mix as the first (a value of 1 means no difference).

Because both estimates depend only on land-cover mix, they can be calculated for hypothetical cells. In simulated data, the composition estimate recovered its target under varying recording effort; the richness estimate understated anthropogenic richness ratios, so its intervals include model error (Supplementary Note 1).

### Evaluation in independent bird surveys

I evaluated the estimates with three standardized bird surveys that played no part in the estimation (Fig. 1c–e): the Southern African Bird Atlas Project 2 (SABAP2), Birdata in Australia and the North American Breeding Bird Survey (BBS). Their sampling units (squares of about 9 km, 5 km cells and routes) are hereafter survey units. In each unit I measured rarefied richness and field composition similarity, the mean presence/absence Sørensen similarity to natural units of the same ecoregion (survey units with a natural-cover share of at least 0.95). Similarity declines with distance regardless of land cover, so I removed the effects of distance to natural units and of survey effort (Methods).

I compared the two GBIF-based estimates with the seven indicators, the IUCN AoH richness map and, as the baseline, natural-cover share. For each survey unit, the composition estimate is the predicted Sørensen similarity to natural units, relative to that predicted between natural units. Because differences among ecoregions mainly reflect regional species pools rather than land cover, I measured performance as the Spearman rank correlation within ecoregions, after removing ecoregion means (and, for composition, the effects of distance and effort) from both the field quantity and the predictor. Uncertainty is reported as 90% intervals (Methods).

### Richness: the GBIF-based estimate and AoH outperformed natural-cover share; most indicators did not

Most indicators captured almost none of the differences in richness within ecoregions, whereas the GBIF-based estimate and AoH did. Natural-cover share was itself uninformative (rank correlations of −0.10, 0.00 and −0.01 in southern Africa, Australia and North America, respectively; same order below; Fig. 2a), and so were the coefficient versions of MSA, BHI and PDF. MSA and BHI set all natural classes to 1 and assign lower values only to anthropogenic classes, so within an ecoregion they rank units almost as natural-cover share does (rank correlation 0.98–1.00). The rank correlations of the BII and BHI maps were near zero or negative. LIFE was a partial exception, outperforming natural-cover share in southern Africa and North America but not in Australia (0.15, −0.05 and 0.12). The GBIF-based richness estimate reached 0.12, 0.21 and 0.26, higher than all seven indicators in Australia and North America (90% intervals of the differences excluded 0); in southern Africa LIFE was slightly higher (difference −0.04; 90% interval −0.16 to 0.07). By effect size (the variance explained beyond the baseline as a proportion of the maximum explainable by land-cover mix, which can exceed 1; Methods), the estimate exceeded the baseline in two of the three surveys (0.03, 0.76 and 0.77; Extended Data Fig. 1a).

**Fig. 2.**
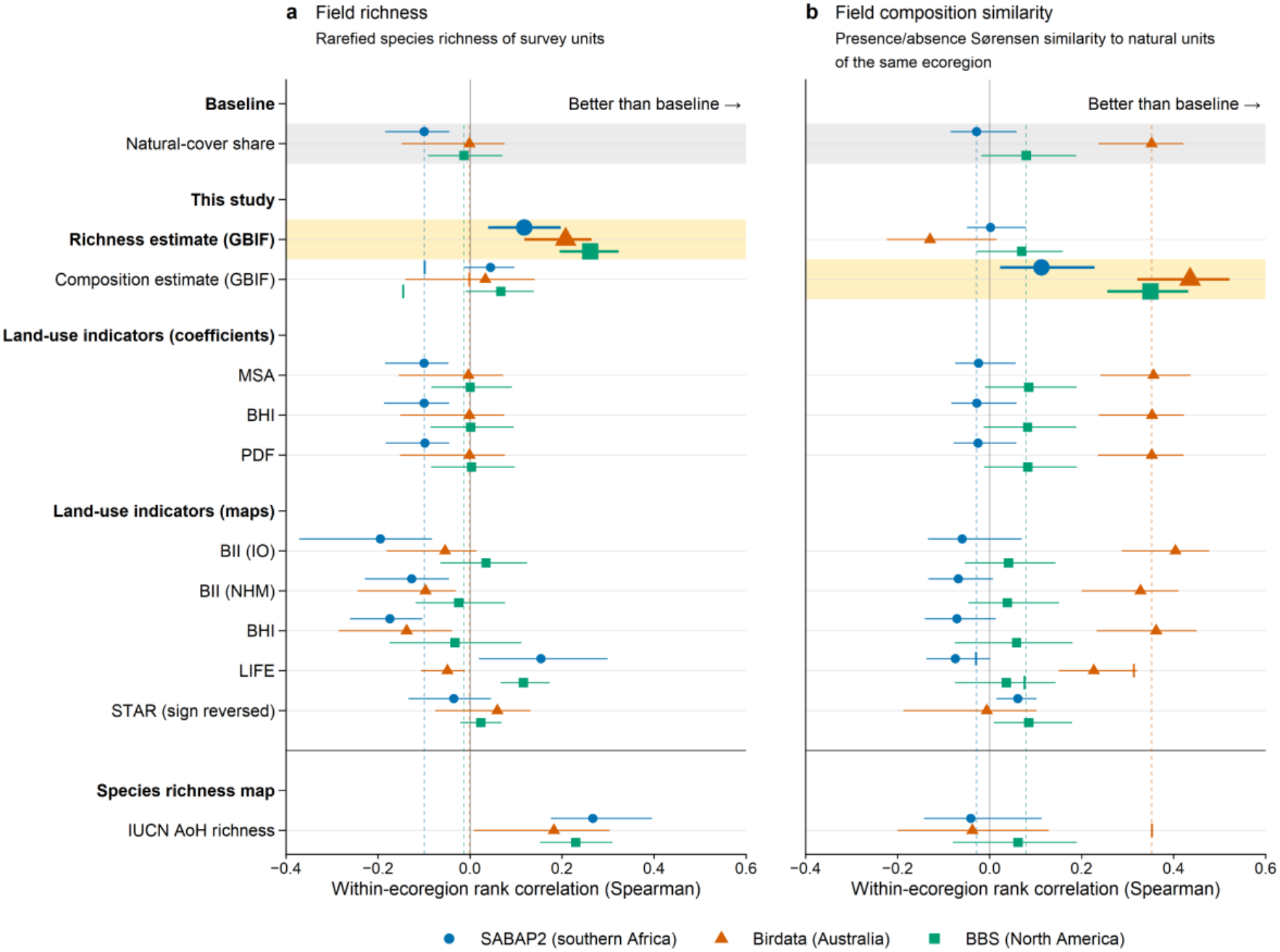
Most existing indicators rank survey units no better than the share of natural land cover, whereas the GBIF-based estimates do better, each for its own target. (a) Field richness, rarefied to a common number of surveys. (b) Field composition similarity, the mean presence/absence Sørensen similarity to natural units of the same ecoregion (survey units with a natural-cover share of at least 0.95), adjusted for survey effort and distance to natural units (Methods). Each point is the within-ecoregion Spearman rank correlation between the field quantity and a predictor in one survey, so points further right rank survey units more correctly; lines are 90% intervals. A point to the right of the dotted line of its colour outperforms natural-cover share (grey band) in that survey. The GBIF-based estimate of each panel (yellow band) lies to the right of the dotted lines in all three surveys. Coefficient indicators lie on the lines, and most map indicators lie on or close to them. IUCN AoH richness, a map of richness itself, outperforms natural-cover share for richness but not for composition. The composition estimate in (a) and LIFE and IUCN AoH richness in (b) use fewer survey units; for these rows, the short vertical mark gives natural-cover share for the same units. Numbers of survey units (southern Africa, Australia, North America) are 3,105, 1,504 and 1,349 in (a) and 3,078, 1,501 and 912 in (b) (Supplementary Table 11 gives the other rows). MSA, mean species abundance; BHI, Biodiversity Habitat Index; PDF, potentially disappeared fraction; BII, Biodiversity Intactness Index (IO, Impact Observatory; NHM, Natural History Museum); STAR, Species Threat Abatement and Restoration.

The most direct comparator is the IUCN AoH richness map, which maps richness itself rather than land-use impact. It outperformed the baseline in all three surveys (0.27, 0.18 and 0.23; Fig. 2a). The GBIF-based estimate performed as well as AoH in Australia and North America, whereas AoH was higher in southern Africa (Supplementary Note 2). The two carry different information: adding the GBIF-based estimate to the AoH map raised the share of variance explained by 0.007, 0.035 and 0.068.

By contrast, STAR, which is built from the AoH of threatened and Near Threatened species and which I rebuilt for these surveys (Methods), captured almost none of the differences in richness (−0.04, 0.06 and 0.02; Fig. 2a). Restricting AoH to these species alone lowered its rank correlations sharply (from 0.27, 0.18 and 0.23 to 0.10, 0.08 and 0.08; Supplementary Note 2). STAR’s mismatch with richness therefore appears to stem from what it is designed to measure, the opportunity to reduce threats to these species, rather than from the quality of the underlying distributions.

### Composition: only the GBIF-based estimate outperformed natural-cover share in all three surveys

The rank correlations of the GBIF-based composition estimate with field composition similarity were 0.11, 0.44 and 0.35, higher than those of all seven indicators in all three surveys (Fig. 2b). Its precision-weighted mean advantage over natural-cover share across surveys was +0.121 in rank correlation (lower end of the 90% interval +0.074; Methods), and its advantages over BII (IO) and the BHI map were +0.098 (lower end +0.060) and +0.147 (lower end +0.087). The BII and BHI maps exceeded natural-cover share by at most 0.05 (BII (IO) clearly only in Australia; precision-weighted advantage +0.028, lower end +0.001), and the coefficient versions of MSA, BHI and PDF were within 0.01 of it. Effect sizes of the composition estimate exceeded the baseline in all three surveys (0.70, 0.83 and 1.79; 90% intervals excluded 0; Extended Data Fig. 1b). The AoH richness map, although it outperformed the baseline for richness, did not do so for composition (−0.04, −0.04 and 0.06, against −0.03, 0.35 and 0.08 for the baseline on the same survey units), and the composition estimate exceeded AoH in all three surveys, although only just in southern Africa (lower ends of the 90% intervals of the difference 0.000, 0.28 and 0.15). The conclusion was the same with Bray–Curtis similarity, which uses abundance information (Extended Data Fig. 2).

### Neither estimate can stand in for the other

As a predictor of composition, the richness estimate captured none of the differences (rank correlations with field composition similarity of 0.00, −0.13 and 0.07); as a predictor of richness, the composition estimate captured almost none (0.04, 0.03 and 0.07; Fig. 2).

### Changing the natural reference changes the compositional assessment

For each ecoregion I built a natural reference cell, a hypothetical cell unlike the real natural units used in the evaluation, in three ways, from all natural classes in proportion to their area, from forest alone and from open natural classes alone (shrubland, savanna and grassland), and compared each with cells covered entirely by one anthropogenic class.

For birds, expected richness on anthropogenic land was lower than in the natural reference cell under every reference: across the three references, the ratio of its expected richness to that of the reference cell ranged from 0.80 to 0.89 for cropland, 0.81 to 0.90 for urban areas and 0.63 to 0.73 for the mosaic.

The cropland and urban values agree well with an analysis of local surveys in the PREDICTS database (−18% for high-intensity cropland and −22% for urban areas relative to primary vegetation)^20^. The compositional assessment of the same land, in contrast, changed greatly with the reference (Fig. 3).

**Fig. 3.**
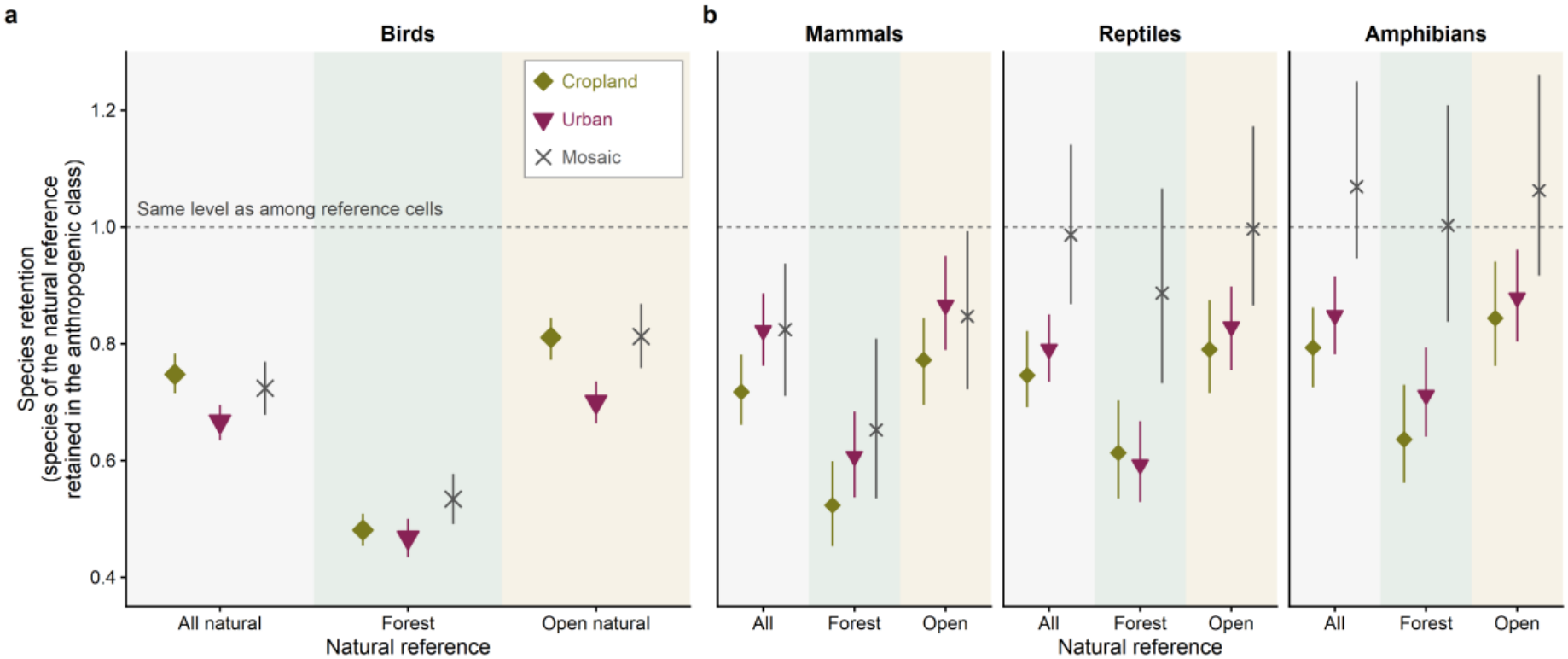
Species retention on anthropogenic land depends strongly on the natural reference. (a) Birds and (b) mammals, reptiles and amphibians. The vertical axis shows how much of the species of a natural reference cell is also found in a cell of an anthropogenic land-cover class, relative to the level among natural reference cells; at the dotted line, the anthropogenic cell holds as many of the reference species as another reference cell does. The natural reference cell is hypothetical and is built from forest alone, from open natural classes alone (shrubland, savanna and grassland) or from all natural classes in proportion to their area in the ecoregion. Bird communities on cropland retain about half of the species of the forest reference but four-fifths of those of the open natural reference, and the same contrast appears in the other taxa. Points are medians across ecoregions and lines are 90% intervals (Methods). Numbers of ecoregions for cropland, urban and mosaic are 283, 232 and 122 for birds, 117, 147 and 47 for mammals, 101, 134 and 40 for reptiles, and 88, 109 and 32 for amphibians.

I compared composition using the retention of the natural reference cell’s species in the anthropogenic cell. A retention of 1 here means that the anthropogenic cell holds as many of the reference species as a second hypothetical cell with the same natural land-cover mix does. For birds, retention against forest was 0.48 (90% interval 0.45–0.51) on cropland, 0.47 in urban areas and 0.53 on the mosaic, whereas against open natural classes it was 0.81 (0.77–0.84), 0.70 and 0.81 (Fig. 3a); against all natural classes it was 0.75, 0.67 and 0.72. On cropland, the highest and lowest values across the three references differed by 0.09 for the richness ratio but by 0.33 for retention. The species of the forest reference are thus the least retained on anthropogenic land, and cropland bird communities are closer to open natural habitats than to forest. Urban communities were further from the natural reference than cropland communities against all natural and open natural classes; against forest, the intervals for the two overlap.

The effect of the reference held across all four taxa, although only birds could be validated in independent surveys. Retention on cropland was 0.48–0.64 against forest and 0.77–0.84 against open natural classes, and in urban areas 0.47–0.71 and 0.70–0.88 (Fig. 3). In ecoregions with values for both the mosaic and cropland, the mosaic retained more of the species of the forest reference than cropland in all four taxa (differences in retention of +0.08 to +0.35; all 90% intervals excluded 0, with lower ends of +0.02 for birds and reptiles; Supplementary Note 3). For species of the open natural reference, the intervals of the difference between the mosaic and cropland included 0 in all four taxa.

Read in the opposite direction, retention shows how many of the species of an anthropogenic cell also occur in the natural reference cell, relative to a second cell of the same class (a value of 1 means as often). For birds against all natural classes, the retention of the anthropogenic cell’s species in the natural reference cell was 0.83 (0.79–0.89) in urban areas, 0.90 (0.86–0.95) on cropland and 1.02 (0.94–1.12) on the mosaic (Extended Data Fig. 3): mosaic species were found in the natural reference as often as in other mosaic cells, whereas urban areas tended to hold a larger share of species not found in the natural reference than cropland did.

### Land-cover-driven change from 2020 to 2024: richness and composition differ in spatial pattern

Between 2020 and 2024, richness and composition differed less in their global mean change than in the spatial pattern of that change. From each cell’s land-cover mix in 2020 and 2024, I calculated the change in expected richness and the compositional change ratio, the share of the 2020 community’s species predicted to be found in 2024 relative to no change (Fig. 4 for birds; Extended Data Fig. 4 for the other three taxa). The ratio exceeds 1 when the 2024 land cover is predicted to share more species with the 2020 community than unchanged land cover does. For birds, the global mean change relative to 2020 was −0.070% in expected richness (90% interval −0.094 to −0.046%) and −0.075% in composition (−0.117 to −0.031%; Methods). Of the 500 m pixels that changed, 88% changed among natural classes, which may reflect real change or classification noise; these changes produce most of the fine-scale variation in Fig. 4a,b. At a resolution of 0.5°, the two maps had a rank correlation of 0.51, and their signs were opposite in 28% of the grid cells in which either changed.

**Fig. 4.**
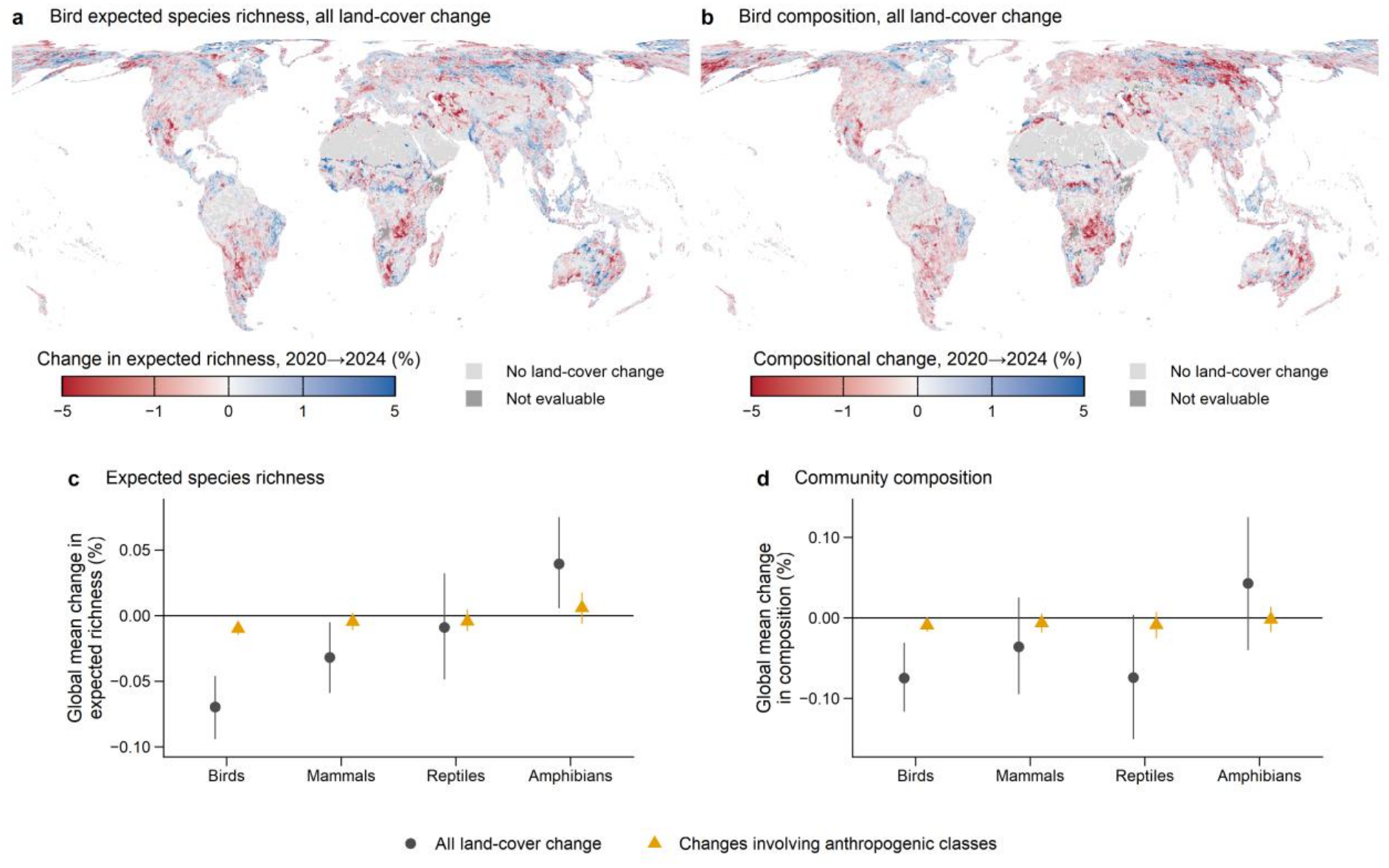
Global changes in richness and composition predicted from land-cover change between 2020 and 2024 are small on average, but the two differ in spatial pattern. (a) Change in the expected richness of birds. (b) Compositional change of birds: the share of the species of the 2020 community predicted to be found in 2024, relative to no change. Both maps include all land-cover change, and much of their fine-scale variation arises from transitions among natural classes. Each 0.5° grid cell (about 50 km) shows the mean of its 5-km cells with values (in (a), cells without land-cover change count as zero). Pale grey (No land-cover change) marks grid cells with no 5-km cell whose land cover changed, and dark grey (Not evaluable) marks grid cells with changed cells but none with a value. (c), (d) Global mean change in expected richness and composition for four taxa, for all land-cover change (dark grey circles) and for changes to or from anthropogenic classes (orange triangles), which are smaller still. Points are estimates and vertical lines are 90% intervals. Global means use only cells supported by data (Methods), so averaging the maps does not reproduce them. In (b), cells in ecoregions without a composition fit take the mean value of the same biome (see Supplementary Fig. 3).

In the other taxa, the global mean change in expected richness ranged from −0.032% to +0.039%, with intervals excluding 0 for mammals and amphibians (Fig. 4c); the amphibian increase came mainly from reclassification between savanna and forest and from conversion of grassland to cropland (Supplementary Note 4). The change in composition ranged from −0.074% to +0.043%, with intervals including 0 (Fig. 4d). Restricting the analysis to changes involving anthropogenic classes made the changes smaller still (for birds, −0.010% in expected richness and −0.009% in composition; Fig. 4c,d; Supplementary Note 4).

## Discussion

### Testing indicators against independent surveys

Indicators used in target tracking and disclosure should be tested against independent surveys, with the share of natural land cover as the minimum baseline. Most of those tested here did not pass it, partly because of their form: many express similarity or ratios of richness or abundance as coefficients for each land-cover type, often treating natural land cover as a single state, so within an ecoregion they distinguish little beyond how much natural land remains. The coefficients continue to be updated^6,7,21^, but I found no update that estimates them directly from occurrence records and tests them in independent surveys. My results argue for re-estimating such coefficients directly from species records, with separate values for natural classes in each ecoregion.

Occurrence records can support this. The GBIF-based richness estimate performed as well as the IUCN AoH richness map, in two of three regions, showing that occurrence records can yield information approaching that of maps compiled from expert knowledge. AoH’s strength lies in location information within ecoregions (range edges and elevation; Supplementary Note 2), so combining the two should improve richness estimates where the licence terms of AoH allow; the GBIF-based estimate alone remains open to commercial use.

Because the estimates take the form of coefficients, they can be applied directly to land-cover change (Fig. 4): an indicator could hold the richness ratio and the retention of the natural community for each land-cover class and ecoregion, and calculate values for any land-cover mix. Independent evaluation can provide the evidence of performance that is reportedly missing when companies choose indicators for disclosure^22^, and the coefficients can separate contributions by type of transition and by region (Supplementary Note 4), helping to identify effective interventions.

### Reporting richness and composition against a stated reference

Because richness and composition cannot substitute for each other, indicators need to report them as a pair. This agrees with calls for multiple complementary indicators^23^ and with evidence that richness, abundance and evenness are often decoupled among sites^24^. Unlike earlier work, this study estimated both quantities from the same data and against a common natural reference across global landscapes that include converted land, and tested each against its field counterpart in independent surveys. The Essential Biodiversity Variables framework calls for monitoring species populations and community composition separately^25^ and for correcting biases in recording effort and detection^26^; the two estimates here show one way to do so from global opportunistic records.

Reporting richness and composition as a pair is not enough on its own: the natural reference against which they are measured should be stated and justified. The same land can look heavily altered or largely intact: bird retention on cropland was 0.48 against forest but 0.81 against open natural classes, although the richness ratio changed little (Fig. 3), so the dependence is invisible when only richness is monitored. The reference also sets the direction of action: measured against forest, tree planting in grassy biomes would count as recovery, even though it can devastate their biodiversity^27^. The choice of baseline can even reverse the sign of the estimated effect of land use on species richness^28^. Many indicators have used primary vegetation^2^ or an undisturbed state^6^, yet most land now regarded as natural has a long history of human use^29^. The State of Nature framework accepts a natural or nearnatural state, the best attainable state or another justifiable reference^17^, and my results provide global support for its call to justify that choice. Because the estimates depend only on land-cover mix, richness and composition can be quantified against any explicitly chosen reference, which provides a basis for indicators that state and justify theirs.

### Semi-natural ecosystems as references

The choice of reference also bears on semi-natural ecosystems, which often combine farmland with natural vegetation and in which long-term human use has maintained habitats, such as wetlands, grasslands and open woodlands, together with the species that depend on them^30^. Where such landscapes are valued for conservation, they may serve as a reference and a target for land management, rather than counting only as altered land. A landscape that keeps more of the natural community than cropland and is not dominated by species absent from natural land is a candidate for that role. Even at the coarse level of MODIS land cover, the cropland/natural mosaic, the class closest to semi-natural ecosystems, met both conditions globally: it retained more of the forest reference’s species than cropland did in all four taxa (Supplementary Note 3), and its bird species were found in natural reference cells as often as in other mosaic cells, unlike those of cropland and urban areas (Extended Data Fig. 3). These patterns accord with countryside biogeography^31,32^, a meta-analysis of landscape complexity^33^, a synthesis for mammals^34^ and studies of Japanese satoyama^30,35^. The mosaic class is only a coarse proxy, not equivalent to semi-natural ecosystems such as satoyama, and is not itself a conservation target. Nor does this study resolve the debate over land sharing and land sparing^36^ or contradict the finding that no landscape can substitute for primary forest^37^. Treating semi-natural ecosystems as part of nature is nevertheless the direction taken by the State of Nature framework^17^.

Methods that score land only as natural or anthropogenic, such as natural-cover share and coefficients that set all natural classes to 1, cannot follow that direction: they count semi-natural landscapes as anthropogenic land and can underestimate their value. The estimates here instead give the mosaic its own richness and retention values.

### Validation data and updatable indicators

Many limitations of this study stem from the scarcity of validation data. Temporal change could not be tested with independent data. Large-scale monitoring schemes are concentrated on birds, butterflies and plants and in a few regions^38^, and the large-scale accumulation of opportunistic records is recent (GBIF records increased 12-fold from 2007 to 2019)^39^. More records^40^ and citizen science that collects contextual information on how observations are made^41^ will improve the precision of biascorrecting estimation. Because the estimates can be rebuilt as records accumulate, they can be extended to other taxa and poorly recorded regions, and their predictions of temporal change can be tested independently. Large public biodiversity datasets, open science and bias-correcting estimation are becoming the foundation for measuring progress towards, and ultimately achieving, nature-positive goals^42^. Paired, tested and updatable indicators of richness and composition built on that foundation can tie the tracking of these goals to what happens to species on the ground.

## Methods

### Data

#### Occurrence records

I downloaded GBIF records of birds, mammals, amphibians and reptiles (Squamata, Testudines, Crocodylia and Rhynchocephalia) in September 2026^43–46^, restricted to records licensed under CC0 1.0 or CC BY 4.0 that had coordinates and no geospatial issue flags. I kept presence records identified to species, excluded fossil and captive records, and assigned the remainder to land cells of a 0.05° latitude–longitude grid (about 5.6 km at the equator; hereafter 5 km cells). Records of the same species in the same cell and date unit (year, month and day, or year and month or year alone when the finer fields were missing) were merged, and each date unit with records in a cell is called a recording day.

#### Inclusion criteria

I excluded records from years in which the land cover of the cell may have differed from that in 2020. Using only information held by GBIF, I also excluded three groups of records that cannot be treated as community surveys: areal records (atlases and compilations that place the species list of a grid cell at its centre), surveys targeting particular species (from 0.6% of records for birds to 15.1% for mammals), and records of marine mammals or extinct species or more than 2,000 km from the ecoregions where records of the species are concentrated (Supplementary Note 5). Exotic species are included, but their influence is small (Supplementary Note 6). Fig. 1a shows the number of records after cleaning; Supplementary Table 9 breaks down the data used in the estimation.

#### Land cover and ecoregions

I used the IGBP classification of MODIS MCD12Q1 Collection 6.1 (17 classes at 500 m)^47^. The 17 classes *k* were divided into anthropogenic classes *h* (cropland [12], urban and cropland/natural mosaic [14]) and the set of natural classes *K*_nat_: forest (1–5), shrubland (6, 7), savanna (8, 9), grassland (10), wetland (11), snow and ice (15), barren land (16) and water (17). The land-cover mix *a* of a cell (the area shares of the 17 classes, summing to 1) is the share of each class in 2020 among all 500 m pixels overlapping the cell, and the mixes of survey units were built from the same 2020 classification (Supplementary Note 7). I write the natural-cover share as 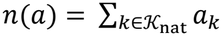 and call real cells and survey units with *n*(*a*) ≥ 0.95 natural cells and natural units (the natural reference cell introduced below is a separate, hypothetical cell). Each cell was assigned to the RESOLVE Ecoregions 2017 ecoregion *e* covering most of its area (846 ecoregions in 14 biomes)^18^.

### Richness estimate

#### Expected richness as a function of the land-cover mix

I expressed the expected richness of a cell in ecoregion *e* as a function of its land-cover mix *a* (Fig. 1a):

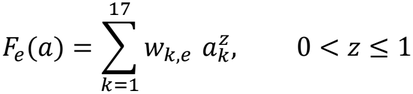

The sum is over the 17 classes. The weight *w*_*k,e*_ is the expected richness of a cell consisting only of class *k*, and the mixing exponent *z* sets how much richness is added when classes are mixed: the expected richness of a cell split equally between two classes is 2^1−*z*^ times the mean of the two weights. Because *F*_*e*_(*a*) depends only on the land-cover mix, it can also be calculated for hypothetical cells. This form is a special case of the countryside species–area relationship^48^ in which each class has its own group of species and all classes share one exponent. Because species are in fact shared among classes, *z* is an empirical quantity that absorbs both area effects and the sharing of species among classes. The form applies the same *z* to every pair of classes and builds in no difference between natural and anthropogenic classes in how species are shared. Since this study compares change in richness with compositional turnover, I chose this symmetric form so that the relationship under study is not built into the equation. A generalization with a separate exponent for mixtures of anthropogenic classes, tested on simulated data, reduced the bias in the ratio of expected richness between anthropogenic and natural cells by only 20–30% and could not be estimated for most ecoregions with the available records, so I did not use it (Supplementary Note 1).

#### Observation model

Observed richness falls short of true richness to a degree that depends on the volume of records. I therefore used a hierarchical model that estimates true richness 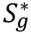 and the recording rate of each species from the number of recording days of cell *g, T*_*g*_, and the number of those days on which species *s* was recorded, *x*_*gs*_ (Supplementary Note 8). True richness follows a Poisson distribution with mean *M*_*g*_, the species-pool intensity of the cell, which allows richness to vary among cells with the same mix. *M*_*g*_ follows a gamma distribution with shape *ξ*_*e*_ and mean 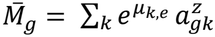, where 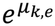 is the true expected richness of a cell consisting only of class *k*. The daily recording rate of a species, *ν*_*gs*_, follows a gamma distribution with shape *κ*_*e*_ and scale *θ*_*g*_ (the recording intensity of the cell, modelled as a normal random effect on the log scale). Because each species is recorded at least once in *T*_*g*_ days with probability *q*_*g*_, observed richness *S*_*g*_ follows a negative binomial distribution with mean 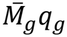.

#### Weights

Estimates of the true expected richness 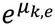 rely more heavily on extrapolation where records are sparse. I therefore defined the weight as the expected number of species found when a cell is surveyed until sample coverage reaches 0.5 (sample coverage is the expected share of the species occurrences recorded on a given day in that cell that belong to species already on the list)^19^:

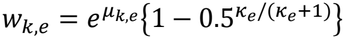

The term in braces is common within an ecoregion, so ratios of weights within an ecoregion equal ratios of true expected richness. I chose 0.5 because the records of many cells reach this level, so little extrapolation is required. Expected richness in the main text and below is calculated from these weights.

#### Sharing information within biomes

To stabilize estimates for ecoregion–class combinations with few records, class effects *β*_*m,k*_ were shared within each biome *m*(*e*):

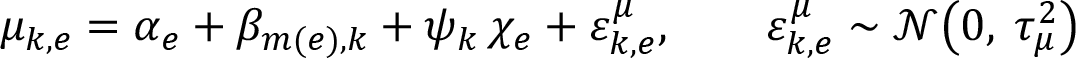

Here *α*_*e*_ is the level of the ecoregion, *χ*_*e*_ its recording density, *ψ*_*k*_ the effect of that density and *τ*_*µ*_ the between-ecoregion variation (Supplementary Note 8). All quantities used in this study are ratios of expected richness within an ecoregion (anthropogenic to natural cell, 2024 to 2020) or within-ecoregion comparisons, so *α*_*e*_ cancels and does not enter the results; it is therefore not shared. I alternated fitting by ecoregion with updating the biome effects and variances five times (iterative empirical Bayes). Each taxon has a single *z*, chosen to maximize the sum over all ecoregions of the log marginal posterior density (birds 0.880, mammals 0.954, reptiles 0.943, amphibians 0.966, which gives a cell split equally between two classes about 2–9% more expected richness than the mean of the two weights). Surveys not used in the estimation confirmed that *z* estimated in this way is appropriate (Supplementary Note 9).

#### Intervals and usable weights

I drew 400 sets of values from the Laplace-approximated posterior of each ecoregion and took the 5th to 95th percentiles of the draws as 90% intervals, with *z* fixed at its estimate. In simulated data these intervals tended to be too narrow and to contain the true value too rarely, so I corrected the intervals in the main text by adding the model error measured in the simulated data (Supplementary Note 1). Uncertainty in *z* is examined by recalculating the richness ratios with *z* raised by 0.03, with the weights unchanged (Supplementary Note 1). I flagged two kinds of weight: imprecise weights (posterior standard deviation of the within-ecoregion relative value above 0.5) and extrapolation-dependent weights (those that fell by more than half when the extrapolation of each cell was limited to what would be found with three times its current records).

Only the remaining usable weights (94.7–98.6% of ecoregion–class combinations) were used in subsequent analyses (Supplementary Note 8).

### Composition estimate

#### Species lists standardized to equal sample coverage

Cells with more records have longer lists, so a direct comparison makes well-recorded cells look alike. Rarefaction to an equal number of days leaves lists shorter in cells where daily effort is low, even if the same species are present; standardizing by coverage removes this difference^19^. I therefore standardized the lists so that sample coverage reached the same value *C*\* in every cell. A list is built by adding recording days in random order and stopping when estimated coverage reaches *C*\*, and the probability *p*_*gs*_ that species *s* is on this list for cell *g* was calculated exactly from *T*_*g*_ and *x*_*gs*_ (Supplementary Note 10). The length of the list (the expected number of species) and the expected number of species shared by the lists of two cells *g* and *g*′ in the same ecoregion are

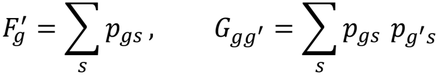

The list length 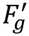 is the number of species found when surveying up to coverage *C*\*, a different quantity from the expected richness *F*_*e*_(*a*) of the richness estimate; *F*′ is used only within the composition estimate, to convert the number of shared species to a share. The estimation used cells with at least five recording days whose coverage using all days reached *C*\*, which I set to 0.547, the level that gives lists of about five days in a typical natural cell (Supplementary Note 11; because the choice also used the three surveys, the evaluation is not fully independent of it).

#### Sharing model

For pairs of cells 5–500 km apart in the same ecoregion, I modelled the number of species on the list of cell *g*′ that are also on the list of cell *g* with a beta-binomial distribution, fitted by a pseudo-likelihood that uses the non-integer expected counts directly (Supplementary Note 3):

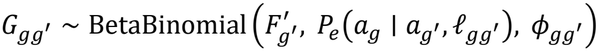

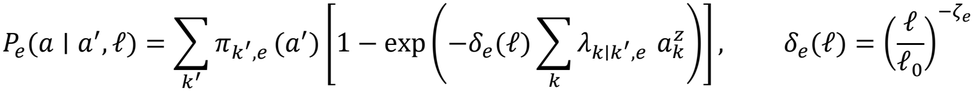

The sharing probability *P*_*e*_(*a* ∣ *a*′, ℓ) is the probability that a species on the list of a cell with mix *a*′ is also on the list of a cell with mix *a* at distance ℓ, and 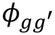 is the precision of the beta-binomial distribution. The origin share 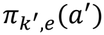 is the share of the species on a list with mix *a*′ that originate in class *k*′; the strength 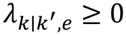 describes how strongly species originating in class *k*′ persist in a cell of destination class *k*; and *δ*_*e*_(ℓ) captures the decline in sharing with distance, with exponent *ζ*_*e*_ and with ℓ_0_ the median distance between pairs of natural cells. All predictions are evaluated at distance ℓ_0_. Supplementary Note 3 explains how the origin shares are built.

#### Shrinkage within biomes and retention values used in the results

To stabilize estimates for poorly recorded classes, I grouped similar origin classes and shrank the strengths *λ* towards the group and biome values (Supplementary Note 3). I drew 400 sets of parameter values from a normal approximation at the posterior mode. Only retention values well supported by data were used in the results: those that are sufficiently precise (posterior standard deviation at most half the value) and for which the data contain at least five pairs of a cell dominated by that class and a cell dominated by natural classes. As for richness, the intervals include the model error measured in simulated data (Supplementary Note 1).

The richness and composition estimates were made in R 4.6.1^49^ with the packages RTMB 2.0^50^, TMB 1.9.25^51^ and statmod 1.5.2^52^.

### Comparison of richness and composition under alternative natural references

To examine how the assessment of anthropogenic land changes with the natural state it is compared against (Fig. 3), I built three versions of a natural reference cell, a hypothetical cell consisting only of natural classes of the ecoregion. The land-cover mix ā_*e*_ of the all-natural reference is the 2020 area of each natural class in the ecoregion, rescaled to sum to 1 across natural classes (if the natural part of an ecoregion is 60% forest and 40% grassland, so is the reference cell). The other two versions restrict ā_*e*_ to forest (1–5) or to open natural classes (6–10) and rescale (ecoregions where that part has no area are excluded). The mosaic is compared as an anthropogenic class and is not used as a reference. Writing the mix consisting only of anthropogenic class *h* as **1**_*h*_, I calculated

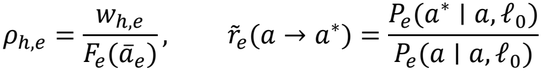

The richness ratio *ρ*_*h,e*_ is the expected richness of a cell consisting only of class *h* divided by that of the natural reference cell. Species retention 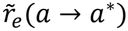 is the share of the species on the list of a cell with mix *a* that are also on the list of a cell with mix *a*\*, divided by the same share between two cells with mix *a*. Even two cells with the same mix do not share all species, because lists are incomplete and species turn over with distance; the denominator cancels this part. A retention of 1 means that a cell with mix *a*\* holds as many of the species of a cell with mix *a* as another cell with mix *a* does, and values above 1 mean that it holds more. Fig. 3 shows the retention 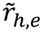 of the natural reference cell’s species in the anthropogenic cell (*a* = ā_*e*_, *a*\* = **1**_*h*_). For example, if 80 of the 100 species on the list of a natural reference cell are on the list of another natural reference cell, and 60 are on the list of an urban cell, retention is 0.75. Extended Data Fig. 3 shows the reverse direction, the retention of the anthropogenic cell’s species in the natural reference cell (*a* = **1**_*h*_, *a*\* = ā_*e*_); the further below 1, the more of the anthropogenic cell’s species are absent from the natural reference. Unless the direction is stated, retention means 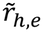.

I calculated these quantities only for combinations of ecoregion and anthropogenic class supported by data for both richness (weights with at least five recorded cells in which the class covers at least half the area) and composition (the retention values used in the results above). So that the references could be compared, I further restricted them to combinations for which all three references could be built. For each of the 400 draws (including model error), I took the median across ecoregions; the point estimate and 90% interval are the median and the 5th and 95th percentiles of these 400 medians.

Differences in retention between the mosaic and cropland were taken per draw in ecoregions with both classes and summarized in the same way.

### Independent bird surveys and field quantities

I used three standardized bird surveys not used in the estimation: the Southern African Bird Atlas Project 2 in southern Africa (SABAP2; squares of about 9 km)^53^, Birdata in Australia (BirdLife Australia; 5 km cells)^54^ and the North American Breeding Bird Survey (BBS; 400 m strips on both sides of routes of about 40 km)^55^ (Supplementary Note 12 gives the survey protocols). Survey units were assigned to the ecoregions of the 5 km cells they contain: SABAP2 squares only if all their cells were in one ecoregion, Birdata cells to their own ecoregion, and BBS routes to the ecoregion containing the largest share of route length, provided that share was at least 80%. SABAP2 records were excluded from the estimation as atlases of five-minute grid cells, Birdata is not in the input because of its licence (CC BY-NC), and the BBS is not in GBIF. The spatial comparison used surveys from 2017–2023, centred on 2020, and only ecoregions with at least five units (Supplementary Table 11).

#### Field richness and composition

Field richness *Y*_*u*_ is the expected number of species when a fixed number of surveys *T*\* (four for SABAP2, ten for Birdata and three for the BBS) is drawn from those of survey unit *u*. Field composition similarity Ō_*u*_ is the mean field overlap *O*_*uv*_ with the set *V*_*u*_ of natural units of the same ecoregion (only units with at least three natural units other than *u* are used), where *O*_*uv*_ is the presence/absence Sørensen similarity of lists of *T*\* surveys drawn from each of the two units, averaged over 50 redraws:

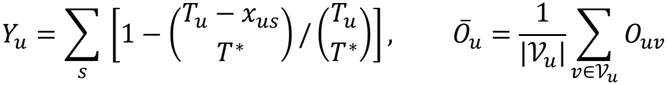

Here *T*_*u*_ is the number of surveys of the unit and *x*_*us*_ the number on which species *s* was recorded. The abundance-based measure replaces Sørensen with Bray–Curtis similarity (Supplementary Note 12, Extended Data Fig. 2).

### Predictors for survey units and existing indicators

For each unit I built the predictors *X*_*u*_ to be compared (the superscript gives the type). The baseline is natural-cover share, 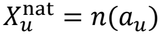, and the richness estimate is 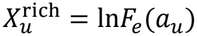 (mean over draws). The composition estimate is the log ratio of the share of species predicted to be shared with natural units to the same share between natural units:

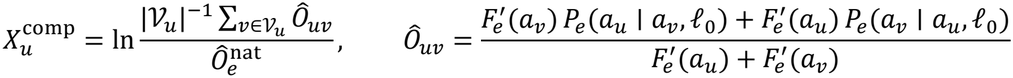

Here 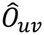 is the Sørensen similarity predicted from the two mixes (mean over draws), and 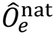 is the mean of 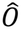 over all pairs of natural units in the ecoregion. 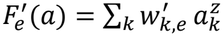 expresses the standardized list length as a function of the mix *a*, with class weights 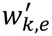 estimated by fitting to the list lengths 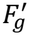 of cells (Supplementary Note 3).

#### Seven existing indicators of land-use impact

MSA (mean species abundance; the GLOBIO 4 model)^6^ is the mean ratio of species abundance to that in an undisturbed state. BHI (Biodiversity Habitat Index)^5^ is the share of the diversity of the original state that remains, weighted by compositional turnover. PDF (potentially disappeared fraction of species)^7^ is the fraction of species lost relative to natural habitat. BII (Biodiversity Intactness Index) is total abundance multiplied by compositional similarity to primary vegetation; I used the versions of Impact Observatory (BII (IO))^4^ and the Natural History Museum, London (BII (NHM))^3^. LIFE (Land-cover change Impacts on Future Extinctions)^8^ is the change in global extinctions per unit area when natural habitat is converted to arable land. STAR (Species Threat Abatement and Restoration)^9^ is the opportunity to reduce threats to threatened and Near Threatened species.

For MSA, BHI and PDF, the coefficients 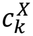 given by the original sources for each land-cover type were multiplied by the mix of the unit and summed (coefficient versions: 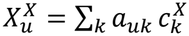, with natural classes set to 1 and the mosaic to the mean of the cropland and natural values; Supplementary Note 2). For the two versions of BII, BHI and LIFE, the 2020 values of the published maps were averaged over the unit with area weights (map versions; Supplementary Table 4).

STAR was rebuilt for birds to match the conditions of the evaluation (2020 land cover; breeding and resident distributions). I built each species’ Area of Habitat (AoH; the part of its range within its habitat and elevation limits)^11^ in 5 km cells by restricting BirdLife ranges (breeding and resident)^56^ with IUCN habitat and elevation limits^57^ and 2020 MODIS land cover. For each species from Near Threatened to Critically Endangered, I multiplied its AoH in each cell by its Red List category weight (from 1 for Near Threatened to 4 for Critically Endangered) and divided by its total AoH. I then summed these values over species. The original STAR multiplies by the proportional contribution of each threat, but these proportions sum to 1 for each species, so STAR across all threats can be calculated without threat information (Supplementary Note 2). Unit values are area-weighted means of cell values over the unit. The correspondence between IUCN habitat types and MODIS classes was built from the table of Lumbierres et al.^58^ (Supplementary Note 2 and Supplementary Data 1).

All predictors were oriented so that larger values mean less modification: natural classes are 1 in all coefficient versions, and the sign of STAR, for which larger values are worse, was reversed (Supplementary Table 4 gives the directions).

#### Richness map

Separately from the indicators of land-use impact, I compared the IUCN AoH richness map for birds (stacked species AoH from Red List 2021-3 on a 5 km Mollweide grid)^10,59^. The map was transferred to the 0.05° grid by nearest neighbour and averaged over each unit with area weights. To separate the part of the map’s information that comes from land cover from the part that comes from location, I regressed the log of the unit values on ecoregion fixed effects and the shares of the 17 classes. The fitted values minus the mean fitted value of the units in the ecoregion form the land-cover part, and the residuals form the location part (range edges, elevation and other effects not explained by the mix). Both parts were evaluated with the same rank correlation as the AoH map (Supplementary Note 2).

### Comparison procedures

#### Rank correlation

How correctly each predictor ranks units within an ecoregion was measured by the Spearman rank correlation *r*_*X,j*_ for each survey *j*. For richness, I took the rank correlation between ln*Y*_*u*_ and the predictor after subtracting ecoregion means from each (field richness is already rarefied to a common number of surveys, so covariates for effort and area enter only the effect-size models). For composition, I took the rank correlation of the residuals of Ō_*u*_ and of the predictor after regressing each on ecoregion fixed effects and covariates (survey effort and the mean log distance to the natural units compared; Supplementary Note 12). Because species sharing declines with distance regardless of land cover, a predictor that simply ranks units close to natural units highly would appear to perform well without capturing any effect of land cover; distance therefore had to be removed. To test whether the coefficient versions carry information beyond natural-cover share, I also measured the within-ecoregion rank correlation between their values and natural-cover share on the survey units of the richness comparison.

#### Effect size

Effect size expresses where the explained variance lies between the baseline and an upper limit:

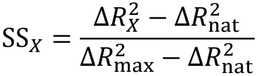

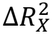 is the increase in the within-ecoregion coefficient of determination when predictor *X* is added to a model with only ecoregion fixed effects and covariates for effort and area, and nat denotes the baseline, natural-cover share. For composition, the covariates are effort and the log distance to natural units (plus the log cosine of latitude for surveys in which unit area varies with latitude), and the field quantity is Ō_*u*_. The upper limit 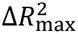 indicates how much a prediction based only on land-cover mix can explain: it is the value of a model with the shares of the 17 classes as separate explanatory variables and coefficients fitted directly to the field data. No predictor built only from the mix is likely to explain more than a model that fits the mix freely to the field data, so I treat this value as a practical upper limit. To prevent overfitting from inflating it, it was evaluated by five-fold cross-validation within ecoregions (Supplementary Note 13). An effect size of 0 means the same as the baseline and 1 means reaching the upper limit. Because 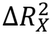 and 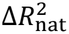 are in-sample values, a single predictor can nevertheless exceed the upper limit, giving an effect size above 1. Where the denominator is small, the value from the full data is unstable (for the composition estimate in North America, 2.48 from the full data against a bootstrap median of 1.79), so effect-size points are bootstrap medians, whereas rank-correlation points are full-data values. Effect size does not distinguish the sign of the relation between predictor and field quantity, which is read from the rank correlation.

To measure what the richness estimate adds to the AoH richness map, I calculated the increase in the within-ecoregion coefficient of determination when the richness estimate is added to a model with ecoregion fixed effects, covariates and the AoH map.

#### Intervals and decision rules

I ran 2,000 bootstrap replicates, resampling ecoregions and using one set of estimation draws per replicate (Supplementary Note 13). I compared the estimate from this study with each predictor through the difference in rank correlation, Δ_*j*_ = *r*_est,*j*_ − *r*_*X,j*_. For richness, the estimate was considered better than a predictor if the 90% interval of Δ_*j*_ was above 0 in at least two of the three surveys. For composition, the 90% interval widths of the difference from natural-cover share differ by more than two-fold among the surveys (0.27 for North America, 0.12 for Australia), so I combined the surveys into one precision-weighted mean and considered the estimate better if the lower end of its 90% interval was above 0 (a one-sided test):

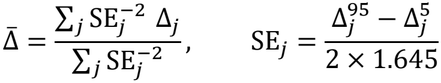

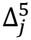 and 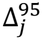 are the 5th and 95th bootstrap percentiles. The main comparisons for composition were with BII (IO) and the BHI map on the presence/absence measure, because they include compositional similarity in their definitions (BII (NHM), which shares the definition of BII (IO), is also shown in Fig. 2b and Extended Data Fig. 1b); the secondary comparison was with BII (IO) on the abundance-based measure. For effect size, I checked whether the 90% interval was above 0 in each survey. Differences between the existing indicators and natural-cover share were not tested formally; I describe them by their rank correlations and effect sizes in each survey (Fig. 2 and Extended Data Fig. 1).

### Three taxa other than birds

Richness and composition of mammals, reptiles and amphibians were estimated with the same coverage level and models, with *z* set to the value from the richness estimate of each taxon. A version that lowered the minimum number of recording days per cell to three moved retention by more than 10%, so I kept the minimum of five days (Supplementary Note 14).

### Global change from 2020 to 2024

This calculation illustrates the application of the estimates to land-cover change, by showing how much global richness and composition are predicted to have changed from 2020 to 2024.

#### Land-cover mixes at two times

Separately from the mixes used in the estimation, I built the 2020 mix *a*_*g*_ and the 2024 mix *b*_*g*_ of cell *g* from those of its 100 pixels (500 m) that kept one class in all five years or changed persistently between 2020–2021 and 2022–2024, excluding pixels that alternated between years (classification noise). Classes with an area share below 0.02 were treated as absent (Supplementary Note 4). Of 6,065,077 land cells, 2,678,963 contained pixels with persistent change (changed cells).

#### Cell-level change and global values

For each cell, I calculated

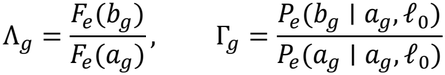

Λ_*g*_ is the ratio of expected richness, and Γ_*g*_ is the compositional change ratio: the share of the species of a community with the 2020 mix predicted to be found in a cell with the 2024 mix, relative to no change. Both are reported as percentages after subtracting 1, and unchanged cells are set to 1. I calculated Λ_*g*_ and Γ_*g*_ only for changed cells in which every class appearing at either time (for composition, also classes appearing in the intermediate mixes that separate transitions among natural classes from those involving anthropogenic classes) was supported by data: for expected richness, a usable weight with at least five recorded cells in which the class covers at least half the area; for composition, the retention values used in the results. Which cells were included was determined solely from the GBIF-based estimates. The share of changed area included was 78%, 56%, 44% and 41% for expected richness in birds, mammals, reptiles and amphibians, and 70%, 38%, 27% and 23% for composition.

Global values are means over unchanged land cells (for composition, those in ecoregions with a fit) and the changed cells included. For expected richness, Λ_*g*_ − 1 was averaged with weights equal to the product of cell area (proportional to the cosine of the central latitude) and 2020 expected richness, which equals the rate of change in the sum over land of expected richness times area. Without weights, the gain or loss of a few species in species-poor cells would count as much as changes in species-rich cells. Because Γ_*g*_ is already the share of a community’s species that remain, it was averaged with area weights alone. When the mixing exponent was fixed at 1 for comparison, I used the same weights (across all cells for which these ratios can be calculated, −0.070% for birds becomes +0.036%; Supplementary Note 4 compares four weighting schemes).

#### Point estimates and intervals

For expected richness, intervals are the 5th to 95th percentiles of the global values across 400 draws of the weights, with model error added and the mixing exponent fixed. For composition, they are the same percentiles across 400 draws of the composition estimate, with model error added to the strengths *λ*. Point estimates are values at the posterior mean of the weights for expected richness and means of the draws for composition. Global values are weighted means of cell values and are therefore summarized by the mean of the draws, unlike the quantities under alternative natural references, which are ecoregion medians of ratios with skewed distributions and are summarized by the median of the draws. Neither interval accounts for land-cover misclassification.

### Use of generative AI

I used Anthropic’s Claude (through Claude Code; models Claude Opus 5 and 5.5, Claude Sonnet 5 and Claude Fable 5.1) to implement analysis code, write plotting code, search the literature, and draft and translate text. I decided the research questions, chose the methods, interpreted the results, drew the conclusions and wrote the manuscript. Code written by the AI was checked by validating the estimates on simulated data with known true values (Supplementary Note 1) and by comparison with independent surveys not used in the estimation (Supplementary Note 12). I reviewed all released code, text and citations, and take responsibility for the content of this paper.

## Data availability

The results presented in the paper (tables of coefficients, estimated maps and source data for the figures) are available from Zenodo^60^ (https://doi.org/10.5281/zenodo.24102473; public on acceptance). The estimates, including the selection of cells that receive values in the global change analysis, use only data that can be used commercially: GBIF occurrence records licensed under CC0 or CC BY 4.0^43–46^, MODIS land cover^47^ and RESOLVE ecoregions^18^. The survey data of SABAP2^53^ and the BBS^55^ are publicly available. Data with non-commercial terms (Birdata^54^ and the BHI and BII (NHM) maps) and data derived from BirdLife ranges^56^ or the IUCN Red List^57^ (LIFE, the rebuilt STAR and the IUCN AoH richness map^59^) were used only for evaluation and comparison; they are available from their providers under their terms (Supplementary Table 4), and survey-unit values derived from them are not distributed.

## Code availability

The code needed to reproduce the results is available from Zenodo^60^ (https://doi.org/10.5281/zenodo.24102473; public on acceptance).

## Acknowledgements

This study was partly supported by the Environment Research and Technology Development Fund of the ERCA (JPMEERF26S21310) funded by the Ministry of the Environment.

## Author contributions

T.K. conceived the study, designed and performed the analyses, interpreted the results and wrote the manuscript.

## Extended Data

**Extended Data Fig. 1.**
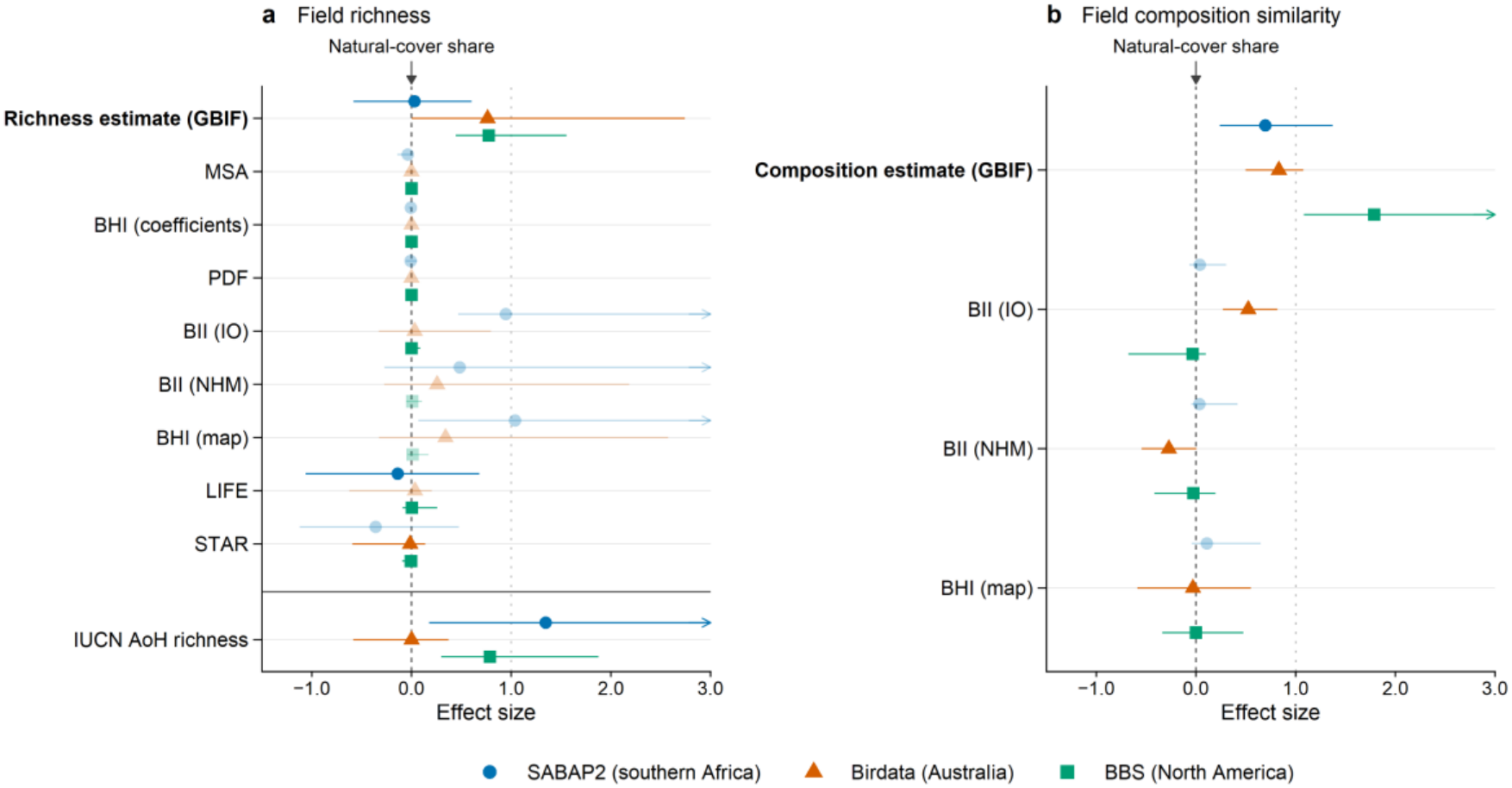
Effect sizes support the rank correlations: in most surveys, the GBIF-based estimates explain much of the gap between natural-cover share and what land-cover mix can explain at best. (a) Field richness. (b) Field composition similarity. An effect size of 0 (dashed line) equals natural-cover share and 1 (dotted line) reaches an upper limit, a model that fits the shares of the 17 land-cover classes to the field data, so values further right are better; values can exceed 1 because that limit is cross-validated (Methods). Pale symbols mark predictors with a negative rank correlation (Fig. 2), which can still have a large effect size. Points are bootstrap medians and lines are 90% intervals; arrowheads mark intervals extending beyond the axis. Rows in (b) are the composition estimate and the map indicators whose definitions include compositional similarity.

**Extended Data Fig. 2.**
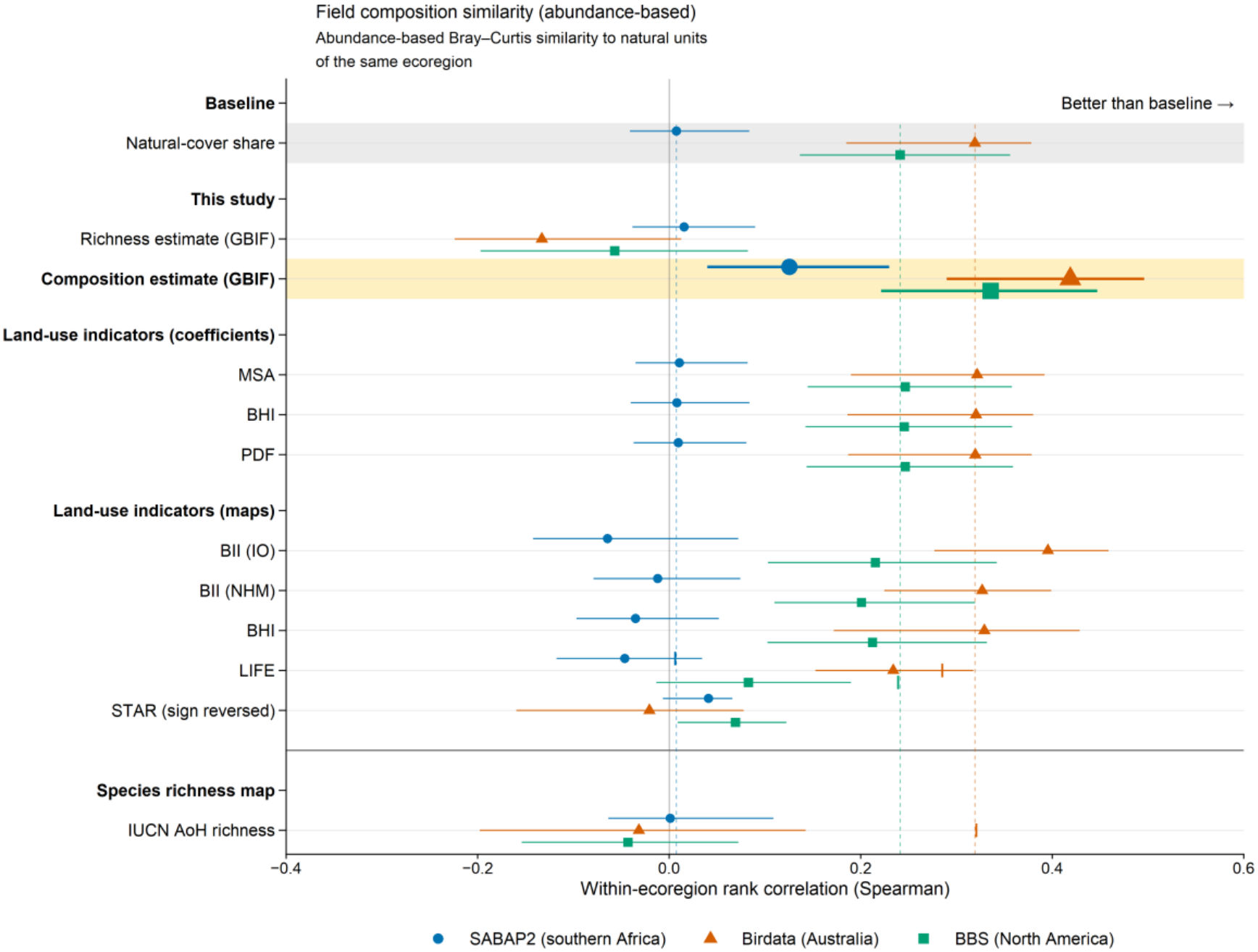
The conclusion for composition holds with an abundance-based measure. As Fig. 2b, but with field composition measured as the mean abundance-based Bray–Curtis similarity to natural units of the same ecoregion. Numbers of survey units are 3,078, 1,379 and 912 (Supplementary Table 11 gives the other rows).

**Extended Data Fig. 3.**
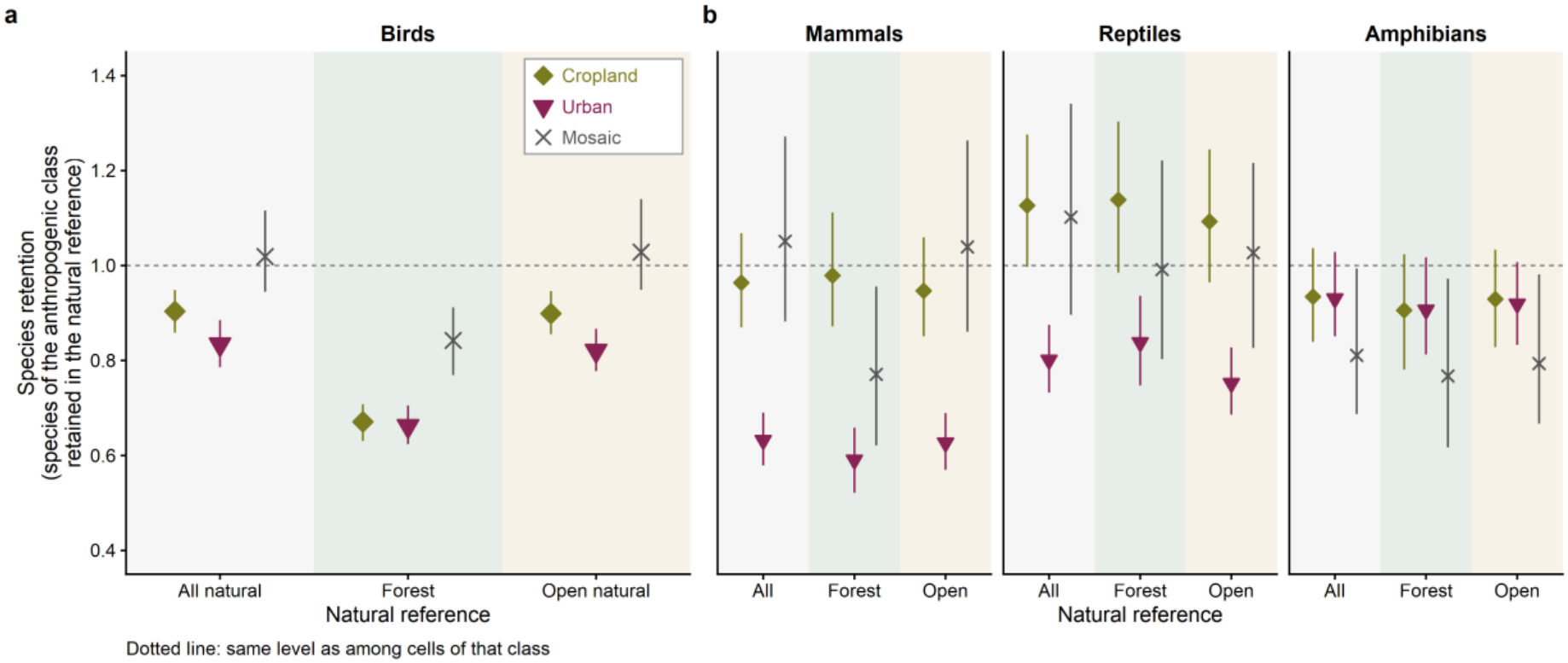
In birds, mammals and reptiles, urban areas tend to hold more species absent from the natural reference than cropland does. (a) Birds and (b) mammals, reptiles and amphibians. The vertical axis is retention in the direction opposite to Fig. 3: how much of the species of a cell of an anthropogenic class is also found in the natural reference cell, relative to the level among cells of that class. Below the dotted line, more of the species of that class are absent from the natural reference. References, symbols, intervals and numbers of ecoregions are as in Fig. 3.

**Extended Data Fig. 4.**
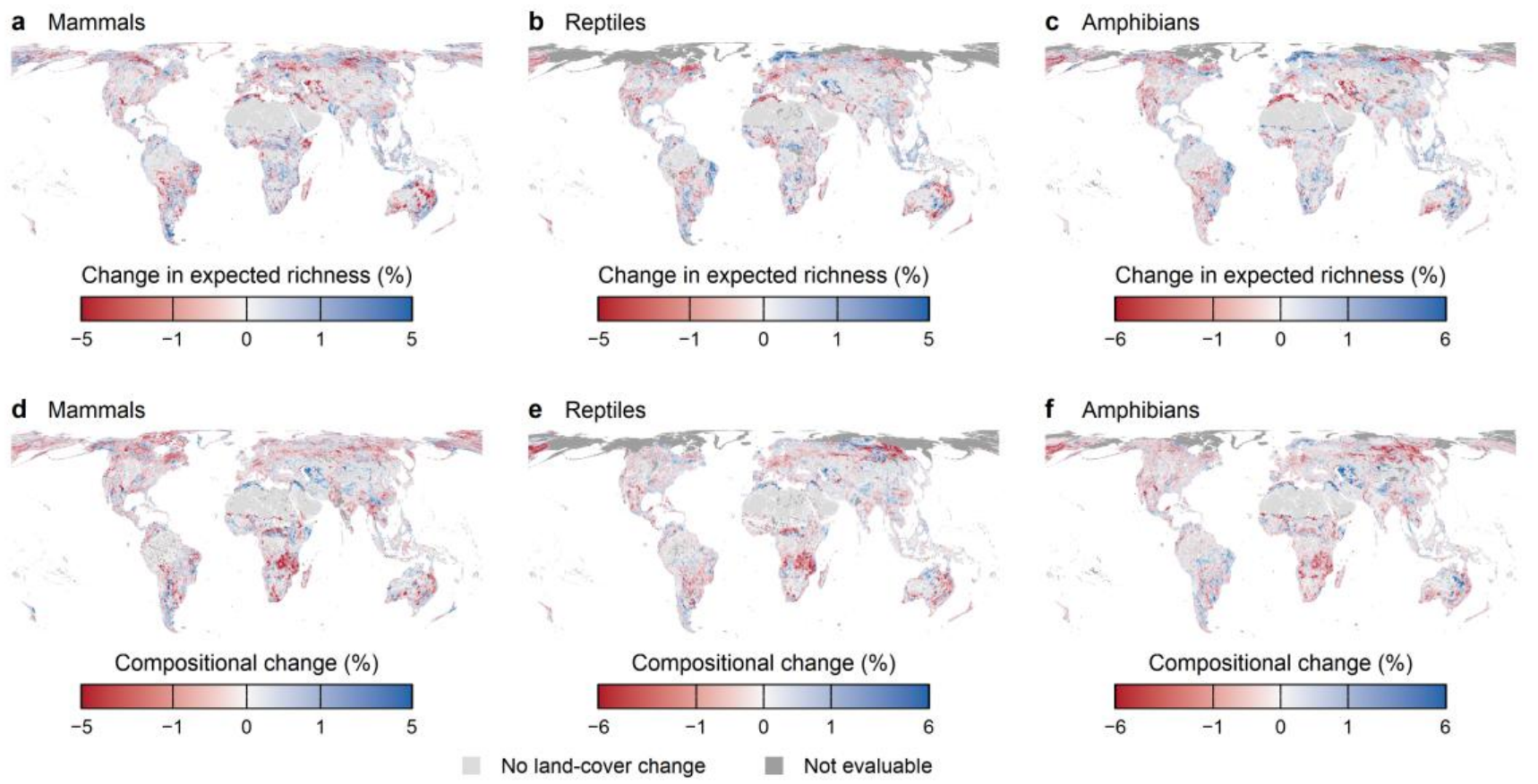
Predicted change from 2020 to 2024 in mammals, reptiles and amphibians. (a)–(c) Change in expected richness and (d)–(f) compositional change, defined and mapped as in Fig. 4a,b, including the pale and dark grey. Colour scales differ among panels. In (d)–(f), cells in ecoregions without a composition fit take the mean value of the same biome (see Supplementary Fig. 3).

## References

1. Affinito, F., Williams, J. M., Campbell, J. E., Londono, M. C. & Gonzalez, A. Progress in developing and operationalizing the Monitoring Framework of the Global Biodiversity Framework. Nat. Ecol. Evol. 8, 2163–2171 (2024).

2. Newbold, T. et al. Has land use pushed terrestrial biodiversity beyond the planetary boundary? A global assessment. Science 353, 288–291 (2016).

3. De Palma, A. et al. Annual changes in the Biodiversity Intactness Index in tropical and subtropical forest biomes, 2001–2012. Sci. Rep. 11, 20249 (2021).

4. Gassert, F., Mazzarello, J. & Hyde, S. Global 100m Projections of Biodiversity Intactness for the Years 2017–2025 (v1.1) (Impact Observatory & Vizzuality, 2026); https://source.coop/vizzuality/biodiversity-intactness-100m-v1-1

5. Hoskins, A. J. et al. BILBI: supporting global biodiversity assessment through high-resolution macroecological modelling. Environ. Model. Softw. 132, 104806 (2020).

6. Schipper, A. M. et al. Projecting terrestrial biodiversity intactness with GLOBIO 4. Glob. Change Biol. 26, 760–771 (2020).

7. Scherer, L. et al. Biodiversity impact assessment considering land use intensities and fragmentation. Environ. Sci. Technol. 57, 19612–19623 (2023).

8. Eyres, A. et al. LIFE: a metric for mapping the impact of land-cover change on global extinctions. Phil. Trans. R. Soc. B 380, 20230327 (2025).

9. Mair, L. et al. A metric for spatially explicit contributions to science-based species targets. Nat. Ecol. Evol. 5, 836–844 (2021).

10. Lumbierres, M. et al. Area of Habitat maps for the world’s terrestrial birds and mammals. Sci. Data 9, 749 (2022).

11. Brooks, T. M. et al. Measuring terrestrial Area of Habitat (AOH) and its utility for the IUCN Red List. Trends Ecol. Evol. 34, 977–986 (2019).

12. Dornelas, M. et al. Assemblage time series reveal biodiversity change but not systematic loss. Science 344, 296–299 (2014).

13. Blowes, S. A. et al. The geography of biodiversity change in marine and terrestrial assemblages. Science 366, 339–345 (2019).

14. Newbold, T. et al. Global effects of land use on local terrestrial biodiversity. Nature 520, 45–50 (2015).

15. Gonzalez, A. et al. Estimating local biodiversity change: a critique of papers claiming no net loss of local diversity. Ecology 97, 1949–1960 (2016).

16. Hillebrand, H. et al. Biodiversity change is uncoupled from species richness trends: consequences for conservation and monitoring. J. Appl. Ecol. 55, 169–184 (2018).

17. Nature Positive Initiative. State of Nature Indicators and Metrics Framework (Nature Positive Initiative, 2026); https://www.naturepositive.org/app/uploads/2026/09/State-of-Nature-Indicators-and-Metrics-Framework.pdf

18. Dinerstein, E. et al. An ecoregion-based approach to protecting half the terrestrial realm. BioScience 67, 534–545 (2017).

19. Chao, A. & Jost, L. Coverage-based rarefaction and extrapolation: standardizing samples by completeness rather than size. Ecology 93, 2533–2547 (2012).

20. Ding, C. et al. Land use and climate warming interact to reshape global avian biodiversity through ecological filtering and homogenisation. Preprint at 10.64898/2026.10.08.757631 (2026).

21. Chaudhary, A. & Brooks, T. M. Land use intensity-specific global characterization factors to assess product biodiversity footprints. Environ. Sci. Technol. 52, 5094–5104 (2018).

22. Sanchez-Ortiz, K. et al. Overcoming limitations in selection guidelines for state-of-nature metrics in corporate assessments. Preprint at 10.32942/x2vq3r (2026).

23. Durand, M. et al. Comparison of three biodiversity metrics to evaluate corporate no net loss achievement under spatial constraints. J. Environ. Manage. 396, 128158 (2025).

24. Blowes, S. A. et al. Local biodiversity change reflects interactions among changing abundance, evenness, and richness. Ecology 103, e3820 (2022).

25. Pereira, H. M. et al. Essential biodiversity variables. Science 339, 277–278 (2013).

26. Kissling, W. D. et al. Building essential biodiversity variables (EBVs) of species distribution and abundance at a global scale. Biol. Rev. 93, 600–625 (2018).

27. Veldman, J. W. et al. Where tree planting and forest expansion are bad for biodiversity and ecosystem services. BioScience 65, 1011–1018 (2015).

28. Bueno, A. S. & Peres, C. A. The role of baseline suitability in assessing the impacts of land-use change on biodiversity. Biol. Conserv. 243, 108396 (2020).

29. Ellis, E. C. et al. People have shaped most of terrestrial nature for at least 12,000 years. Proc. Natl Acad. Sci. USA 118, e2023483118 (2021).

30. Kadoya, T. & Washitani, I. The Satoyama Index: a biodiversity indicator for agricultural landscapes. Agric. Ecosyst. Environ. 140, 20–26 (2011).

31. Daily, G. C., Ehrlich, P. R. & Sánchez-Azofeifa, G. A. Countryside biogeography: use of human-dominated habitats by the avifauna of southern Costa Rica. Ecol. Appl. 11, 1–13 (2001).

32. Mendenhall, C. D., Karp, D. S., Meyer, C. F. J., Hadly, E. A. & Daily, G. C. Predicting biodiversity change and averting collapse in agricultural landscapes. Nature 509, 213–217 (2014).

33. Estrada-Carmona, N., Sánchez, A. C., Remans, R. & Jones, S. K. Complex agricultural landscapes host more biodiversity than simple ones: a global meta-analysis. Proc. Natl Acad. Sci. USA 119, e2203385119 (2022).

34. Kuipers, K. J. J. et al. Land use diversification may mitigate on-site land use impacts on mammal populations and assemblages. Glob. Change Biol. 29, 6234–6247 (2023).

35. Katoh, K., Sakai, S. & Takahashi, T. Factors maintaining species diversity in satoyama, a traditional agricultural landscape of Japan. Biol. Conserv. 142, 1930–1936 (2009).

36. Phalan, B., Onial, M., Balmford, A. & Green, R. E. Reconciling food production and biodiversity conservation: land sharing and land sparing compared. Science 333, 1289–1291 (2011).

37. Gibson, L. et al. Primary forests are irreplaceable for sustaining tropical biodiversity. Nature 478, 378–381 (2011).

38. Chandler, M. et al. Contribution of citizen science towards international biodiversity monitoring. Biol. Conserv. 213, 280–294 (2017).

39. Heberling, J. M., Miller, J. T., Noesgaard, D., Weingart, S. B. & Schigel, D. Data integration enables global biodiversity synthesis. Proc. Natl Acad. Sci. USA 118, e2018093118 (2021).

40. Callaghan, C. T., Borda-de-Água, L., van Klink, R., Rozzi, R. & Pereira, H. M. Unveiling global species abundance distributions. Nat. Ecol. Evol. 7, 1600–1609 (2023).

41. Kelling, S. et al. Using semistructured surveys to improve citizen science data for monitoring biodiversity. BioScience 69, 170–179 (2019).

42. Gonzalez, A. et al. A global biodiversity observing system to unite monitoring and guide action. Nat. Ecol. Evol. 7, 1947–1952 (2023).

43. GBIF.org. GBIF occurrence download (birds; CC0 and CC BY 4.0). GBIF 10.15468/dl.pwdmqt (2026).

44. GBIF.org. GBIF occurrence download (mammals; CC0 and CC BY 4.0). GBIF 10.15468/dl.jgsvdv (2026).

45. GBIF.org. GBIF occurrence download (reptiles; CC0 and CC BY 4.0). GBIF 10.15468/dl.9sqgwd (2026).

46. GBIF.org. GBIF occurrence download (amphibians; CC0 and CC BY 4.0). GBIF 10.15468/dl.s272tx (2026).

47. Friedl, M. & Sulla-Menashe, D. MODIS/Terra+Aqua Land Cover Type Yearly L3 Global 500m SIN Grid V061 (NASA EOSDIS Land Processes DAAC, 2022); 10.5067/MODIS/MCD12Q1.061

48. Pereira, H. M. & Daily, G. C. Modeling biodiversity dynamics in countryside landscapes. Ecology 87, 1877–1885 (2006).

49. R Core Team. R: A Language and Environment for Statistical Computing, Version 4.6.1 (R Foundation for Statistical Computing, 2026); https://www.R-project.org/

50. Kristensen, K. RTMB: ‘R’ bindings for ‘TMB’. R package version 2.0 10.32614/CRAN.package.RTMB (2026).

51. Kristensen, K., Nielsen, A., Berg, C. W., Skaug, H. & Bell, B. M. TMB: automatic differentiation and Laplace approximation. J. Stat. Softw. 70, 1–21 (2016).

52. Smyth, G. et al. statmod: statistical modeling. R package version 1.5.2 10.32614/CRAN.package.statmod (2026).

53. FitzPatrick Institute of African Ornithology. Southern African Bird Atlas Project 2. Africa Bird Atlas Project https://api.birdmap.africa (University of Cape Town).

54. GBIF.org. GBIF occurrence download (Birdata of BirdLife Australia; 2 ha, 20-minute surveys). GBIF 10.15468/dl.2qhj7a (2026).

55. Ziolkowski, D., Lutmerding, M., Skalos, S. M., English, W. B. & Hudson, M.-A. R. 2026 release – North American Breeding Bird Survey dataset (1966–2025). U.S. Geological Survey 10.5066/P144YU3S (2026).

56. BirdLife International & Handbook of the Birds of the World. Bird Species Distribution Maps of the World, Version 2024.2 (BirdLife International, 2024); https://datazone.birdlife.org/contact-us/request-our-data

57. IUCN. The IUCN Red List of Threatened Species, Version 2024-2 (IUCN, 2024); https://www.iucnredlist.org

58. Lumbierres, M. et al. Translating habitat class to land cover to map area of habitat of terrestrial vertebrates. Conserv. Biol. 36, e13851 (2022).

59. IUCN. Species Richness from AoH Rasters, Birds (Based on IUCN Red List Version 2021-3) (IUCN, 2023); https://www.iucnredlist.org/resources/spatial-data/other-spatial-downloads

60. Kadoya, T. Code and data for: Global biodiversity indicators do not fully capture how land cover alters species richness and composition (v1.0). Zenodo 10.5281/zenodo.24102473 (2026).

