## Supplementary Information for "Most biodiversity indicators add little beyond natural land cover"

**Contents**

- Supplementary Note 1. Validation of the estimates on simulated data (Supplementary Tables 1, 2; Supplementary Figs. 1, 2)
- Supplementary Note 2. Values of the existing indicators and the AoH richness map (Supplementary Tables 3, 4)
- Supplementary Note 3. The composition estimate (a model of the probability that the lists of two cells share a species) and comparison of richness and composition under alternative natural references (Supplementary Table 5)
- Supplementary Note 4. Details of global change from 2020 to 2024 (Supplementary Tables 6, 7; Supplementary Fig. 3)
- Supplementary Note 5. Retrieval of GBIF records and inclusion criteria (Supplementary Tables 8, 9)
- Supplementary Note 6. Check for exotic species
- Supplementary Note 7. Preparation of land cover and ecoregions
- Supplementary Note 8. Details of the richness estimate
- Supplementary Note 9. Validity of the mixing exponent and scale of predicted richness contrasts (Supplementary Fig. 4)
- Supplementary Note 10. Lists standardized to equal sample coverage (Supplementary Table 10)
- Supplementary Note 11. Choice of the sample-coverage level
- Supplementary Note 12. Independent bird surveys and field quantities (Supplementary Table 11)
- Supplementary Note 13. Details of intervals and effect sizes
- Supplementary Note 14. Fitting the three taxa other than birds
- Supplementary Data 1. Correspondence between IUCN habitat types and MODIS land-cover classes (CSV, built following Lumbierres et al.[1]; Supplementary Note 2)
- Supplementary references

### Supplementary Note 1. Validation of the estimates on simulated data

From simulated communities with known properties, I generated records in which recording effort varies among places as it does in GBIF, applied the same procedure as to the real data and compared the estimates with the true values. The composition estimate absorbs differences in recording effort and broadly recovers the ecoregion median of species retention. The richness estimate, by contrast, understates the richness ratios of anthropogenic classes, so model error was added to its intervals.

#### Building the simulated data

**Simulated ecoregions.** Each simulated region (hereafter simulated ecoregion) has 60 × 60 cells of 5 km (300 km square) and 300–500 species. There are 88 simulated ecoregions of three types that mimic the size, number and survey frequency of the units of the three bird surveys (41 North American, 23 Australian and 24 southern African). Species belong to four types: specialists of forest, woodland, open vegetation, waterside and similar habitats; natural generalists; species tolerant of human environments; and newly arrived species. In urban areas natural species decline and species tolerant of human environments increase.

**Landscapes.** Of the 17 land-cover classes, area was placed in 10 natural classes (1, 2, 4, 7, 8, 9, 10, 11, 16 and 17) and the three anthropogenic classes. Landscapes were not copied from the land-cover mixes of the three bird surveys but generated from spatially smooth random fields. In the natural part, smooth fields set the shares of five groups (forest, woodland and shrubland, open vegetation, waterside and barren land), and further fields set the shares of classes within groups. Anthropogenic land was placed only where a separate smooth field of "intensity of human use" was high, divided into cropland and mosaic, with urban areas placed as towns centred on those parts. Because the recovery of the richness estimate can depend on the landscape, I built six sets of simulated data (hereafter six worlds) that share the way species, populations and records are generated and differ only in how the landscape is built (Supplementary Table 1). The six differ in the grain of the fields (patch size), the purity of classes within patches and the amount of anthropogenic land. One world was matched to real data, not to the sites of the three surveys but to the land-cover mix of all real cells with bird records (the share of cells in which one class covers at least 90% of the area and the area share of anthropogenic classes). Supplementary Figs. 1 and 2 show the base world.

**Supplementary Table 1 Six worlds differing in how the landscape is built.** Pure cells: share of cells in which one class covers at least 90% of the area. Anthropogenic share: mean area share of anthropogenic classes in a cell. Both are for all cells of the simulated ecoregions.

| World | How the landscape is built | Pure cells | Anthropogenic share |
|---|---|---|---|
| Finely mixed | Small patches, classes grading into each other within patches, and much of the anthropogenic land as mosaic | 0.02 | 0.44 |
| High anthropogenic | More anthropogenic land and more towns than in the base world | 0.09 | 0.70 |
| Base | Intermediate patch size and amount of anthropogenic land | 0.06 | 0.48 |
| Low anthropogenic | Less anthropogenic land and fewer towns than in the base world | 0.03 | 0.09 |
| Coarse and pure | Large patches, nearly one class within each patch and little mosaic | 0.36 | 0.42 |
| Matched to real data | Patches of the coarse and pure world, with anthropogenic land reduced to match real cells with records | 0.45 | 0.16 |

In real cells with bird records, pure cells are 0.34 and the anthropogenic share is 0.21; in cells with records in the world matched to real data, 0.44 and 0.19.

**Ranges.** Each species occupies a range within the simulated ecoregion: the cells in which a spatially smooth random field (Gaussian kernel with a width of 30, 60 or 120 km, chosen for each species) is above the quantile that leaves a share $p_s$ of the ecoregion inside the range. The median of $p_s$ is 0.6, and common species (those with high abundance densities) have wider ranges. Because of the ranges, cells farther apart are less likely to share species.

**Populations.** Within its range, the abundance density of a species differs among classes. Specialists are dense in the classes of their own group, and species differ in which class of the group they prefer. Natural generalists are spread over the natural classes. Species tolerant of human environments are dense in cropland and urban areas and sparse in natural classes. Newly arrived species occur only in anthropogenic classes. The density in the mosaic is the mean of the density in cropland and the density in the mean natural composition of the ecoregion. The abundance $N_{gs}$ of species $s$ in cell $g$ follows a Poisson distribution with a mean determined by the land-cover mix of the cell:

$$N_{gs} \sim \text{Poisson}\left(\iota_{gs} \sum_{k} a_{gk}\ N^{\circ}_{sk}\right)$$

Here $\iota_{gs}$ is 1 if cell $g$ is in the range of species $s$ and 0 otherwise, $a_{gk}$ is the area share of class $k$ in cell $g$ and $N^{\circ}_{sk}$ is the abundance density of species $s$ in class $k$.

**Records.** Urban and cropland cells received more recording days. On a recording day, species $s$ present in cell $g$ is recorded with probability $1 - \exp(-\nu_{gs})$, with recording rate (the same quantity as the recording rate of the observation model in Supplementary Note 8)

$$\nu_{gs} = \theta_g \, \nu^{\circ}_s \left(N_{gs}/\bar{N}_s\right)^{0.5}$$

Here $\theta_g$ is the daily recording effort of the cell (log-normal), $\nu^{\circ}_s$ is the detectability of the species and $\bar{N}_s$ is the mean abundance of species $s$ in cells where it occurs. The power of 0.5 represents the difficulty of recording a few occasional individuals (abundance-dependent detection)[2].

**Half-effort records.** From the same populations I also generated records with daily effort reduced to $\theta_g \cdot 0.5^{1-n(a_g)}$, where $n(a_g)$ is the area share of natural classes in the cell, so effort is halved in purely anthropogenic cells and unchanged in purely natural cells. This shrinks the cropland-to-natural ratio of ten-day list lengths to about 0.7 times that of equal-effort records, which is close to the value in real data for southern Africa (0.74, the GBIF ten-day cropland-to-natural ratio of 0.76 divided by the same ratio of field richness, 1.03).

**Surveys and two time points.** From the same populations I also generated the results of the three survey types and a second time point with altered land cover in some cells (four scenarios converting 30% or 70% of natural cells to cropland or urban, and two scenarios of exchange among natural classes).

**True values.** True retention was calculated by the same formula as in Supplementary Note 3 from the expected sharing of lists whose number of days reaches the target level of true sample coverage, based on the daily recording probability of each species in each cell. The true richness ratio is the true expected richness of a cell consisting only of an anthropogenic class divided by that of the natural reference cell (a hypothetical cell whose natural classes occur in the same proportions as in the natural part of the ecoregion). True expected richness is the sum over species of the probability that the species is in the cell (abundance of at least 1), weighted by the probability that it is in the ecoregion species pool; it was calculated without using the richness function $F_e(a)$ of the estimate.

### The composition estimate absorbs differences in recording effort

With coverage-standardized lists, halving daily recording effort on anthropogenic land barely changed retention on cropland and in urban areas (0.96–0.97 times the equal-effort value; Supplementary Fig. 1a). With rarefaction to the same number of days in each cell, the same halving reduced retention to 0.74–0.77 times the equal-effort value, so natural species appeared to be lost simply because effort on anthropogenic land was low (Supplementary Fig. 1b). In the base world, the median ratio of estimated to true retention was 1.04 for cropland, 1.05 for urban and 0.99 for the mosaic with equal-effort records (0.97–1.01 with half-effort records), so the estimates broadly recovered the true values (Supplementary Fig. 1c). The differences between estimated and true ecoregion medians were −0.03 to +0.04 for the three anthropogenic classes with equal-effort and half-effort records. The estimates of compositional change between two time points recovered the true value determined by the change in land-cover mix alone (the target of the estimate), with median ratios of 0.97–0.99 (0.96–0.99 with half-effort records; Supplementary Fig. 1d). Against the true value in which populations on unchanged land carry over directly, the median ratios were slightly higher, 1.02–1.07 (Supplementary Fig. 1e). The precision of retention per ecoregion was high in landscapes with finely mixed classes and lower in landscapes with many pure cells (rank correlation with true values of 0.91 in the finely mixed world and 0.37 in the world matched to real data; in the latter, the standard deviation of the log difference between estimated and true retention was 0.32).

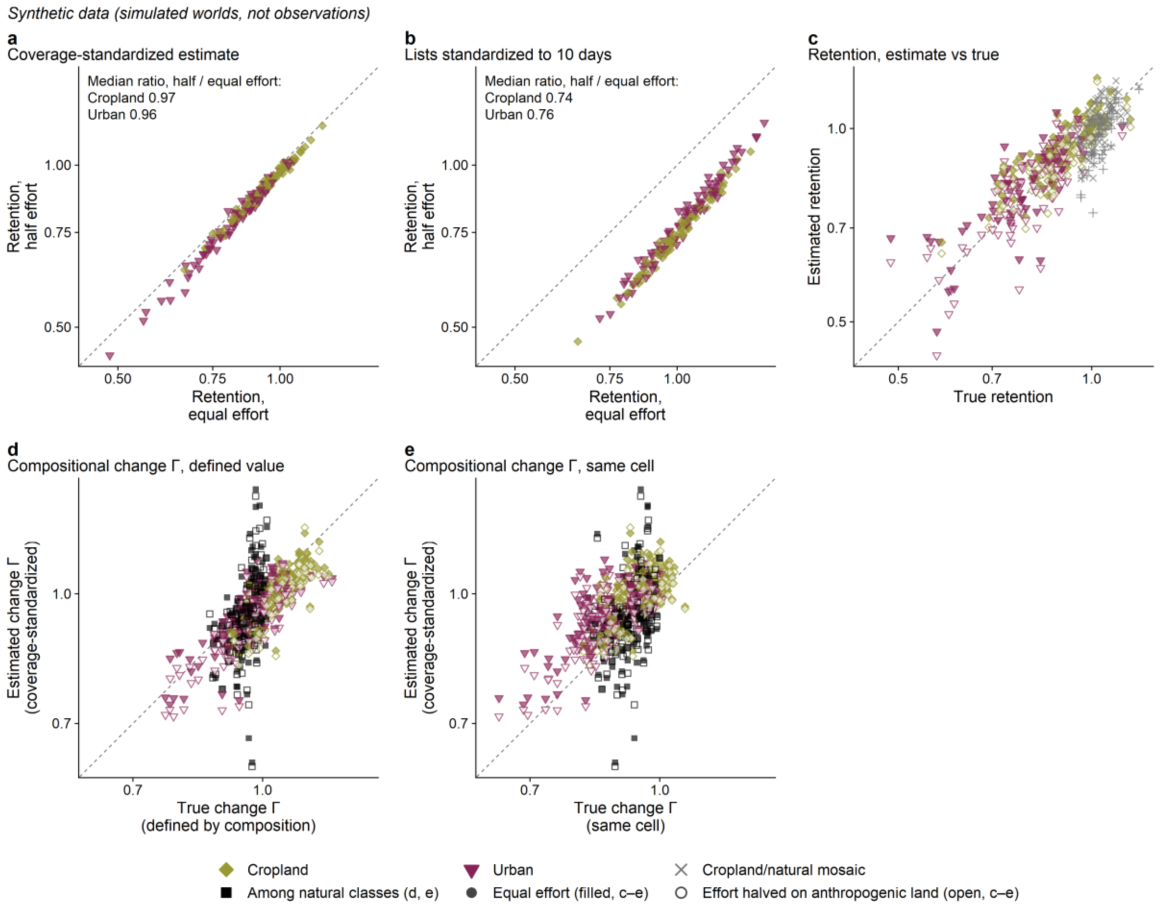


**Supplementary Fig. 1. Simulated data: estimates standardized by sample coverage absorb differences in recording effort.** GBIF-like records were generated from simulated communities with known properties (88 simulated ecoregions); these are not real data. Recording effort is the daily recording effort of a cell (equivalent to the number of observers or the list length on that day), and in the "half" records it was halved on anthropogenic land only. (a), (b) Retention on cropland (olive diamonds) and in urban areas (wine inverted triangles) of each ecoregion, compared between equal-effort records (horizontal axis) and half-effort records (vertical axis). (a) The coverage-standardized estimate (this study). (b) The estimate from lists standardized to ten days in each cell. Points on the 1:1 line mean that differences in effort are absorbed; points below it mean that retention looks lower simply because effort is low. (c) Estimated (vertical axis) and true (horizontal axis) retention. Filled symbols are equal-effort records, open symbols records with effort halved on anthropogenic land (× and + for the mosaic). (d), (e) Estimated (vertical axis) and true (horizontal axis) compositional change between two time points (the ratio of the share of the species of the 2020 community that remain at the second time point; the same quantity as in Fig. 4b). In (d) the true value is determined by the change in land-cover mix alone (the target of the estimate); in (e) it assumes that the same populations of the same cell remain. Symbols are scenarios (conversion to cropland, conversion to urban, change among natural classes). In (c)–(e), points on the 1:1 line mean that the estimate recovers the true value.

### The richness estimate understates the richness ratios of anthropogenic classes

The richness estimate broadly recovered the ratios of class weights but understated the richness ratios of anthropogenic classes, to a degree that depended on the landscape.

In the base world, the within-ecoregion relative class weights had a rank correlation of 0.76 with the true values. The richness ratios of anthropogenic classes were a median 4% below the true values, with a rank correlation of 0.74 (Supplementary Fig. 2). The estimated mixing exponent of 0.876 was almost the same as the true value of 0.878 (the true value is the median across simulated ecoregions of the exponent obtained by fitting the richness function $F_e(a)$ to the true expected richness of cells with less than 5% of their area in anthropogenic classes; the same definition is used in Supplementary Table 2 and under Robustness in the real data below). In all six worlds the estimated richness ratios were below the true values. The bias was small in landscapes with many mixed cells and large in landscapes with many pure cells or little anthropogenic land (Supplementary Table 2).

The main cause lies in the form of the richness function $F_e(a)$. Fitting it directly to true expected richness, without records, produced a bias in the same direction. The reason is that mixed cells constrain mainly the richness that a class adds to a mixture, whereas the target is the richness of a cell consisting only of that class. Because the function adds class contributions, it cannot represent species shared among classes, and so it understates the richness of a single-class cell when few cells are dominated by that class. Generalizing the function with a separate mixing exponent for mixtures of anthropogenic classes reduced the bias by 20–30%. That generalization requires many records from cells that mix anthropogenic classes and is difficult to estimate with real data, so I did not use it.

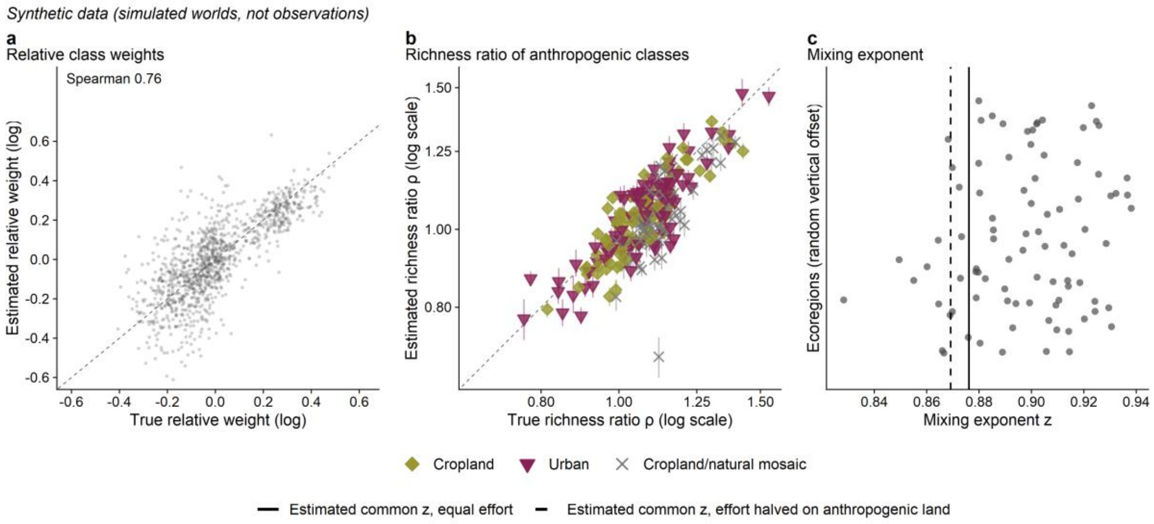


**Supplementary Fig. 2. Simulated data: the richness estimate broadly recovers the ratios of class weights and the richness ratios.** The richness estimate used for the real data was applied to GBIF-like records (equal effort) generated from simulated communities with known properties (base world, 88 ecoregions); these are not real data. (a) Within-ecoregion relative class weights (log difference from the mean of each ecoregion). The horizontal axis is the true value, the vertical axis the estimate, and points are combinations of ecoregion and class (1,138). The dashed line is 1:1. (b) Richness ratios of anthropogenic classes (expected richness of a cell consisting only of that class ÷ expected richness of the natural reference cell, a hypothetical cell whose natural classes occur in the same proportions as in the natural part of the ecoregion). The horizontal axis is the true value (calculated from the presence probability of each species in the simulated world, without using the richness function), the vertical axis the estimate (both on log scales), and vertical lines are 90% intervals of the estimate (before model error is added). Points are combinations of ecoregion and anthropogenic class (264): olive diamonds for cropland, wine inverted triangles for urban and grey × for cropland/natural mosaic. Points below the dashed line mean that the estimate is below the true value. (c) Mixing exponent. Points are the true values of the 88 ecoregions (vertical positions are random jitter), the solid line is the common estimate from equal-effort records and the dashed line the estimate from records with daily effort halved on anthropogenic land.

**Supplementary Table 2 Recovery of the richness estimate in the six worlds.** Summary over three anthropogenic classes × 88 simulated ecoregions (264 combinations). The landscapes of the worlds are in Supplementary Table 1. The true mixing exponent is the median of the exponent obtained by fitting the richness function to the true expected richness of cells with less than 5% of their area in anthropogenic classes. Bias is the median log ratio of estimated to true richness ratio (negative means understated). Coverage of intervals is the share of combinations whose 90% interval contains the true value; the right-hand column is for intervals with model error (standard deviation 0.12) added to the log weights of anthropogenic classes (see Model error added to intervals below).

| World | Estimated mixing exponent (true value) | Bias | Rank correlation with true value | Interval coverage | Same, with model error |
|---|---|---|---|---|---|
| Finely mixed | 0.906 (0.885) | −0.023 | 0.88 | 37% | 100% |
| High anthropogenic | 0.897 (0.890) | −0.018 | 0.77 | 30% | 100% |
| Base | 0.876 (0.878) | −0.045 | 0.74 | 24% | 99% |
| Low anthropogenic | 0.852 (0.891) | −0.115 | 0.63 | 17% | 81% |
| Coarse and pure | 0.842 (0.877) | −0.103 | 0.50 | 11% | 80% |
| Matched to real data | 0.842 (0.873) | −0.140 | 0.60 | 10% | 73% |

### Model error added to intervals

The 90% intervals of the estimates represent only the estimation uncertainty within each ecoregion and contained the true value too rarely (Supplementary Table 2). I therefore widened the intervals for the real data by adding model error, with standard deviations equal to the per-ecoregion differences between estimated and true values in the world matched to real data: 0.12 (anthropogenic classes) and 0.08 (natural classes) on log richness weights and 0.32 on log retention. The errors do not correct bias and were applied independently per ecoregion, class and draw. For global compositional change, errors of the same size were added to the draws of the log persistence strengths of the sharing model (Supplementary Note 3) and propagated through the model equations to the amount of change. With these additions, the intervals of richness ratios contained the true value in 73–100% of cases across the six worlds. By class in the world matched to real data, coverage was 86% for cropland and 93% for urban but only 40% for cropland/natural mosaic. In the simulated worlds the mosaic is a mixture of cropland and natural with no species of its own, and cells dominated by mosaic are almost absent, so the estimation error for the mosaic exceeds twice the added model error and coverage remains low.

### Robustness in the real data

**Extrapolation to pure cells (one class covering at least 90%).** Real data have many more cells dominated by a class than the simulated worlds (108,730 cells with records are at least 90% cropland, against 2,921 in the world matched to real data). Ecoregions with at least five cells in which the class covers at least half the area made up 90% of ecoregions with estimates for cropland, 64% for urban and 81% for cropland/natural mosaic. Richness ratios from weights built from the records of cells dominated by the class, without the richness function, were higher than those from the fitted weights by 0.027 for cropland, 0.028 for urban and 0.025 for the mosaic, less than in the same comparison in the simulated world (about 0.07 for cropland). The reduction of the richness ratios of anthropogenic classes through extrapolation in the real data is estimated at about 0.02–0.03.

**Number of recording days.** In real bird data, cells with more recording days recorded more species per day within an ecoregion. Refitting a model that accounts for this pattern (a model that allows daily recording effort to increase with the number of recording days) lowered the richness ratio of birds in urban areas from 0.846 to 0.790. Cropland (0.813 to 0.823), the mosaic (0.632 to 0.640) and global change in expected richness (−0.070% to −0.067%) barely moved. In the simulated worlds, accounting for this pattern neither improved nor worsened the difference in richness ratios, so the values in the main text come from the model without it (all values compared are from all usable weights, before model error is added).

**Mixing exponent.** In the simulated worlds, the estimated mixing exponent was 0.03–0.04 below the true value in landscapes with many pure cells or little anthropogenic land (Supplementary Table 2). Recalculating the real data with the exponent 0.03 above its estimate (weights unchanged) raised the richness ratios of birds against all natural classes (with model error added, for combinations supported by data for both richness and composition) from 0.81 to 0.84 for cropland, 0.83 to 0.86 for urban and

0.63 to 0.66 for the mosaic (over all ecoregions with values for each class, so the starting values differ slightly from those in Fig. 3, which uses ecoregions with values under all three references). An underestimated exponent therefore gives lower richness ratios.

#### Conclusions

The composition estimate standardized by sample coverage does not generate apparent losses from low recording effort on anthropogenic land and broadly recovers the ecoregion median of retention (precision per ecoregion declines in landscapes with many pure cells). The richness estimate understates the richness ratios of anthropogenic classes, and model error measured in simulated data was added to the intervals. Extrapolation lowers the ratios by an estimated 0.02–0.03 in the real data, and any underestimate of the mixing exponent would lower them further, so the ratios may overstate the loss of richness on anthropogenic land.

### Supplementary Note 2. Values of the existing indicators and the AoH richness map

This note shows how the values of the seven existing indicators and the IUCN AoH richness map were built for each survey unit (coefficients, map versions and the rebuilt STAR) and the results of decomposing the AoH map and STAR. In rank correlations with field richness, the part of the AoH map not explained by the land-cover mix (the location part, such as range edges and elevation) ranked differences in field richness better than the land-cover part. STAR's mismatch with richness stems not from the quality of the rebuilt AoH but from its restriction to threatened and Near Threatened species.

**Seven indicators.** MSA (mean species abundance, the mean ratio of species abundance to that in an undisturbed state; the GLOBIO 4 model)[3]; BHI (Biodiversity Habitat Index, the share of the diversity of the original state that remains, weighted by compositional turnover)[4]; PDF (potentially disappeared fraction of species, the fraction of species regionally lost relative to nature)[5]; BII (Biodiversity Intactness Index, total abundance multiplied by compositional similarity to primary vegetation; the two versions BII (IO) of Impact Observatory[6] and BII (NHM) of the Natural History Museum, London[7]); LIFE (the change in global extinctions per unit area when natural habitat is converted to arable land)[8]; and STAR (Species Threat Abatement and Restoration, the opportunity to reduce threats to threatened and Near Threatened species)[9]. Because BII has two versions, there are seven indicators.

**Two forms.** For MSA, BHI and PDF, the coefficients $c_k^X$ given by the original sources for each land-cover type (the coefficient of indicator $X$ for class $k$) were weighted by the land-cover mix of the survey unit and summed (coefficient versions; coefficients in Supplementary Table 3). The value of survey unit $u$ is $\sum_k a_{uk}\, c_k^X$, where $a_{uk}$ is the area share of class $k$ in the 2020 land-cover mix of unit $u$. For the two versions of BII, BHI and LIFE, the 2020 values of the published maps were averaged over the unit (map versions). BHI is published in both forms and was compared in both. STAR was rebuilt for birds to match the conditions of the evaluation (2020 land cover; breeding and resident distributions; see Rebuilding STAR below). Separately from the indicators of land-use impact, the IUCN AoH richness map for birds was compared in the same way as the map versions.

#### Land-cover coefficients

**Supplementary Table 3 Coefficients c by land-cover class in the coefficient versions.** Natural classes (IGBP 1–11 and 15–17) are 1 for all indicators.

| Indicator | Cropland (12) | Urban (13) | Mosaic (14) | Source |
|---|---|---|---|---|
| MSA | 0.427 | 0.240 | Mean of cropland and natural | GLOBIO v4.3.1, birds and mammals[10] |
| BHI | 0.035 | 0 | Mean of cropland and natural | State coefficients of BHI v4 (annual crops and urban; Table S1.4 of the BHI v4 methods document)[11] |
| PDF | 0 | Equation below | Mean of cropland and natural (0.5) | Bird characterization factors v1.0.1 of Scherer et al.[5,12] |

The MSA coefficients are the GLOBIO land-use coefficients (cropland 0.296, urban 1.15) entered in the GLOBIO equation (for cropland, $1/(1+e^{0.296})$). The BHI state coefficients are the original BHI coefficients (cropland 0.432, urban 0) raised to the fourth power, following the definition in the same document (the reciprocal of the exponent z = 0.25).

The urban coefficient of PDF expresses the urban loss in an ecoregion as a ratio to the cropland loss:

$$c_{13,e}^{\mathrm{PDF}} = 1 - \frac{\mathrm{CF}_e(\text{urban, light})}{\mathrm{CF}_e(\text{cropland, light/intense})}$$

Here $\mathrm{CF}_e$ is the bird characterization factor for regional average occupation of Scherer et al.[5]. Because the cropland loss was set to a value of 1 common to the ecoregion, the cropland coefficient is 0 (a common multiple does not change rank correlations or increases in the coefficient of determination). Since the shares of a land-cover mix sum to 1, this coefficient version is equivalent to a linear score of 1 minus the losses of cropland, urban and mosaic (the urban loss being the above ratio and the mosaic loss 0.5). For land-use intensity, the light coefficient was used for urban. For cropland, the animal coefficients of the source have only one category that does not separate light and intense use (light/intense), and that coefficient was used. Wetland and water have no corresponding class in PDF and were treated as having a coefficient of 1.

The regional divisions of PDF (WWF 2001 ecoregions)[13] were matched to each ecoregion of this study[14] by the division with the largest overlap on the 0.05° grid. The share of the ecoregion area covered by the matched division, as a median across ecoregions with compared survey units, was 0.66 in southern Africa (minimum 0.31), 0.95 in Australia and 0.87 in North America (southern Africa has the largest differences in how the two sets of boundaries are drawn). Ratios could be built for all compared survey units. However the urban ratio is set, the coefficient version of PDF ranks units within an ecoregion almost as natural-cover share does (main text), so the coarseness of this matching barely affects the results.

Predictors from the coefficient versions were used as proportions without logs (because the urban coefficient of BHI is 0, and the log of a fully urban unit cannot be taken).

### Map versions and calculation of unit values

**Supplementary Table 4 Versions of the map indicators.** All use 2020 values, averaged over the unit with area weights (weighted by the share of the unit covered by each pixel). Direction shows whether the values were oriented so that larger values mean less modification (the sign was reversed for indicators in which larger values are worse).

| Indicator | Version and source | Resolution | Values used | Direction |
|---|---|---|---|---|
| BII (IO) | v1.1 of Impact Observatory and Vizzuality (an Earth Engine copy, confirmed to differ from the original map by at most 0.005 in 60 random units)[6] | About 100 m | 0–1 | Unchanged |
| BII (NHM) | v2.1.1 of the Natural History Museum, London (limited release)[15] | 5 arcmin | 0–100 rescaled to 0–1 | Unchanged |
| BHI | v4 of CSIRO[11] | About 1 km | bhi layer (0–1) | Unchanged |
| LIFE | v1.01 of Eyres et al.[8,16], birds | 1 arcmin | Change in extinctions per unit area when natural habitat is converted to arable land (layer with exponent 0.25) | Unchanged |
| STAR | Bird version rebuilt in this study (below) | 0.05° | Score per unit area | Sign reversed |
| IUCN AoH richness | IUCN Species Richness from AoH rasters, Birds (Red List 2021-3)[17,18]; the Mollweide map transferred to the 0.05° grid by nearest neighbour | 5 km | Number of species | Unchanged |

LIFE also has a restoration layer, but it has values only for units with cropland, which would change the set of units, so it was not used.

### Rebuilding STAR

STAR was rebuilt for birds from public materials to match the conditions of the evaluation (2020 land cover; breeding and resident distributions). The value of 5 km cell $g$ is

$$\mathrm{STAR}_g = \sum_s W_s \frac{\mathrm{AoH}_{s,g}}{\mathrm{AoH}_s}$$

Here $W_s$ is the Red List category weight of species $s$ (Near Threatened 1, Vulnerable 2, Endangered 3, Critically Endangered 4, Least Concern 0), $\mathrm{AoH}_{s,g}$ is the area of habitat (AoH)[19] of the species in the cell and $\mathrm{AoH}_s$ is the AoH summed over the whole range of the species. The original divides the score by threat, but the threat proportions of a species sum to 1, so the score across all threats is given by this equation. Because the Red List version, habitat map and construction of AoH differ from the original, this is not the official STAR. The official STAR[9] uses Red List 2019-3, ESA CCI land cover for 2015 and 5 km AoH.

The target species were 317 birds that were Near Threatened to Critically Endangered in the 2024 Red List[20] and whose breeding or resident BirdLife range[21] falls in the three survey regions; 314 species with positive AoH (122 Near Threatened, 94 Vulnerable, 77 Endangered and 21 Critically Endangered) contributed to the values. The AoH of species $s$ in cell $g$ is the product of three shares (distinguished by superscripts) and the area of the cell:

$$\mathrm{AoH}_{s,g} = A_g\, \eta_{s,g}^{\mathrm{rng}}\, \eta_{s,g}^{\mathrm{hab}}\, \eta_{s,g}^{\mathrm{elv}}, \qquad \mathrm{AoH}_s = \sum_g \mathrm{AoH}_{s,g}$$

Here $\eta_{s,g}^{\mathrm{rng}}$ is the share of the cell covered by the range of the species (breeding and resident), $\eta_{s,g}^{\mathrm{hab}}$ is the share of the 500 m pixels of the cell in MODIS classes (2020) that correspond to the species' habitat, and $\eta_{s,g}^{\mathrm{elv}}$ is the share of the cell within the elevation range of the species. $\eta^{\mathrm{hab}}$ was built from a table linking IUCN habitat types to MODIS classes (Supplementary Data 1; see Correspondence between habitats and MODIS classes below); species with no habitat types were not restricted by habitat (species that occur only at sea were excluded). $\eta^{\mathrm{elv}}$ was calculated from the mean $\overline{\mathrm{el}}_g$ and standard deviation $\mathrm{sd}_g^{\mathrm{el}}$ of cell elevation (0.05° summaries of GMTED2010)[22] as

$$\eta_{s,g}^{\mathrm{elv}} = \Phi\left(\frac{\mathrm{el}_s^{\max} - \overline{\mathrm{el}}_g}{\mathrm{sd}_g^{\mathrm{el}}}\right) - \Phi\left(\frac{\mathrm{el}_s^{\min} - \overline{\mathrm{el}}_g}{\mathrm{sd}_g^{\mathrm{el}}}\right)$$

where $\mathrm{el}_s^{\min}$ and $\mathrm{el}_s^{\max}$ are the lower and upper elevation limits of the species and Φ is the standard normal distribution function (0 or 1 if the limits are outside the lowest or highest elevation of the cell). The denominator $\mathrm{AoH}_s$ was summed over the whole range of the species, not only within the survey regions. The value of a unit is the mean of $\mathrm{STAR}_g/A_g$ weighted by the share of the unit covered by each cell (score per unit area). The weights $W_s$ used the 2024 Red List categories in version 9 of the Handbook of the Birds of the World and BirdLife checklist[23].

### Correspondence between habitats and MODIS classes

This table is used only to determine the MODIS classes corresponding to species' habitat ($\eta^{\mathrm{hab}}$) in the STAR rebuilt above. It assigns MODIS IGBP classes to each of 126 IUCN habitat types (18 level 1 types and their level 2 subdivisions) and was built following Lumbierres et al.[1] (Supplementary Data 1).

**Source table.** Lumbierres et al.[1] estimated associations between IUCN habitat types (level 1, with artificial terrestrial habitats split into arable land and pasture, degraded forest and plantations, and urban and rural gardens) and ESA-CCI land-cover classes from observation records by odds ratios. They divided significantly positive odds ratios into three levels by tertiles (lower bounds 1.121, 1.396 and 1.708) and left the choice of level to users.

**Translation.** I transcribed that ESA-CCI table and translated each ESA-CCI class to the MODIS class closest in definition (four cropland classes to 12; two cropland/natural mosaic classes to 14; five tree classes to 1–5; tree and shrub mosaic to 8; herbaceous mosaic to 9; shrubland to 6 and 7; grassland and lichens and mosses to 10; sparse vegetation and bare areas to 16; flooded tree, shrub and herbaceous cover to 11; urban to 13; water to 17). For each habitat, the MODIS classes of a habitat were the translations of ESA-CCI classes with positive associations at or above the lower bound of the middle level (1.396). Level 2 types inherited the result of their level 1 parent. The 53 types not treated by Lumbierres et al. (caves, marine habitats, artificial marine structures and others) were left with no correspondence. Snow and ice (15) has no row in the ESA-CCI table and is in no habitat.

**Why the middle level, and classes added by hand.** Lumbierres et al.[1] compared the share of bird observation points falling in the AoH built at each level with the share for AoH built from an expert correspondence. The middle level is the closest to the expert correspondence. The highest level would also reduce savanna to shrubland (6, 7), leaving MODIS savanna (8, 9) without a correspondence. To forest I added deciduous needleleaf forest (MODIS 3), for which no significant association was estimated in the table of Lumbierres et al. (the same paper recommends that users add classes whose associations could not be estimated).

#### Decomposing the values of the AoH richness map and STAR

In the comparison in the main text (Fig. 2a), the IUCN AoH richness map ranked differences in field richness about as well as the richness estimate, whereas STAR ranked them hardly at all. I examined the reasons with two decompositions. Rank correlations were calculated by the same method and on the survey units of Fig. 2a in the main text that have IUCN AoH values (3,105, 1,502 and 1,349; Supplementary Table 11), and intervals are 90% intervals from a bootstrap over ecoregions (2,000 replicates). Values are in the order southern Africa, Australia and North America. The 90% intervals of the difference in rank correlation between the richness estimate and the AoH richness map (richness estimate minus AoH) were −0.27 to −0.08, −0.12 to 0.22 and −0.07 to 0.13.

**Land-cover and location parts of the AoH map.** I regressed the log of AoH richness of the units on ecoregion fixed effects and the shares of the 17 classes of the unit. The fitted values minus the mean fitted value of the units in the ecoregion form the land-cover part, and the residuals form the location part (range edges, elevation and other effects not explained by the land-cover mix). The mix explained 0.14, 0.16 and 0.11 of the within-ecoregion variance. The location part ranked field richness better than the land-cover part in all three surveys: rank correlations were 0.27, 0.18 and 0.23 for the whole AoH map, 0.14, 0.10 and 0.08 for the land-cover part and 0.23, 0.13 and 0.20 for the location part. Compared with the land-cover part, the richness estimate (0.12, 0.21 and 0.26) was higher in North America and similar in the other two regions (90% intervals of the difference, estimate minus land-cover part, −0.09 to 0.03, −0.01 to 0.25 and 0.06 to 0.30).

**Steps in building STAR.** I compared the quantities obtained by changing one step at a time between AoH richness of all species and STAR: (i) AoH richness of all species; (ii) the number of threatened and Near Threatened species (Near Threatened to Critically Endangered); (iii) the sum over these species of the share of the unit area covered by the AoH of each species; (iv) (iii) weighted by category; and (v) STAR (the term of each species in (iv) divided by the total AoH of the species). Quantity (i) is the IUCN map, and (ii)–(v) were built from the species AoH rebuilt in this study. Rank correlations (positive where larger values correspond to higher field richness; unlike Fig. 2a in the main text, the sign of STAR was not reversed) were 0.27, 0.18 and 0.23 for (i), 0.10, 0.08 and 0.08 for (ii), 0.07, 0.01 and −0.06 for (iii), 0.05, 0.01 and −0.06 for (iv) and 0.04, −0.06 and −0.02 for (v). The largest declines came with the restriction to these species ((i) to (ii); 90% intervals of the difference 0.08 to 0.24, −0.12 to 0.24 and 0.05 to 0.25) and, in North America, with the change to a sum of shares ((ii) to (iii); 0.09 to 0.20).

**Quality of the rebuilt AoH.** Richness from stacking the species AoH rebuilt in this study for all species had rank correlations with the IUCN map on 0.05° cells of 0.77 globally, 0.86 in southern Africa, 0.79 in Australia and 0.27 in North America (the low value in North America probably arises because this study uses only breeding and resident ranges, whereas the IUCN map also includes wintering ranges). Rank correlations with field richness were 0.24, 0.12 and 0.15, and the 90%

intervals of the differences from the IUCN map all included 0 (−0.11 to 0.03, −0.16 to 0.09 and −0.19 to 0.03). STAR's mismatch with richness therefore stems not from the quality of the rebuilt AoH but from its definition, the opportunity to reduce threats to threatened and Near Threatened species.

### Supplementary Note 3. The composition estimate (a model of the probability that the lists of two cells share a species) and comparison of richness and composition under alternative natural references

This note gives the details of the composition estimate, which models the number of species shared by the lists of two cells from the land-cover mixes of the two cells (the sharing model of the Methods and main text), and the definitions of the quantities under alternative natural references (richness ratio and species retention) built from this estimate and the richness estimate (Supplementary Note 8). Other notes refer to these definitions.

If $p_{gs}$ is the probability that species $s$ is on the standardized list of cell $g$ (Supplementary Note 10), the list length (expected number of species) is $F'_g = \sum_s p_{gs}$, and the expected number of species shared by the lists of two cells $g$ and $g'$ in the same ecoregion is $G_{gg'} = \sum_s p_{gs}\, p_{g's}$. The sharing probability $P_e(a \mid a', \ell)$ is the probability that a species on the list of a cell with mix $a'$ is also on the list of a cell with mix $a$ at distance $\ell$:

$$P_e(a \mid a', \ell) = \sum_{k'} \pi_{k',e}(a') \left[1 - \exp\left(-\delta_e(\ell) \sum_k \lambda_{k|k',e}\, a_k^z\right)\right], \qquad \delta_e(\ell) = \left(\frac{\ell}{\ell_0}\right)^{-\zeta_e}$$

Here $\pi_{k',e}(a')$ is the share of species originating in class $k'$ (see Origin shares below). $\lambda_{k|k',e} \geq 0$ is the strength with which species originating in class $k'$ also occur on the list of a cell of class $k$ (hereafter persistence strength; in a cell consisting only of class $k$, the probability of occurring is $1 - e^{-\lambda}$, so larger values mean a greater probability). $\delta_e(\ell)$ is the term for the decline in sharing with distance (distance decay), with exponent $\zeta_e$. $\ell_0$ is the median distance between pairs of natural cells (cells with a natural-cover share of at least 0.95; or the median distance of all pairs in ecoregions with fewer than 20 such pairs), and $z$ is the mixing exponent of the richness estimate (Supplementary Note 8). So that the decline in sharing with distance is the same in all comparisons, all predictions are evaluated at distance $\ell_0$.

#### Comparison of richness and composition under alternative natural references

To assess land of anthropogenic class $h$ (cropland, urban or cropland/natural mosaic), I compared it with a hypothetical cell consisting only of natural classes of the ecoregion (natural reference cell). The land-cover mix $\bar{a}_e$ of the all-natural reference cell is the 2020 area of the natural classes of the ecoregion with usable richness weights, rescaled to sum to 1 across natural classes (if the natural part of an ecoregion is 60% forest and 40% grassland, so is the reference cell). To vary the natural state compared, I also restricted $\bar{a}_e$ to forest alone (IGBP 1–5) or to open natural classes alone (6–10) and calculated the same quantities (ecoregions where that part has no area are excluded). The natural reference cell is a hypothetical cell, separate from real natural cells (cells with a natural-cover share of at least 0.95). The mosaic is not used as a reference. Writing the mix consisting only of class $h$ as $\mathbf{1}_h$, I calculated

$$\rho_{h,e} = \frac{w_{h,e}}{F_e(\bar{a}_e)}, \qquad \tilde{r}_e(a \to a^*) = \frac{P_e(a^* \mid a, \ell_0)}{P_e(a \mid a, \ell_0)}$$

Here $w_{h,e}$ is the weight of class $h$ in the richness estimate (the expected richness of a cell consisting only of class $h$) and $F_e$ is expected richness from the richness estimate, both defined in Supplementary Note 8.

The **richness ratio** $\rho_{h,e}$ is the expected richness of a cell consisting only of class $h$ divided by the expected richness of the natural reference cell.

**Species retention** $\tilde{r}_e(a \to a^*)$ is the share of the species on the list of a cell with mix $a$ that are also on the list of a cell with mix $a^*$, divided by the same share between two cells with mix $a$. Even two cells with the same mix do not share all species, because lists are incomplete and species turn over with distance; the denominator cancels this part, and a retention of 1 means that a cell with mix $a^*$ holds as many of the species of a cell with mix $a$ as another cell with mix $a$ does. Fig. 3 of the main text shows the retention $\tilde{r}_{h,e}$ of the natural reference cell's species in the anthropogenic cell ($a = \bar{a}_e$, $a^* = \mathbf{1}_h$), and Extended Data Fig. 3 shows the reverse direction, the retention of the anthropogenic cell's species in the natural reference cell ($a = \mathbf{1}_h$, $a^* = \bar{a}_e$). Unless the direction is stated, retention means $\tilde{r}_{h,e}$.

**Point estimates and intervals.** These quantities were calculated only for combinations of ecoregion and class with sufficient data of the ecoregion itself for both richness and composition (for richness, usable weights with at least five recorded cells in which the class covers at least half the area [Supplementary Note 8]; for composition, those meeting the two conditions under Values used in the results below), and, so that the references could be compared, only for combinations in which all three references could be built. The 400 draws of each of the two estimates (with model error added) were paired by number and calculated per draw and per ecoregion, and the distribution of the 400 per-draw ecoregion medians was summarized by its median (point estimate) and its 5th to 95th percentiles (90% interval).

**Differences between the mosaic and cropland.** Differences in retention (mosaic minus cropland) were taken per draw in ecoregions with both classes and summarized in the same way. Retention was higher on the mosaic than on cropland for species of the forest reference, by +0.08 for birds (90% interval +0.02 to +0.14), +0.18 for mammals (+0.05 to +0.34), +0.20 for reptiles (+0.02 to +0.43) and +0.35 for amphibians (+0.13 to +0.56); those for species of the open natural reference were +0.04 for birds (−0.05 to +0.12), +0.06 for mammals (−0.10 to +0.26), +0.13 for reptiles (−0.05 to +0.32) and +0.17 for amphibians (−0.04 to +0.40).

### Why retention was chosen as the index of composition

From the composition estimate, this study builds retention, which expresses how much of a natural community's species also remains in a cell of anthropogenic land. There are many indices of composition, including Jaccard, Sørensen, Bray–Curtis and Morisita–Horn. GBIF records represent only occurrence, which limits the choice to presence/absence indices (time-series studies also measure presence/absence turnover)[24]. Symmetric similarities mix differences in richness with species replacement, whereas a share taken from the side of the reference community directly expresses how many of its species remain. Dividing by the sharing between reference cells also cancels the incompleteness common to both lists. Because the frequencies of GBIF records represent not abundance but recordability, including detectability and observer behaviour, I did not extend the analysis to indices weighted by species abundance (Horn overlap and Morisita–Horn similarity).

### The full sharing model and its estimation

**Observations.** Pairs of cells 5–500 km apart in the same ecoregion were selected, up to 20,000 pairs per ecoregion and 50 uses of any one cell, with at least a quarter of pairs including a cell in which an anthropogenic class covers at least half the area. Each pair yields two observations (directed pairs), the number of species on the list of $g'$ also in $g$ and the reverse, each with a weight of 0.5 so that the same pair is not counted twice. Ecoregions with fewer than 50 observations were not estimated. Numbers of cell pairs and ecoregions are given in Supplementary Table 10.

**Likelihood.** The pseudo-log-likelihood of a directed pair $(g, g')$ is the log-likelihood of a beta-binomial distribution with the real-valued number of trials $F'_{g'}$ and number of successes $G_{gg'}$ entered directly (the quasi-likelihood idea of placing non-integer responses in a likelihood form):

$$\log\mathcal{L}_e = \frac{1}{I_e} \sum_{(g,g')} 0.5 \ \log\mathrm{BB}\left(G_{gg'} \mid F'_{g'},\ P_e(a_g \mid a_{g'}, \ell_{gg'}),\ \phi_{gg'}\right)$$

Here BB is the beta-binomial probability (extended to real values through gamma functions) and $I_e$ is the jackknife inflation factor (see Estimation and draws below), which lowers the weight of the likelihood accordingly (the weight is unchanged if it is below 1). For the precision $\phi_{gg'}$, the logit of the correlation among species of the same list, $1/(1 + \phi_{gg'})$, is a linear function of the log list length:

$$\operatorname{logit}\frac{1}{1+\phi_{gg'}} = \phi_{0,e} + \phi_{1,e}\ln F'_{g'}$$

Here $\phi_{0,e}$ and $\phi_{1,e}$ are the intercept and slope of each ecoregion ($\ln F'_{g'}$ is centred within the ecoregion). Because the two lists are standardized to the same coverage, the model has no term correcting for species missed in poorly recorded cells.

**Origin shares.** The term $\pi_{k',e}(a')$ in the sharing probability is the share of the contribution of class $k'$ when list length is expressed as a function of the land-cover mix:

$$\pi_{k',e}(a') = \frac{w'_{k',e}\, {a'}_{k'}^{z}}{\sum_k w'_{k,e}\, {a'}_{k}^{z}}, \qquad \ln w'_{k,e} = \ln w_{k,e} + \varepsilon^{\mathrm{grp}}_{\mathrm{grp}(k),e} + \varepsilon^{\mathrm{cls}}_{k,e}$$

Here $w'_{k,e}$ is the class weight for standardized list length, $w_{k,e}$ is the weight of the richness estimate (posterior mean), $\mathrm{grp}(k)$ is the origin group of class $k$ (Supplementary Table 5), $\varepsilon^{\mathrm{grp}}$ is a correction for each origin group and $\varepsilon^{\mathrm{cls}}$ is a correction for each class. $w'$ was estimated by fitting the log list length $\ln F'_g$ to $\ln \sum_k w'_{k,e}\, a^{z}_{gk}$ (normal errors; the variance was set from the residuals of a fit with corrections for origin groups alone, with normal priors of mean 0 and standard deviation 3 for $\varepsilon^{\mathrm{grp}}$ and mean 0 and standard deviation 0.5 for $\varepsilon^{\mathrm{cls}}$). $z$ was taken directly from the richness estimate (0.88 for birds).

**Nesting of persistence strengths.** The destination $k$ stays at 17 classes, while the origin classes $k'$ are grouped into the eight origin groups of Supplementary Table 5, and class strengths are drawn towards the strength of the origin group:

$$\ln\lambda_{k|k',e} = \ln\lambda_{k|\mathrm{grp}(k'),e} + \varepsilon^{\lambda}_{kk',e}, \qquad \varepsilon^{\lambda}_{kk',e} \sim \mathcal{N}(0,\ 0.5^2)$$

Here $\varepsilon^{\lambda}$ is the difference among classes within an origin group. Without data to support such differences, $\varepsilon^{\lambda}$ shrinks to 0 and the result equals an estimate with the eight origin groups alone. Results were almost the same in simulated data when the standard deviation of 0.5 was changed to 0.25 or 1.0.

**Supplementary Table 5 The eight origin groups.** Numbers are IGBP land-cover class numbers.

| Origin group | Classes |
|---|---|
| Forest | 1 Evergreen needleleaf forest, 2 Evergreen broadleaf forest, 3 Deciduous needleleaf forest, 4 Deciduous broadleaf forest, 5 Mixed forest |
| Woodland and shrubland | 6 Closed shrubland, 7 Open shrubland, 8 Woody savanna |
| Open vegetation | 9 Savanna, 10 Grassland |
| Waterside | 11 Permanent wetland, 17 Water |
| Barren and snow/ice | 15 Snow and ice, 16 Barren land |
| Cropland | 12 Cropland |
| Urban | 13 Urban and built-up |
| Cropland/natural mosaic | 14 Cropland/natural vegetation mosaic |

**Priors.** In the first fit, the log strengths of origin groups had a normal prior with mean $\ln(-\ln 0.7)$ and standard deviation 2 (the mean corresponds to a probability of 0.3 that a species of class $k'$ is on the list of a cell consisting only of class $k$), the distance-decay exponent $\zeta_e$ had mean 0 and standard deviation 1, the intercept $\phi_{0,e}$ of the precision equation had mean $-3$ and standard deviation 3, and the slope $\phi_{1,e}$ had mean 0 and standard deviation 1. In the second fit, the prior on the log strengths of origin groups was replaced by that described under Shrinkage within biomes below.

**Estimation and draws.** I maximized the sum of the pseudo-log-likelihood and the log priors, and drew 400 sets of parameter values from a multivariate normal distribution built from the Hessian at that point (Laplace approximation; hereafter the posterior). Observations share cells and are not independent. I calculated the variance by a ten-block jackknife over cells and lowered the weight of the likelihood by the ratio to the model variance (inflation factor $I_e$) before the normal approximation (a correction for a likelihood that treats observations as independent)[25].

#### Shrinkage within biomes

For the log strengths of origin groups (per combination of destination $k$ and origin group), each ecoregion was first fitted independently with the weak priors above. The prior mean $\bar{l}_m$ and standard deviation $\sigma_{\lambda,m}$ of each combination were then reset from the estimates of the ecoregions of the same biome $m$ (the 14 units above RESOLVE ecoregions)[14] (empirical Bayes). $\sigma_{\lambda,m}$ combines the between-ecoregion variation within biomes, $\tau_\lambda$ (one value across combinations), with the uncertainty in the estimate of the biome mean. Finally, the model was fitted again with this prior. The fewer the records of an ecoregion, the more strongly it is drawn to the biome mean.

Shrinkage was required to meet two conditions. (1) The prior must not conflict with the independent estimates of each ecoregion: for the independent estimate $\hat{l}$ (the value from the first fit with the influence of the weak prior removed) and its standard error, the share of combinations with independent estimates that satisfies the following inequality must lie between 0.80 and 0.97 (0.90 for a correct prior; it comes out slightly higher because each ecoregion also contributes to the estimate of the prior).

$$\left|\hat{l} - \bar{l}_m\right| < 1.645\sqrt{\mathrm{se}(\hat{l})^2 + \sigma_{\lambda,m}^2}$$

(2) In ecoregions with fewer observations than the median of the taxon, the median ratio of the posterior standard deviation after shrinkage to that before must be at most 0.8. For birds, the share in (1) was 0.942 and the median ratio in (2) was 0.28, meeting both conditions.

#### Values used in the results

Only values of retention $\tilde{r}$ that can be estimated precisely and for which data directly compare the class and natural classes were used. The two conditions were: (i) the posterior standard deviation of retention is at most 0.5 times the estimate of retention; and (ii) among the cell pairs used in the estimation, at least five have the class covering at least 0.5 of the area of one cell and one of the natural origin groups (Supplementary Table 5) covering at least 0.5 of the other. Condition (i) was judged on retention against all natural classes and applied also to the forest-only and open-natural-only references. For the quantities under alternative natural references (above), only classes with draws of the richness estimate were used, and combinations of ecoregion and anthropogenic class with non-finite values in the draws were excluded (for birds, eight ecoregions for cropland, eight for urban and two for the mosaic under the forest-only reference, and none under the all-natural and open-natural-only references).

### Supplementary Note 4. Details of global change from 2020 to 2024

This note gives the details of the calculation of global change from 2020 to 2024 (construction of land-cover mixes, cells receiving values, weighting, contributions by transition type and biome, and maps). Writing the land-cover mixes of 5 km cell $g$ in 2020 and 2024 as $a_g$ and $b_g$, I calculated the expected-richness ratio $\Lambda_g = F_e(b_g)/F_e(a_g)$ ($F_e$ is expected richness from the richness estimate; Supplementary Note 8) and the compositional change ratio ($P_e$ is the sharing probability of the composition estimate; Supplementary Note 3):

$$\Gamma_g = \frac{P_e(b_g \mid a_g, \ell_0)}{P_e(a_g \mid a_g, \ell_0)}$$

$\Gamma_g$ is the share of the species of a community with the 2020 land-cover mix predicted to be found in a cell with the 2024 mix, relative to no change. Global values are weighted means, with weights equal to the product of area and 2020 expected richness for expected richness and to land area for composition (Methods).

#### Details of the land-cover mixes

- The mixes for calculating change are built separately from those of the estimation (Supplementary Note 7). The 500 m pixels were transferred to a 0.005° grid by nearest neighbour, giving 100 pixels per 5 km cell. The mixes at two times, $a_g$ and $b_g$, were built from the integer counts of pixels that kept one class in all five years or changed persistently between 2020–2021 and 2022–2024 (together, consistently classified pixels). Classes with a share below 0.02 (fewer than two pixels) were treated as absent.

- The 648 cells without any stable pixel (all pixels alternating between years) were counted as unchanged ($\Lambda_g = 1$) for expected richness and excluded as cells without values for composition.
- The 2020 expected richness of unchanged cells was calculated only from classes with usable weights.
- After classes below 0.02 were treated as absent, shares were rescaled to sum to 1 for composition but not for expected richness ($F_e$ was calculated from the original shares of the remaining classes).
- Within one cell, a change from one class to another and the reverse change cancel in the land-cover mix. Values depend only on the mixes at the two times, not on the combination of pixel transitions.

### Breakdown of cells without values

Expected richness could not be calculated for changed cells when any class appearing at either time lacked a usable weight (Supplementary Note 8). The population $\mathcal{G}_\Lambda$ for the global value of expected richness consists of unchanged land cells and changed cells with values. For composition, values could not be calculated because the ecoregion had no fit or did not converge (5.1% of changed cells for birds) or because a class appearing in the cell had no value usable in the results (1.8%).

**Cells used for global values.** Global values in the main text (Fig. 4c,d) use, among cells with map values, only cells in which every class appearing is supported by the data of its own ecoregion (for expected richness, at least five recorded cells in which the class covers at least half the area; for composition, at least five pairs in which the class covers at least 0.5 of the area of one cell and one of the natural origin groups [Supplementary Table 5] covers at least 0.5 of the area of the other; Methods gives the share of area meeting the conditions). Global change in expected richness of birds barely differed with or without this condition (−0.0696% with it, as in the main text; −0.0701% without it). Global change in composition, compared before adding model error, was −0.077% with the condition and about twice that, −0.138%, without it, with most of the difference coming from cells containing classes with few pairs with natural classes. The breakdowns by weighting, transition and biome below were calculated on cells with map values.

### Four weighting schemes for global change in expected richness

The cell-level change $\Lambda_g - 1$ was averaged (1) without weights, (2) with area (cosine of latitude), (3) with 2020 expected richness, and (4) with area × 2020 expected richness (the weights used in the main text). The target cells are the same in all four ($\mathcal{G}_\Lambda$). Scheme (3) equals the rate of change in the sum of expected richness over the cells of $\mathcal{G}_\Lambda$, and (4) the rate of change in the sum multiplied by area. In Supplementary Table 6, the increase in reptiles becomes a decrease when weighted by expected richness, whereas the increase in amphibians remains. For birds, weighting by expected richness makes the decrease more than twice that without weights.

**Supplementary Table 6 Global change in expected richness by weighting scheme (2020 to 2024, %).** Values from all land-cover change (for changes involving anthropogenic classes only, see the section of that name below). Values are at the posterior mean of the class weights. Because they are for all cells with map values ($\mathcal{G}_\Lambda$), they differ for taxa other than birds from the global values in the main text, which use only cells supported by data (for mammals, −0.049% against −0.032%).

| Taxon | No weights | Area | Expected richness | Area × expected richness |
|---|---|---|---|---|
| Birds | −0.029 | −0.044 | −0.070 | −0.070 |
| Mammals | −0.063 | −0.062 | −0.056 | −0.049 |
| Reptiles | +0.016 | +0.009 | −0.016 | −0.011 |
| Amphibians | +0.005 | +0.004 | +0.025 | +0.026 |

### Contributions by transition type and biome

**Transition types.** The 17 classes were grouped into 11 (forest [five classes], shrubland [two], savanna [woody savanna and savanna], grassland, wetland, cropland, urban, mosaic, snow and ice, barren land and water), and a pair of groups before and after change is called a transition pair. For each transition pair (for example, savanna to forest), I built a 2024 land-cover mix in which only the changes of pixels in that pair occurred and all other changes were undone, and calculated the global value with the weights of the main text (area × 2020 expected richness) as the contribution of the pair. Changes within one group (among forest types, for example) were combined in one transition pair. Because Λ is non-linear in the mix, the sum of the contributions does not match the overall value (−0.074% against −0.070% for birds in Supplementary Table 7).

**Biomes.** The overall change was divided by the biome of the cell's ecoregion (the numerator was summed by biome and divided by the same denominator). This division is additive, and the contributions of the biomes sum to the overall value. Of the decrease for birds (−0.070%), −0.044% came from tropical and subtropical grasslands, savannas and shrublands. The increase for amphibians (+0.026%) came mainly from tropical and subtropical dry broadleaf forests (+0.019%), moist broadleaf forests (+0.011%) and grasslands, savannas and shrublands (+0.011%), and contributions were negative in temperate, Mediterranean and desert biomes.

**Supplementary Table 7 Contributions by transition type to global change in expected richness (2020 to 2024, %).** Weights of area × 2020 expected richness; values at the posterior mean of the class weights. There are 97 transition pairs in all. Pairs not involving anthropogenic classes are the top nine by the maximum absolute contribution across the four taxa, plus barren land to grassland, the reverse of grassland to barren land. Pairs involving anthropogenic classes are the top five ordered in the same way.

| Transition pair | Anthropogenic classes | Birds | Mammal s | Reptiles | Amphibians |
|---|---|---|---|---|---|
| Savanna to forest | Not involved | +0.0464 | +0.0539 | +0.0545 | +0.0644 |
| Forest to savanna | Not involved | −0.0614 | −0.0279 | −0.0200 | −0.0189 |
| Grassland to savanna | Not involved | +0.0040 | +0.0258 | +0.0295 | +0.0911 |
| Savanna to grassland | Not involved | −0.0188 | −0.0361 | −0.0251 | −0.1190 |
| Grassland to shrubland | Not involved | −0.0294 | −0.0119 | +0.0000 | −0.0005 |
| Shrubland to grassland | Not involved | +0.0160 | −0.0008 | −0.0365 | +0.0092 |
| Grassland to barren land | Not involved | −0.0160 | −0.0158 | −0.0050 | −0.0044 |
| Barren land to grassland | Not involved | +0.0105 | +0.0045 | +0.0022 | +0.0030 |
| Forest to grassland | Not involved | −0.0037 | −0.0079 | −0.0073 | −0.0141 |
| Change within one group | Not involved | −0.0111 | −0.0223 | −0.0095 | −0.0038 |
| Grassland to cropland | Involved | −0.0011 | −0.0034 | −0.0088 | +0.0209 |
| Cropland to grassland | Involved | −0.0062 | +0.0059 | +0.0095 | −0.0122 |
| Mosaic to cropland | Involved | +0.0012 | −0.0032 | −0.0077 | +0.0056 |
| Savanna to mosaic | Involved | −0.0057 | −0.0057 | −0.0030 | −0.0059 |
| Savanna to cropland | Involved | −0.0018 | −0.0042 | −0.0030 | −0.0025 |
| Sum of pairs involving anthropogenic classes | | −0.0115 | −0.0066 | −0.0095 | +0.0055 |
| Sum of all 97 pairs | | −0.0738 | −0.0514 | −0.0133 | +0.0243 |
| Overall value (all cells with map values) | | −0.0701 | −0.0495 | −0.0111 | +0.0263 |

In Supplementary Table 7, the positive value for amphibians comes mainly from the net of savanna and forest transitions (+0.0644 − 0.0189 = +0.046%) and, among pairs involving anthropogenic classes, from grassland to cropland (+0.021%). Savanna–forest transitions occur at almost the same scale in both directions globally (1.11 million persistently changed 500 m pixels from savanna to forest and 1.06 million in reverse), and such symmetrical exchanges cannot be distinguished from noise at the classification boundary. In the unweighted mean (No weights in Supplementary Table 6), the increases in reptiles and amphibians came mainly from savanna and forest transitions in boreal forests. When the unweighted mean was divided by transition pair and biome in the same way (not shown in the table), boreal forests accounted for +0.030% (reptiles) and +0.015% (amphibians) of the savanna and forest contribution (+0.045% and +0.029%).

#### Version with only changes involving anthropogenic classes

To see how far global values move with only changes involving anthropogenic classes, I also calculated global values with a 2024 land-cover mix in which all persistent changes between natural classes (IGBP 1–11 and 15–17) were undone (orange triangles in Fig. 4c,d of the main text). The undone pixels were 88.3% of persistently changed pixels. Global change for birds was −0.010% in expected richness and −0.009% in composition.

#### Extent of map values relying on biome means

In ecoregions where the composition model could not be estimated (too few records, or the fit did not converge), map values of cells were filled with the mean persistence strengths built from fits in the same biome. With this filling, the share of changed area with map values rose from 93.2% to 97.8% for birds, 63.2% to 91.1% for mammals, 46.2% to 85.2% for reptiles and 45.7% to 93.1% for amphibians. I checked the accuracy of the filling by removing each ecoregion with a fit in turn, filling it with the biome mean and comparing the result with the fit of that ecoregion. The rank correlations were 0.39–0.50 for the mean change of an ecoregion but only 0.01–0.38 for retention. Filled values should be read as the typical direction and size of change in the biome, not as differences among ecoregions. Expected-richness weights are also determined mainly by the class effects of the biome in ecoregions with few recorded cells in which the class covers at least half the area (Supplementary Note 8). Supplementary Fig. 3 shows, for each map grid cell, the share of cell area relying on such unsupported classes.

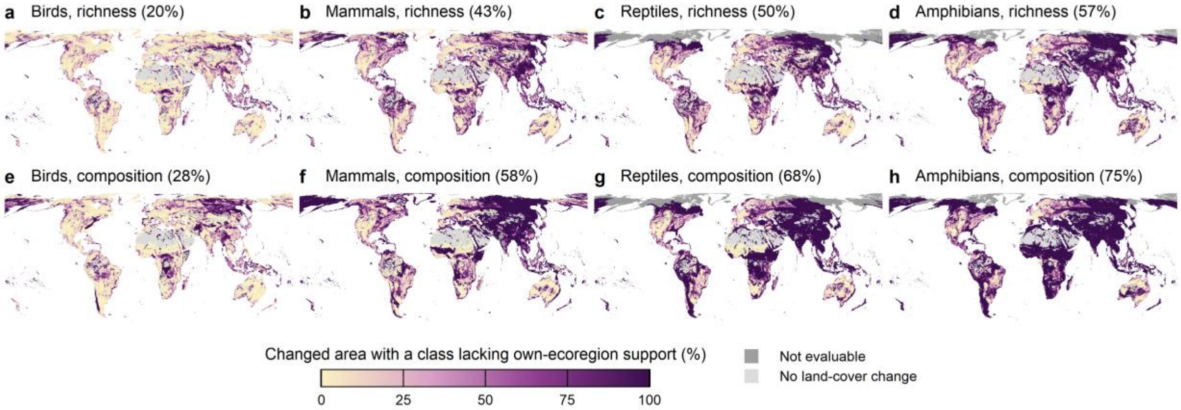


**Supplementary Fig. 3. Extent to which map values rely on classes without support from the data of their own ecoregion.** The upper row (a–d) is expected richness (the maps in Fig. 4a and Extended Data Fig. 4a–c), the lower row (e–h) is composition (the maps in Fig. 4b and Extended Data Fig. 4d–f), and columns are birds, mammals, reptiles and amphibians. Colour shows, for each 0.5° grid cell, the share of the area of cells with changed mixes and values in which at least one class appearing lacks support from the data of its own ecoregion (0–100%; darker is higher). Parentheses in the titles give this share globally. Support means at least five recorded cells in which the class covers at least half the area for expected richness, and at least five pairs with natural cells for composition. Cells in ecoregions with no composition fit (filled wholly with biome means) count as unsupported. For expected richness, each ecoregion is fitted, and only the weights of classes with few data are determined mainly by the biome class effects; the filling for composition relies more heavily on biome information. Coloured cells are not included in the global values in the main text. This flag is provided as a column in the released 5 km values of expected richness and composition and in the coefficient tables (Methods). Pale grey (No land-cover change) is grid cells with no 5-km cell whose land cover changed, dark grey (Not evaluable) is grid cells with changed cells but none with a value, and white is grid cells without land data.

#### Maps and agreement between the two maps

The maps in Fig. 4a,b and Extended Data Fig. 4 of the main text average the values of 5 km cells in 0.5° grid cells of about 50 km (without weights for expected richness and with area weights for composition). Agreement between the maps of change in expected richness and in composition was measured by the Spearman rank correlation and the share of grid cells with opposite signs, over the grid cells drawn in Fig. 4 in which either is non-zero (60,006 grid cells; for composition, the version in Fig. 4b, with cells in ecoregions without a fit filled with biome means). For birds, the rank correlation was 0.51, and signs were opposite in 28% of the grid cells (richness increases while composition is lost, or the reverse). The unfilled version (58,000 grid cells in ecoregions where the composition estimate was fitted) also gave 0.50 and 28%.

### Supplementary Note 5. Retrieval of GBIF records and inclusion criteria

This note gives the conditions under which GBIF records were retrieved and the criteria for inclusion, which removed records that cannot be treated as community surveys. There are three inclusion criteria, described below under Species lists from grid cells georeferenced to a single point, Surveys targeting particular species, and Marine mammals, extinct species and records outside the range.

**Download.** I made one bulk download for each taxon (Supplementary Table 8)[26–29]. The conditions were taxon (key in the GBIF taxonomic backbone)[30], presence of coordinates, no geospatial issue flags, and a licence of CC0 1.0 or CC BY 4.0. Records were not filtered by year, country or basis of record. Because reptiles are not a single class in the GBIF backbone, four orders were combined: Squamata, Testudines, Crocodylia and Rhynchocephalia.

**Supplementary Table 8 Downloads.** The number of records is the number included in the download (before cleaning). The creation date is the date on which GBIF created the download (the date in the GBIF citation), and the DOI of each download is in Supplementary references.

| Taxon | Created (UTC) | Records | Reference |
|---|---|---|---|
| Birds | 2026-09-24 | 2,122,526,659 | GBIF.org[26] |
| Mammals | 2026-09-18 | 34,770,191 | GBIF.org[27] |
| Reptiles | 2026-09-18 | 5,778,177 | GBIF.org[28] |
| Amphibians | 2026-09-18 | 6,176,583 | GBIF.org[29] |

**Cleaning.** I kept records whose latitude and longitude were within valid ranges and not (0, 0); that were resolved in the GBIF backbone to species or a lower rank (identifications to genus or family have an empty species key and are removed automatically); that were presence records (excluding records of "surveyed but absent"); and whose basis of record was neither fossil specimen nor living specimen (captive individuals in zoos and elsewhere). Records were assigned to a 0.05° grid (hereafter 5 km cells), and only land cells with an ecoregion were kept (the 22 cells with an ecoregion but no 2020 land-cover class were not counted as land cells).

**Coordinate precision.** Records were not filtered by coordinate precision (the coordinate uncertainty field). That field is empty for about 90% of bird records, and filtering by precision would lose most bird records. Bias from records with coarse coordinates being assigned to neighbouring cells was not evaluated in this study.

**Recording days.** Records of the same species in the same cell with the same finest available date unit (year, month and day; year and month if the day was missing; year if the month was also missing) were counted as one. The number of recording days of a cell, $T_g$, is the number of distinct values of this unit with records in the cell, and $x_{gs}$ is the number of units on which species $s$ was recorded. Because records without a day count a month or year as one unit, $T_g$ strictly includes units other than days.

**Year of the record.** The estimates use land-cover mixes for 2020, so I used only records from years in which the land cover of the cell can be regarded as the same as in 2020. The land-cover class of each cell is represented by the modal class of its 500 m pixels. Records from a year in 2001–2024 were used if the class of the cell was the same as in 2020 in at least two of the three years centred on that year (the nearest three years for 2001 and 2024), treating a difference in one year as classification noise. Records before 2001 were used only in cells judged to have the same class as in 2020 in every year of 2001–2024. Records from 2025–2026 used the result for 2024. Records with no year or with a year of 2027 or later were excluded. There is no lower limit on the year of a record.

**Species lists from grid cells georeferenced to a single point.** Records that place at one coordinate a species list collected over a large grid cell, rather than an observation at a point, assign to one cell species from an area larger than a 5 km cell. Such a cell looks like one where many species are found in a day of recording and so distorts the richness estimate, which corrects for recording effort (Supplementary Note 8). Three types of such dataset were removed from the input.

**1. Duplicated lists.** Records of the same dataset, coordinates (four decimal places) and date unit form one list. Lists were flagged in which the identical combination of at least ten species appeared at three or more locations (lists with year, month and day were compared within the same date; lists with only year and month, or only year, were compared with each other). Datasets in which such lists accounted for at least half of the records, counted by summed species, were removed as a whole. Compiled datasets not captured by the rule (two animal censuses of Benin) were also removed as a whole.

**2. Coarse grid atlases.** For the distinct coordinates of a dataset I calculated the distance to the nearest other coordinate, and flagged datasets in which at least 0.5 of the distances were within ±5% of the median (an arrangement of grid centres) and the spacing exceeded 6 km. Grids of 6 km or less were kept as point records. The check applied to datasets with 30–200,000 distinct coordinates, and datasets with more than 20,000 were judged on the coordinates within a random window.

**3. Atlases of five-minute grid cells.** Atlases that collect species lists in grid cells of five minutes of latitude and longitude (SABAP2, the Southern African Bird Atlas Project 2[31], and the Nigerian Bird Atlas [two GBIF datasets][32,33]).

**Surveys targeting particular species.** I excluded datasets that contained at most five species of the taxon and that either had a majority of records from machine observation or placed records in at least 50 of the 5 km cells. Datasets whose records were mostly copies of these (republications) were also excluded. If such records were retained, the cells would read as well surveyed but species-poor, unduly reducing the class weights (the expected richness of a cell consisting only of the class; Supplementary Note 8). Datasets flagged by the rule held 0.6% (birds), 15.1% (mammals), 5.2% (reptiles) and 1.2% (amphibians) of all records before cleaning (the judgement also used all records before cleaning). Local single-species surveys confined to fewer than 50 cells are not removed by this rule.

**Marine mammals, extinct species and records outside the range.** All were judged only from GBIF records and taxonomic information. (a) Marine mammals: Cetacea, Sirenia, Phocidae, Otariidae and Odobenidae, and the sea otter (*Enhydra lutris*) and marine otter (*Lontra felina*). (b) Extinct species: species with at least one checklist marking them as extinct and none marking them as not extinct (from the extinction flags of the checklists listing the species in the GBIF species information), and with fewer than five cells having human observation or machine observation records since 2001 (counted on all records before cleaning). (c) Records outside the range: for each species, ecoregions with at least three cells with records were the core, and species–ecoregion combinations were removed if the 0.25° grid cells with records in a non-core ecoregion were more than 2,000 km from the nearest core grid cell. For species with no core, combinations outside the biogeographic realm (RESOLVE realm)[14] with the greatest total of recording days were removed. The thresholds of the rule were not set by agreement with IUCN ranges. For reference, checking against IUCN ranges[20] and BirdLife ranges[21], 71–97% of the records in removed combinations were outside the range, and 0.4–2.6% of combinations within the range were removed in error.

**Supplementary Table 9 Summary of the GBIF data used in the estimates.** Records are GBIF records falling in land 5 km cells (licensed under CC0 or CC BY, after cleaning). Species, cell ×

species combinations (combinations of cell and species with at least one record), cells in the estimate and total recording days are the inputs to the estimates after the inclusion criteria. Total recording days is the sum over all cells of the number of recording days of each cell (see Recording days above). The number of records is also shown in Fig. 1a of the main text.

| Taxon | Records | Cells with records | Species | Cell × species combinations | Cells in the estimate | Total recording days |
|---|---|---|---|---|---|---|
| Birds | 2,074,442,263 | 1,289,059 | 11,262 | 56,659,621 | 1,167,203 | 91,141,658 |
| Mammals | 28,901,258 | 485,982 | 5,426 | 1,925,619 | 390,127 | 5,612,831 |
| Reptiles | 5,336,388 | 343,614 | 9,340 | 959,901 | 277,140 | 1,682,730 |
| Amphibians | 5,657,108 | 274,202 | 6,385 | 664,767 | 214,306 | 1,245,545 |

The number of records and cells with records are counts after cleaning and before the inclusion criteria (one GBIF record counts as one). Species, cell × species combinations (values of $x_{gs}$), cells in the estimate and total recording days (sum of $T_g$) are counts of the inputs to the estimates after the inclusion criteria.

### Supplementary Note 6. Check for exotic species

This note gives the details of the check for exotic species. For birds, the difference in the share of exotic species on lists between anthropogenic land and natural cells is small, so the estimates can include exotic species.

**Exotic species.** To check whether the estimates can include exotic species, I counted the share of species on the lists of the cells used in the composition estimate that are exotic in the country of the cell. Exotic status was determined from national lists of introduced species (Global Register of Introduced and Invasive Species, GRIIS)[34] and the "introduced" flags of GBIF records in that country[26–29]. The richness ratio (Supplementary Note 3) is the ratio of the expected richness of a cell consisting only of an anthropogenic class to that of the natural reference cell, so the amount by which the ratio would move if exotic species were removed depends on the difference, within an ecoregion, in the mean share of exotic species between cells with a cropland (urban) share of at least 0.5 and natural cells (natural-cover share of at least 0.95).

#### Details of the check on the share of exotic species

**Lists.** GRIIS comprises country and territory checklists published in GBIF by the IUCN Invasive Species Specialist Group. Of the 383 lists retrieved, I used 382, excluding one whose archive could not be read (Global Invasive Species Database, with no statement of terms of use) (a DOI will be assigned as a derived dataset on publication)[34,35]. The terms of the 382 lists were CC BY 4.0 or CC0. Species were matched by their keys in the GBIF backbone[30], subspecies were merged into species, and entries listed as absent were excluded. For each country, the list with the most species, among candidates excluding lists of protected areas, lists with titles of the form "region, country" and lists with region codes (US-HI and others), was taken as the national list (for the United States, the mainland list). The GBIF record flag required at least one record of the species in the country marked as introduced (introduced, introducedAssistedColonisation).

**Cells and quantities.** The targets were the cells used in the composition estimate (cells that reached the target coverage in the 752 ecoregions where composition was estimated) and the species tables used in estimating those cells (after the inclusion criteria): 529,567 cells for birds. The country of a cell was determined by the majority of the ISO country codes of its GBIF records. The share of exotic species in cell $g$ is

$$f_g = \frac{|L_g \cap \mathcal{I}_g|}{|L_g|}$$

where $L_g$ is the set of species recorded in cell $g$ and $\mathcal{I}_g$ is the set of species judged exotic in the country of the cell. I compared cells with a cropland share of at least 0.5 (91,877 for birds), cells with an urban share of at least 0.5 (14,763) and natural cells (317,976), using the 2020 land-cover mix. The quantity compared is the difference in the share of exotic species,

$$\mathrm{median}_e\left(\bar{f}_{e,h} - \bar{f}_{e,\mathrm{nat}}\right)$$

where $\bar{f}_{e,h}$ is the mean of $f_g$ in cells of anthropogenic class $h$ (cropland or urban) of ecoregion $e$, $\bar{f}_{e,\mathrm{nat}}$ is the mean of $f_g$ in natural cells, and the median is taken over ecoregions with at least ten cells of each. If this difference was below 0.02 for both cropland and urban, exotic species could be retained; if it was at least 0.02 for either, refitting with native species alone would be required. The value of 0.02 corresponds to a decrease of about 2% in the richness ratio of anthropogenic land if exotic species are removed, and was taken as the size of effect on the richness ratio that can be ignored.

The differences in the share of exotic species for birds (determined from national lists and GBIF flags) were +0.0073 for cropland (215 ecoregions) and +0.0041 for urban (187 ecoregions), both below 0.02, so exotic species were retained in the estimates. National lists alone gave +0.0078 and +0.0044, with the same conclusion. In the version with national lists alone, the most common exotic species, by the share of cropland cells in which they were recorded, were Rock Dove (55%), Common Starling (44%), House Sparrow (42%), Common Pheasant (34%) and Eurasian Collared-Dove (30%), all established in those regions.

**Taxa other than birds (descriptive only).** The same differences were +0.014 for cropland and +0.043 for urban in mammals, +0.006 and +0.032 in reptiles, and 0 and +0.014 in amphibians. Urban values for mammals and reptiles exceeded 0.02. Estimates for taxa other than birds are outside the validation and are only described in the main text.

### Supplementary Note 7. Preparation of land cover and ecoregions

This note shows how the land-cover mixes and ecoregions of the 5 km cells used in the estimates were built.

LC_Type1 (the IGBP classification)[36] of MODIS MCD12Q1 Collection 6.1[37] was processed in Google Earth Engine[38]. The 2020 land-cover mix used in the estimates was built by averaging binary images for each class from 500 m pixels to the 0.05° grid with area weights (the value for each class is the share of the 500 m pixels overlapping the cell that belong to that class). The class of a cell in each year, used to judge the year of a record, is the modal class on the same grid. The mixes used in calculating change from 2020 to 2024 were built separately from pixel counts (Supplementary Note 4). Ecoregions[14] divide the land into 846 units by biogeography and vegetation, and the 14 units above them are called biomes. Ecoregions were rasterized at about 463 m, and each 0.05° cell was assigned its modal ecoregion. Cells without an ecoregion (sea, rock and ice and others) were excluded. The numbers of ecoregions with records are 839 for birds, 814 for mammals, 761 for reptiles and 737 for amphibians.

### Supplementary Note 8. Details of the richness estimate

This note gives the observation model used to estimate the weights and mixing exponent of the richness estimate from GBIF records, its fitting, the judgement of weights used in the results and the construction of intervals. The 17 land-cover classes are divided into 14 natural classes not under human land use (IGBP 1–11 and 15–17: forest, shrubland, savanna, grassland, wetland, snow and ice, barren land and water) and three anthropogenic classes (12 cropland, 13 urban and 14 cropland/natural mosaic).

The richness estimate expresses the expected richness of a 5 km cell in ecoregion $e$ as a function of its land-cover mix $a$ (the area shares $a_k$ of the 17 classes, summing to 1):

$$F_e(a) = \sum_{k=1}^{17} w_{k,e}\, a_k^z, \qquad w_{k,e} = e^{\mu_{k,e}}\{1 - 0.5^{\kappa_e/(\kappa_e+1)}\}$$

The weight $w_{k,e}$ is the expected richness of a cell consisting only of class $k$: the number of species found when that cell is surveyed until sample coverage (the expected share of the species occurrences recorded on a day in that cell that belong to species already on the list)[39] reaches 0.5. This 0.5 is the level that defines the weight and was set separately from the level 0.547 to which lists are standardized in the composition estimate (Supplementary Notes 10, 11). $e^{\mu_{k,e}}$ is the expected richness of a cell consisting only of class $k$ when surveyed completely, which relies more heavily on extrapolation where records are fewer. The weight therefore uses richness up to coverage 0.5, the level chosen as requiring little extrapolation and reached by the records of many cells. $\kappa_e$ is the shape of the gamma distribution that describes how much the daily recording rate varies among species (the smaller it is, the greater the difference between easily and rarely recorded species; see The full observation model below). The factor in braces depends only on $\kappa_e$ and is common within an ecoregion, so ratios of weights within an ecoregion equal ratios of expected richness under complete survey. The mixing exponent $z$ ($0 < z \leq 1$) represents how much richness is added in cells where classes are mixed: there is no addition at 1, and the addition grows as $z$ decreases. This form is a special case of the countryside species–area relationship[40] in which each class has its own group of species and all classes share one exponent. Records are summarized for each 5 km cell $g$ as the number of recording days $T_g$ and the number of recording days on which species $s$ was recorded, $x_{gs}$ (Supplementary Note 5).

#### The full observation model

For cell $g$ of ecoregion $e$, true richness $S_g^*$ and the number of recording days of species $s$, $x_{gs}$, are represented by the following hierarchy (expressing recording probability through an exponential of the recording rate follows Royle & Nichols[2], and the framework of estimating richness from a binomial mixture with recording rates differing among species follows Mao & Colwell[41]):

$$S_g^* \sim \mathrm{Poisson}(M_g), \qquad M_g \sim \mathrm{Gamma}(\text{shape} = \xi_e,\ \text{mean} = \bar{M}_g)$$

$$x_{gs} \mid \nu_{gs} \sim \mathrm{Binomial}(T_g,\ 1 - e^{-\nu_{gs}}), \qquad \nu_{gs} \sim \mathrm{Gamma}(\text{shape} = \kappa_e,\ \text{scale} = \theta_g)$$

Here $M_g$ is the species-pool intensity of the cell (mean true richness, which allows richness to vary among cells with the same mix), $\bar{M}_g = \sum_k e^{\mu_{k,e}} a_{gk}^z$ is its mean determined by the mix, $\nu_{gs}$ is the recording rate of species $s$ per recording day and $\theta_g$ is the daily recording effort of the cell (the larger it is, the more species are recorded in a day). The shape $\kappa_e$ of the variation in recording rates is common to all classes within an ecoregion. Within the model, the evenness of recording rates therefore does not depend on the class, and the ratio of the expected richness of two classes also equals the ratio of the recording-rate-weighted Hill numbers of orders 1 and 2[42] when richness is large. Integrating out $\nu_{gs}$, the probability of $x$ records in $T_g$ days is

$$\Pr(x_{gs} = x \mid \theta_g) = \binom{T_g}{x} \sum_{i=0}^{x} (-1)^i \binom{x}{i} \{1 + (T_g - x + i)\,\theta_g\}^{-\kappa_e}$$

where $i$ is the summation index, and in particular the probability of no records is $(1 + T_g\theta_g)^{-\kappa_e}$. Cells with $T_g \leq 10$ use this equation directly, and cells with $T_g > 10$ are approximated by the negative binomial limit for large $T_g$ (shape $\kappa_e$, mean $\kappa_e T_g \theta_g$; the value 10 is the point up to which numerical error from cancellation in the alternating sum stays sufficiently small). The probability $q_g$ that a species in the cell is recorded at least once is 1 minus this probability of no records, and integrating out $S_g^*$ and $M_g$, observed richness follows a negative binomial distribution:

$$q_g = 1 - (1 + T_g\,\theta_g)^{-\kappa_e}, \qquad S_g \mid \theta_g \sim \mathrm{NB}(\xi_e,\ \bar{M}_g\,q_g)$$

The two arguments of NB are the shape and the mean. Each observed species has $x_{gs}$ following the zero-truncated version of the above binomial–gamma mixture. The likelihood of cell $g$ is

$$\mathcal{L}_g = \int \mathrm{NB}(S_g;\ \xi_e,\ \bar{M}_g q_g) \prod_{s=1}^{S_g} \Pr(x_{gs} \mid x_{gs} \geq 1,\ \theta_g)\,\mathcal{N}(\varepsilon_g;\ 0,\ 1)\,d\varepsilon_g / \Pr(S_g \geq 1)$$

with the product taken over observed species. Daily recording effort is a random effect of the cell:

$$\ln\theta_g = \ln\bar{\theta}_{k(g),e} + \sigma_e\,\varepsilon_g, \qquad \varepsilon_g \sim \mathcal{N}(0,1)$$

Here $k(g)$ is the modal class of the cell in 2020, $\bar{\theta}_{k,e}$ is the centre of daily recording effort for each modal class in ecoregion $e$ and $\sigma_e$ is the variation among cells. The integral over $\varepsilon_g$ was evaluated by adaptive Gauss–Hermite quadrature, and $\Pr(S_g \geq 1)$ by quadrature with 40 points. Cells with only one or two records were kept in the likelihood; their estimates are stabilized by better-recorded cells of the same ecoregion through the random effect of daily recording effort. The species-pool shape and the variation in daily recording effort were bounded to avoid degenerate solutions:

$$\xi_e = 0.1 + \exp(\xi_e^\circ), \qquad \sigma_e = \frac{3}{1 + \exp(-\sigma_e^\circ)}, \qquad \xi_e^\circ, \sigma_e^\circ \sim \mathcal{N}(0,1)$$

As $\xi_e$ approaches 0 the species pool loses its prior information, and as $\sigma_e$ diverges daily recording effort becomes free in each cell; either would produce degenerate estimates. The recording-rate shape has the prior $\ln\kappa_e \sim \mathcal{N}(0,\ 1.5^2)$.

### Hierarchy and priors for sharing information within biomes

To support ecoregion–class combinations with few records, class effects were shared within each biome $m(e)$:

$$\mu_{k,e} = \alpha_e + \beta_{m(e),k} + \psi_k\,\chi_e + \varepsilon_{k,e}^{\mu}, \qquad \varepsilon_{k,e}^{\mu} \sim \mathcal{N}\left(0,\ \tau_\mu^2\right)$$

The terms are as follows. $\alpha_e$ is the intercept of the ecoregion (the overall level of richness; all quantities used in this study are ratios of expected richness within an ecoregion or within-ecoregion comparisons, so it cancels and does not enter the results, and it is therefore not shared within biomes). $\beta_{m,k}$ is the relative species pool of class $k$ in biome $m$ (the quantity shared within the biome). $\chi_e$ is the recording density of the ecoregion (the standardized log of the number of records per cell with records), $\psi_k$ is its effect and $\varepsilon_{k,e}^{\mu}$ is the deviation specific to the ecoregion. Recording density was calculated from records after excluding surveys targeting particular species. The centre of daily recording effort has a hierarchy of the same form (the superscript $\theta$ marks quantities on the daily recording effort side):

$$\ln\bar{\theta}_{k,e} = \alpha_e^{\theta} + \beta_{m(e),k}^{\theta} + \varepsilon_{k,e}^{\theta}, \qquad \varepsilon_{k,e}^{\theta} \sim \mathcal{N}\left(0,\ \tau_\theta^2\right)$$

The ecoregion intercepts have priors $\alpha_e \sim \mathcal{N}(\ln\bar{S}_e + 0.7,\ 1)$ and $\alpha_e^{\theta} \sim \mathcal{N}(-2,\ 3^2)$, where $\bar{S}_e$ is the mean observed richness of cells with records in the ecoregion. The biome class effects are $\beta_{m,k} \sim \mathcal{N}(\bar{\beta}_k,\ 0.3^2)$ and $\beta_{m,k}^{\theta} \sim \mathcal{N}(0,1)$. $\bar{\beta}_k$ was fixed at the mean, across all biomes, of the within-ecoregion relative $\ln w$ of classes that, in estimates fitted to each ecoregion alone, met the precision condition (see Screening the weights below) and had at least five recorded cells in which the class covers at least half the area (in ecoregions with at least two such classes).

The calculation used iterative empirical Bayes, alternating fits by ecoregion (E step) with updates of the biome effects (M step). In the E step, the biome effects, the effects of recording density and the variances ($\beta$, $\beta^{\theta}$, $\psi$, $\tau_\mu$ and $\tau_\theta$, where $\tau$ is the standard deviation among ecoregions) were held fixed to obtain the posterior mode and Hessian of each ecoregion. In the last E step, the uncertainty in the estimates of the hyperparameters was added to the prior standard deviation of $\mu_{k,e}$:

$$\mathrm{SD}\left(\mu_{k,e}\right) = \sqrt{\tau_\mu^2 + \mathrm{SE}\left(\beta_{m(e),k}\right)^2 + \chi_e^2\,\mathrm{SE}(\psi_k)^2}$$

In the M step, the contribution of that round's prior was removed from the Laplace-approximated posterior of each combination[43] to obtain data-only estimates and variances (possible as a difference of precisions because both are normal), and a random-effects model with known variances (mean $\beta_{m,k} + \psi_k\chi_e$ and variance equal to $\tau^2$ plus the variance of the combination) was fitted to update $\beta$, $\psi$ and $\tau$. Combinations with almost no information from the data were left out of the M step, and $\tau$ had a lower bound of 0.1. $\psi_k$ was estimated only for classes with at least ten such combinations. Initial values were the solutions of estimates fitted to each ecoregion alone. After five iterations, the change in $\tau_\mu$ between the last two rounds was at most 0.007.

### Estimating the common mixing exponent

$z$ was handled on the logit scale, $\operatorname{logit} z$, with no prior. In each E step, $z$ was held fixed for the fit by ecoregion, and at the solution I obtained the gradient $\nabla_e$ of the log posterior density in $\operatorname{logit} z$ and the curvature $C_e$ of the profile in which the other parameters are optimized:

$$\mathcal{C}_e = H_{zz} - H_{z\cdot}\, H_{\cdot\cdot}^{-1}\, H_{\cdot z}$$

Here $H$ is the Hessian of the negative log posterior density, the subscript $z$ denotes logit $z$ and the dot denotes the other parameters. The value for the next round is one Newton step using the sums over all ecoregions, logit $z$ minus $\sum_e \nabla_e \,/\, \sum_e \mathcal{C}_e$. Initial values came from estimates fitted to each ecoregion alone. The change in $z$ in the last round was 0.0005 for birds, 0.006 for reptiles, 0.008 for amphibians and 0.010 for mammals.

#### Fitting and convergence checks

The objective function (negative log posterior density) was differentiated automatically with RTMB[44] and TMB[43] in R[45], and the posterior mode was found by numerical optimization. Each ecoregion was fitted in two stages. The first used fixed Gauss–Hermite quadrature with 40 points (nodes and weights calculated with statmod[46]). The second set the centre and width of the quadrature for each cell at that solution and refitted with adaptive quadrature of 10 points. The second stage sometimes failed to converge in ecoregions with $\sigma_e$ near 0, where the prior density of the quadrature diverges; in those cases the converged solution of the first stage was used. In the last round, all ecoregion fits converged. I verified the implementation in two ways: on data in which every cell consists of one class, the objective function matches that of a simple model dividing cells by modal class; and on simulated data, the true $z$ and ratios of weights are recovered.

#### Screening the weights

For the weight of each combination of ecoregion and class, I checked with two conditions that the estimate is sufficiently precise and does not rely too much on extrapolation far from the observations. Weights meeting both are called usable weights, and all calculations in the Methods and main text use only usable weights.

**1. Precision.** The standard deviation of the within-ecoregion relative value (per-draw $\ln w_{k,e}$ minus the mean of $\ln w$ across classes with records) is at most 0.5 (one standard deviation corresponds to a factor of about 1.6). Draws are sets of parameter values drawn at random from the posterior of the estimate (see Intervals below). Uncertainty in the ecoregion intercept $\alpha_e$ and in $\kappa_e$ cancels in ratios, so the judgement used the relative value rather than $\ln w$ itself.

**2. Dependence on extrapolation.** The weights rest on estimates that include undetected species in each cell, and extrapolate far from the observations in poorly recorded cells. I recalculated the weight $w_{k,e}^{\text{cap}}$ with the extrapolation of each cell limited to what would be found by surveying up to three times its current records (with each cell's contribution weighted by $a_{gk}^{z}$ of class $k$), and did not use a weight if $w/w^{\text{cap}} > 2$ (more than half of the weight relying on extrapolation beyond three times). Three times is at the upper end of the proposal that extrapolation be limited to about two to three times the sample[47].

Usable weights meeting both conditions made up 97.8% of combinations of ecoregion and class for birds (9,445 of 9,653), 94.7% for mammals (8,174 of 8,632), 94.8% for reptiles (7,709 of 8,132) and 98.6% for amphibians (7,707 of 7,820).

Separately, I examined how far the data of the ecoregion itself support a class weight. If fewer than five recorded cells have at least half their area in the class, the weight is determined mainly by information from other ecoregions of the same biome. The quantities under alternative natural references (Supplementary Note 3) and global values (Supplementary Note 4) required, in addition to a usable weight, at least five recorded cells in which the class covers at least half the area.

#### Intervals

For each ecoregion, 400 draws were taken from a multivariate normal distribution around the posterior mode (with the inverse Hessian as covariance), and the weights $w_{k,e}$ were calculated for each draw with the equation at the start of this note. Draws are independent among ecoregions. Expected richness and quantities derived from it were calculated with the weights of the same numbered draw, and the 5th to 95th percentiles form the 90% interval. The common $z$ was fixed at its estimate in all draws, so these intervals do not include uncertainty in the estimate of $z$, land-cover misclassification, or the part of the sampling variation in GBIF records that the observation model does not represent. Intervals in the main text additionally include, as model error, the size of the per-ecoregion differences in estimates measured in simulated data (log errors with standard deviation 0.12 for anthropogenic classes and 0.08 for natural classes applied to the draws of the weights; Supplementary Note 1).

### Supplementary Note 9. Validity of the mixing exponent and scale of predicted richness contrasts

This note shows two things using bird surveys not used in the estimation. First, using the mixing exponent estimated from GBIF records in the richness estimate is appropriate (fixing the exponent at 1 lowers performance). Second, the scale of predicted richness contrasts (the size of differences in expected richness among survey units in the same ecoregion) is nearly the same as in the field in North America and larger than in the field in the other two regions.

The richness estimate expresses the expected richness of a 5 km cell in ecoregion $e$ as a function $F_e(a) = \sum_k w_{k,e}\ a_k^z$ of the land-cover mix $a$ (the area shares $a_k$ of the 17 classes, summing to 1). The weight $w_{k,e}$ is the expected richness of a cell consisting only of class $k$, and the mixing exponent $z$ $(0 < z \leq 1)$ represents how much richness is added in cells where classes are mixed, with no addition at 1. This form is a special case of the countryside species–area relationship (Supplementary Note 8)[40]. $z$ was estimated from GBIF records and was 0.880 for birds (Supplementary Note 8).

I checked the following two points in three standardized bird surveys not used in the estimation (SABAP2 in southern Africa, Birdata in Australia and the BBS in North America; Supplementary Note 12). The comparison in the main text (Fig. 2a) uses rank correlations and asks only whether predictors rank survey units correctly. (1) To see whether the size of differences in the estimate matches the size of differences in the field, I used the slope $\gamma$ from regressing log field richness on log expected richness of the estimate. At $\gamma$ of 1, differences in the estimate and in the field are the same size; the further below 1, the larger the differences in the estimate relative to the field, and the further above 1, the smaller. (2) To see whether the effect of added richness in mixed cells $(z < 1)$ is supported in surveys not used in the estimation, I compared the estimate with a version with $z$ fixed at 1 using the same draws of the weights (without refitting). If fixing $z$ at 1 lowers the slope and rank correlation, using the estimated $z$ is appropriate and the added-richness effect is supported by independent data. The equation of the slope and the construction of the compared version are given below under Slopes and the compared version.

The field richness $Y_u$ of survey unit $u$ (SABAP2 square, Birdata 5 km cell or BBS route) is richness rarefied to a common number of surveys.

**Results.** The slopes of the estimate were 0.46, 0.62 and 0.87 (southern Africa, Australia and North America), so differences in the estimate were larger than differences in the field, but nearly the same size in North America (Supplementary Fig. 4a). Fixing the mixing exponent at 1 lowered the slopes to 0.41, 0.52 and 0.60 in all three surveys, and the rank correlations from 0.12, 0.21 and 0.26 to 0.08, 0.11 and 0.11 (Supplementary Fig. 4a,b).

**Conclusions.** Fixing $z$ at 1 lowered the slopes and rank correlations in all three surveys, so using the estimated $z$ is appropriate, and the added richness from mixing classes is supported in surveys not used in the estimation. The scale of predicted richness contrasts is nearly the same as in the field in North America and larger than in the field in the other two regions.

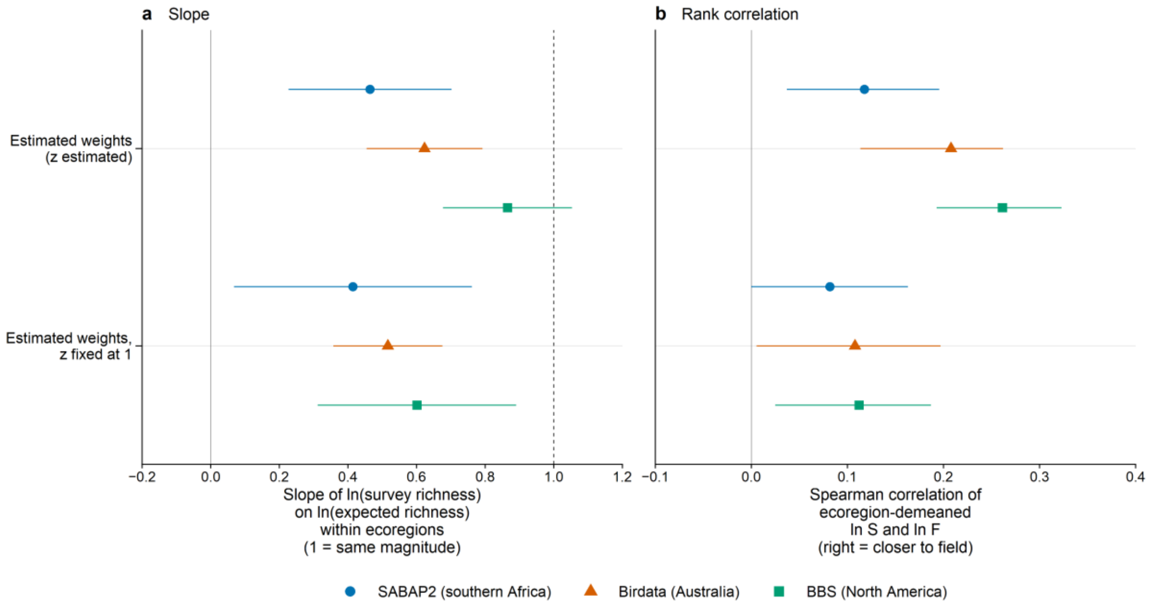


**Supplementary Fig. 4. Validation of the richness estimate in independent bird surveys.** (a) Slope from regressing, within ecoregions, log field richness (rarefied by number of surveys) on log expected richness F of the estimate (after removing ecoregion means and the effects of survey effort and survey-unit area). At 1 (dashed line), differences in the estimate and in the field are the same size; the further below 1, the larger the differences in the estimate relative to the field. Rows are two versions: the estimated weights and mixing exponent (the estimate in the main text; Estimated weights, z estimated) and a version with the mixing exponent fixed at 1 using the same draws of the weights (to see the effect of added richness from mixing classes; not refitted). Points are estimates, and lines are 90% intervals from standard errors clustered by ecoregion (normal approximation). Slopes used all survey units for which expected richness of the estimate can be calculated (3,897, 2,371 and 1,670 units in southern Africa, Australia and North America; 26, 27 and 54 ecoregions). (b) Within-ecoregion rank correlations of the same two versions (Spearman rank correlation between the residuals of field richness [S in the axis title] and of the predictor after subtracting ecoregion means; further right means that survey units within an ecoregion are ranked more correctly). Survey units are the same as in Fig. 2a of the main text (3,105, 1,504 and 1,349). Lines are 90% intervals from a bootstrap over ecoregions (2,000 replicates). In (a) and (b), colours and shapes represent surveys (blue circles SABAP2 in southern Africa, vermilion triangles Birdata in Australia, green squares BBS in North America).

### Slopes and the compared version

Whether differences in the estimate are the same size as differences in the field was examined through the slope $\gamma$ of the following regression (Supplementary Fig. 4a):

$$\ln Y_u = \alpha_e^Y + \gamma \ln F_e(a_u) + \varepsilon_u$$

Here $\alpha_e^Y$ is an ecoregion fixed effect, and terms for the effort and area covariates (Supplementary Note 12) were added to the right-hand side. The slope was estimated on all units for which expected richness can be calculated with the weights of the estimate, so it uses more units than the rank correlations, which use the same units as Fig. 2a of the main text (3,897, 2,371 and 1,670 units [26, 27 and 54 ecoregions] in southern Africa, Australia and North America for the slope; 3,105, 1,504 and 1,349 units for the rank correlations). $F_e$ in the regression is the mean of $F_e(a_u)$ over 400 sets of weights (draws) from the posterior of the estimate. Intervals use a normal approximation from robust standard errors clustered by ecoregion (CR1)[48]. In the version with the mixing exponent fixed at 1, $F_e$ was recalculated with $z = 1$ from the same draws of the weights (without refitting). Rank correlations on the same units (Supplementary Fig. 4b) were calculated, as for Fig. 2a of the main text, by subtracting ecoregion means only.

### Supplementary Note 10. Lists standardized to equal sample coverage

This note shows how the species lists standardized to equal sample coverage, the input to the composition estimate, were built. With standardization by coverage, differences in the daily recording effort of cells do not appear in the lists of cells with the same community (see Estimating sample coverage below). Records are GBIF occurrence records, assigned to a 0.05° latitude–longitude grid (hereafter 5 km cells), with records of the same species in the same cell and date unit (year, month and day; year and month if the day was missing; year if the month was also missing) merged into one (Supplementary Note 5). Each date unit with records in cell $g$ is called a recording day; their number is $T_g$, and the number of recording days on which species $s$ was recorded is $x_{gs}$. Sample coverage[39] is the expected share of the species occurrences recorded on a day in that cell that belong to species already on the list. The list of each cell is extended until coverage reaches the same value $C^*$ (0.547; Supplementary Note 11 explains the choice).

#### Estimating sample coverage

Let the total number of records of cell $g$ (pairs of species and recording day) be $x_{g+} = \sum_s x_{gs}$. The coverage of a list of $t$ days drawn at random from the recording days was estimated, for $t < T_g$, as follows (the formula for incidence records)[49]:

$$\hat{C}_g(t) = 1 - \sum_s \frac{x_{gs}}{x_{g+}} \binom{T_g - x_{gs}}{t} \Big/ \binom{T_g - 1}{t}$$

The sum is over all species recorded in the cell. For the list using all days ($t = T_g$) the denominator above is 0, so coverage was estimated[49] from the numbers of species recorded on exactly one and two days, $Q_1$ and $Q_2$:

$$\hat{C}_g(T_g) = 1 - \frac{Q_1}{x_{g+}} \cdot \frac{(T_g - 1)\, Q_1}{(T_g - 1)\, Q_1 + 2\, Q_2}$$

Coverage increases with the number of days. In a simple model in which the daily recording probability of a species is proportional to the daily recording effort of the cell, $\theta_g$, the coverage of a $t$-day list is determined by the product $t\,\theta_g$ and the structure of the community. In cells with the same community, equal coverage means an equal product $t\,\theta_g$, and differences in daily recording effort vanish. In cells with different communities, what is equalized is the completeness of the list itself.

#### Building the lists by adding recording days

The probability that species $s$ is on a list of $t$ days is given exactly by a hypergeometric probability[47,49]:

$$p_{gs}(t) = 1 - \binom{T_g - x_{gs}}{t} \Big/ \binom{T_g}{t}$$

For target level $C^*$, let $t_g$ be the smallest number of days at which estimated coverage is at least $C^*$. The list is regarded as built from $t_g - 1$ days of records with probability $1 - \omega_g$ and from $t_g$ days with probability $\omega_g$, where $\omega_g$ is set by linear interpolation between the coverages at $t_g - 1$ and $t_g$ days so that expected coverage is exactly $C^*$. The probability that species $s$ is on this list is

$$p_{gs} = (1 - \omega_g)\, p_{gs}(t_g - 1) + \omega_g\, p_{gs}(t_g)$$

From this probability, the list length $F_g' = \sum_s p_{gs}$ and the expected number of species shared by the lists of two cells in the same ecoregion, $G_{gg'} = \sum_s p_{gs}\, p_{g's}$, were calculated exactly. This amounts to calculating, without repeated random sampling, the expectation of the procedure of adding recording days in random order and stopping when coverage reaches the target. Each species counts as one in the number of shared species, but occasional or rare species seldom get onto the lists, so their gains and losses barely move the expected number of shared species. The estimate therefore measures the sharing of well-recorded species.

The cells used are those with at least five recording days whose coverage using all days, $\hat{C}_g(T_g)$, reaches $C^*$. In addition, all classes appearing in the land-cover mix of the cell were required to have a usable richness weight (Supplementary Note 8; the counts in Supplementary Table 10 are after this condition). The lower limit of five days was set to prevent the estimate of coverage from depending strongly on a few days of records. Numbers of cells and list lengths are given in Supplementary Table 10.

**Supplementary Table 10 Size of the data used in the composition estimate.** Level 0.547, cells with at least five recording days. Cells reaching the target are cells whose coverage using all days reaches the target. Cell pairs are pairs 5–500 km apart in the same ecoregion. Estimated ecoregions are those with at least 50 directed cell pairs (observations in Supplementary Note 3; parentheses give those whose fit converged). List length and days used are medians of cells reaching the target.

| Taxon | Cells with at least 5 recording days | Cells reaching the target | Cell pairs | Ecoregions with pairs | Estimated ecoregions | List length (species) | Days used |
|---|---|---|---|---|---|---|---|
| Birds | 617,682 | 529,852 | 6,145,421 | 809 | 752 (734) | 30.3 | 5 |
| Mammals | 145,751 | 127,890 | 1,685,378 | 637 | 458 (447) | 3.4 | 4 |
| Reptiles | 68,249 | 55,153 | 1,146,444 | 541 | 370 (365) | 3.0 | 3 |
| Amphibians | 52,563 | 47,826 | 809,522 | 465 | 291 (290) | 2.7 | 3 |

### Supplementary Note 11. Choice of the sample-coverage level

This note shows that the coverage level 0.547 to which lists are standardized in the composition estimate is the lowest of the six levels examined that meets all three conditions.

The composition estimate extends the species list of each cell until sample coverage (Supplementary Note 10) reaches the same level $C^*$. Lowering the level gives values for more cells but biases the lists towards well-recorded species and makes the estimates of sharing unstable. I considered six candidate levels, two derived only from GBIF records and four preset values (0.30, 0.40, 0.50 and 0.60), and chose the lowest level that meets all three conditions (see The three conditions and the choice below). The level chosen was 0.547, which gives lists of about five days in a typical natural cell.

The quantities in this note are as follows. The field overlap $O_{uv}$ of survey units $u$ and $v$ (SABAP2 square, Birdata 5 km cell or BBS route; Supplementary Note 12) is the presence/absence Sørensen similarity of lists of the same number of surveys drawn from the two units, and the predicted overlap $\hat{O}_{uv}$ is the Sørensen similarity predicted from the land-cover mixes of the two units by the composition estimate (built from the list lengths and sharing probabilities of the two units; Supplementary Note 3). $\bar{O}_u$ is the mean of $O_{uv}$ with natural units of the same ecoregion (survey units with a natural-cover share of at least 0.95), $X_u^{\text{comp}}$ is the corresponding prediction of the composition estimate (the log of the sharing predicted between the survey unit and natural units of the same ecoregion, divided by the sharing predicted between natural units) and $X_u^{\text{nat}}$ is the natural-cover share of the unit.

**Candidates.** Candidates determined only from GBIF records are the medians of the coverage $\hat{C}_g(t)$ of $t$-day lists over all natural bird cells with at least $t$ recording days (all ecoregions): 0.547 (0.5473) at $t$ of 5 and 0.722 at 10. With the preset 0.30, 0.40, 0.50 and 0.60, six levels were compared.

**Comparison between survey sites (condition i).** For all pairs of survey units $u$ and $v$ in the same ecoregion in the three bird surveys (construction of survey units and field species lists in Supplementary Note 12), the field overlap $O_{uv}$ was ranked against the predicted overlap $\hat{O}_{uv}$ and against the land-cover mix similarity $B_{uv} = \sum_k \min(a_{uk}, a_{vk})$. $\hat{O}_{uv}$ is an approximation of the expected Sørensen similarity of the two lists (numerator and denominator each replaced by their expectations), averaged over 400 draws. $B_{uv}$ is the share of area common to the two mixes, a baseline built from land cover alone without GBIF records (equal to $1 - \frac{1}{2}\sum_k |a_{uk} - a_{vk}|$).

**Comparison with natural units (condition ii).** The field quantity is the mean field overlap with natural units, $\bar{O}_u$, the prediction of the estimate is $X_u^{\mathrm{comp}}$ and the baseline is the natural-cover share of the unit, $X_u^{\mathrm{nat}}$.

**Evaluation.** The construction of rank correlations and intervals and the combination of the three surveys are the same as in Supplementary Notes 12 and 13. In comparisons between sites, the covariates were the log distance between the two units and the sum and absolute difference of the efforts of the two units.

**The three conditions and the choice.** (i) Predicted overlap must outrank mix similarity in the comparison between sites: the lower end of the 90% interval of their difference in rank correlation, combined across the three surveys, must be above 0. In addition, the point estimate of this difference must not fall by 0.02 (a preset tolerance) or more from its value at 0.722, the highest candidate level. (ii) In the comparison with natural units, the lower end of the combined 90% interval of the difference in rank correlation between the prediction and natural-cover share is above 0. (iii) In the simulated data (Supplementary Note 1), the ecoregion medians of the ratio of retention from records with daily effort halved on anthropogenic land to that from equal-effort records all lie within 0.95–1.05 for cropland, urban and mosaic. The lowest level meeting all three is chosen. Conditions (i) and (ii) use the same three surveys as the comparison with existing indicators in the main text, so the choice of level is not fully independent of the comparison of composition in the main text. However, the differences due to the level chosen are small (see Results below; in (i) the difference is +0.086 at 0.547 and +0.080 at 0.722).

**Results.** All three conditions were met by 0.722, 0.60 and 0.547, and the lowest, 0.547, was chosen. The level 0.50 failed condition (i), and 0.40 and 0.30 failed condition (iii). Versions of 0.40 and 0.30 with the minimum number of recording days lowered to three also failed conditions (i) and (iii), respectively. The values at 0.547 were +0.086 for (i) (90% interval 0.047–0.125; +0.080 at 0.722), +0.120 for (ii) (0.075–0.166), and 0.97 for cropland, 0.96 for urban and 0.97 for the mosaic for (iii).

### Supplementary Note 12. Independent bird surveys and field quantities

This note shows how the survey units of the three standardized bird surveys not used in the estimation (SABAP2 in southern Africa, the Southern African Bird Atlas Project 2[31]; Birdata in Australia[50]; and the BBS in North America[51]) and the field quantities built from them were constructed. None of the three surveys is in the input to the estimates. SABAP2 records are in the GBIF bird download but were removed from the input as atlases of five-minute grid cells (Supplementary Note 5). Birdata records are licensed CC BY-NC and so are not in the input, which is filtered by licence. BBS route surveys are not in the GBIF download used in the estimation. The field richness $Y_u$ of survey unit $u$ is the expected number of species when a fixed number of surveys $T^*$ (four for SABAP2, ten for Birdata and three for the BBS) is drawn from those of the unit; the field overlap $O_{uv}$ is the presence/absence Sørensen similarity of lists of $T^*$ surveys drawn from each of two units, averaged over 50 redraws; and the field composition similarity $\bar{O}_u$ is the mean of $O_{uv}$ with the set $\mathcal{V}_u$ of natural units of the same ecoregion (survey units with a natural-cover share of at least 0.95).

#### Survey units and inclusion conditions by survey

**Survey protocols.** One survey (card) of SABAP2 is a species list made by an observer who spends at least two hours surveying as many environments as possible within a five-minute square. One survey of Birdata covers 2 ha in 20 minutes within a 5 km cell[52]. One survey of the BBS counts birds for three minutes at each of 50 stops along a route of about 40 km, once a year.

**SABAP2.** I retrieved 405,587 full-protocol survey cards (19,607 squares, 2007–2025) from the Africa Bird Atlas Project API[53]. Species lists were attached by matching uniquely on square, start date and number of records (the total species of the card) with the SABAP2 records in GBIF (bird download)[26], for 325,503 cards (80.3%). These records were removed from the input to the estimates. I used squares with at least four cards in the period 2017–2023. Land-cover mixes were built from the 2020 classes of 0.005° pixels whose centres fall in the square. The ecoregion was determined from the 5 km cells in the square (one to four), and only squares with all cells in the same ecoregion were used. The effort covariate is the log of the mean observation time of the cards, and the area covariate is the log cosine of latitude.

**Birdata.** Among the records of 2 ha, 20-minute surveys retrieved from GBIF (4,411,679 records)[50], 154,270 surveys from 2017–2023 with a survey identifier, a single coordinate and uncertainty of at most 500 m were assigned to 5 km cells. I used cells with at least ten surveys and at least three survey sites in the cell (to exclude cells with only repeats of the same plot). Land-cover mixes were built from the 2020 classes of the 100 pixels of the cell. The effort covariate is the log of the mean number of observers per survey, and the area covariate is the log cosine of latitude.

**BBS.** I used standard surveys from the 2026 release (1966–2025, CC0), excluding 74 species codes for unidentified taxa and hybrids. Route locations were taken from the 1999 National Atlas route lines[54], and I used 2,905 routes whose start point was within 1 km of the line and whose length was 30–50 km (routes established or changed after 1999 cannot be used). Land-cover mixes were built from the 2020 classes of 0.005° pixels with centres in strips 400 m on both sides of the line. The ecoregion was determined by a majority weighted by line length, and only routes with a majority share of at least 0.8 were used. I used routes with at least three surveys in the period 2017–2023 (no surveys in 2020). The effort covariate is the log number of surveys; route lengths are standardized, so no area covariate was included.

**Supplementary Table 11 Numbers of survey units used in the comparisons (numbers of ecoregions in parentheses).**

| Set | SABAP2 | Birdata | BBS |
|---|---|---|---|
| Slope of the richness estimate (Supplementary Note 9) | 3,897 (26) | 2,371 (27) | 1,670 (54) |
| Rank correlation and effect size for richness (Fig. 2a, Extended Data Fig. 1a) | 3,105 (25) | 1,504 (23) | 1,349 (51) |
| Same, BII and BHI maps | 3,105 | 1,501 | 1,349 |
| Same, LIFE | 3,094 (25) | 1,456 (23) | 1,317 (50) |
| Same, IUCN AoH richness | 3,105 | 1,502 | 1,349 |
| Rank correlation for composition (presence/absence; units with values for the BII and BHI maps in Fig. 2b) | 3,078 (24) | 1,501 (23) | 912 (36) |
| Rank correlation for composition (abundance) | 3,078 | 1,379 | 912 |
| Composition, LIFE | 3,067 | 1,453 | 911 |
| Composition, IUCN AoH richness | 3,078 | 1,500 | 912 |

The composition estimate can be built only for the units of the composition comparison, which have natural units in the same ecoregion. The composition-estimate row of Fig. 2a was therefore calculated on units with composition-estimate values (3,078, 1,504 and 912; these include three units without values for the BII and BHI maps, so the number for Birdata exceeds the 1,501 of the composition row). LIFE and IUCN AoH richness have fewer units because of the extents of the maps. The abundance-based measure has no value for units with too few surveys counting the abundance of all species in Birdata.

### Details of rarefaction and field quantities

Field richness $Y_u$ was calculated from the incidence frequencies of surveys within the period of the unit by the following equation, where $T_u$ is the number of surveys of the unit and $x_{us}$ is the number on which species $s$ was recorded[47,49]:

$$Y_u = \sum_s \left[1 - \binom{T_u - x_{us}}{T^*} / \binom{T_u}{T^*}\right]$$

The species lists for composition were built by drawing $T^*$ surveys at random without replacement for each unit (four cards for SABAP2, ten surveys for Birdata and three years for the BBS), and the Sørensen similarity of two units, $2|L_u \cap L_v|/(|L_u| + |L_v|)$, where $L_u$ is the species list of unit $u$, was averaged over 50 redraws. The abundance-based measure is the mean, over natural units $v \in \mathcal{V}_u$, of the Bray–Curtis similarity

$$\mathrm{BC}_{uv} = 1 - \frac{\sum_s |N_{us} - N_{vs}|}{\sum_s (N_{us} + N_{vs})}$$

Here $N_{us}$ is the abundance measure of species $s$ in unit $u$: the mean count per survey in the BBS, the mean count per survey with counts for all records in Birdata (only units with at least five such surveys), and the reporting rate (the share of cards recording the species) in SABAP2.

#### Regression for the composition comparison

For the composition comparison, I subtracted ecoregion means from the field quantity, each predictor compared (the two estimates of this study, the existing indicators and natural-cover share) and the covariates, regressed the field quantity and each predictor on the covariates by least squares, and calculated the Spearman rank correlation between the residuals. This is equivalent to removing ecoregion fixed effects and covariates together. The covariates are the mean over $\mathcal{V}_u$ of the log great-circle distance to natural units (distances below 1 km set to 1 km) and the effort covariate of each survey (above). Distance was removed because compositional similarity falls strongly with the distance to the units compared regardless of land cover; richness does not have this problem, so the rank correlation for the richness comparison (Fig. 2a) subtracted ecoregion means only. For richness, the effort and area covariates entered the regressions for effect size (Supplementary Note 13) and slope (Supplementary Note 9).

### Supplementary Note 13. Details of intervals and effect sizes

This note supplements the construction of intervals for differences in rank correlation and of the upper limit of effect size when comparing the estimates and existing indicators in independent bird surveys.

**Symbols.** For each survey $j$ (of the three bird surveys), I write the rank correlation of predictor $X$ (how correctly the predictor ranks differences among survey units in the same ecoregion) as $r_{X,j}$, that of the estimate from this study as $r_{\text{est},j}$, and their difference as $\Delta_j = r_{\text{est},j} - r_{X,j}$. $\text{SE}_j$ is the standard error of $\Delta_j$, and $\bar{\Delta}$ is the precision-weighted mean of $\Delta_j$ over the three surveys:

$$\bar{\Delta} = \frac{\sum_j \text{SE}_j^{-2}\, \Delta_j}{\sum_j \text{SE}_j^{-2}}$$

Effect size $\text{SS}_X$ expresses where the variance explained by predictor $X$ lies between the baseline (natural-cover share) and the upper limit (it has the same form as a skill score in forecast evaluation, hence the symbol SS):

$$\text{SS}_X = \frac{\Delta R_X^2 - \Delta R_{\text{nat}}^2}{\Delta R_{\text{max}}^2 - \Delta R_{\text{nat}}^2}$$

Here $\Delta R_X^2$ is the increase in the within-ecoregion coefficient of determination when predictor $X$ is added to a model with only ecoregion fixed effects and covariates for effort and area, $\Delta R_{\text{nat}}^2$ is the same increase for natural-cover share, and $\Delta R_{\text{max}}^2$ is the increase for a model with the shares of the 17 land-cover classes as separate explanatory variables (the upper limit). The upper limit indicates how much a prediction based only on the land-cover mix can explain: no such prediction is likely to explain more than this model, which fits the mix freely to the data (it was evaluated by cross-validation, to compare explanatory power for new units; see The upper limit of effect size below). An effect size of 0 means the same as the baseline, and 1 means reaching the upper limit.

#### Details of intervals and precision weights

In the cluster bootstrap, the ecoregions of each survey were resampled with replacement, and all units of the selected ecoregions were included. Replicate $i$ used estimation draw $((i-1) \bmod 400) + 1$, with the values of maps, coefficient versions and natural-cover share fixed. The difference $\Delta_j$ is the difference between the two rank correlations of the same replicate, and intervals come from its distribution. $\text{SE}_j$ in the precision-weighted mean $\bar{\Delta}$ is the width of the 90% interval of each survey divided by $2 \times 1.645$ (normal approximation), treating the bootstraps of the three surveys as independent. The lower end of its 90% interval is $\bar{\Delta} - 1.645(\sum_j \text{SE}_j^{-2})^{-1/2}$. Point estimates are reported for rank correlations and their differences, and bootstrap medians for effect sizes.

**The upper limit of effect size**

$\Delta R_X^2$ and $\Delta R_{\text{nat}}^2$ are in-sample values, and $\Delta R_{\max}^2$ is a cross-validated value. The within-ecoregion coefficient of determination is

$$R_{\text{within}}^2 = 1 - \frac{\text{SSR}}{\sum_u \left(\ln Y_u - \overline{\ln Y}_{e(u)}\right)^2}$$

where $Y_u$ is the field richness of survey unit $u$ (Supplementary Note 12; for composition, the field quantity is the field composition similarity $\bar{O}_u$ and the covariates are effort and the log distance to natural units, plus the log cosine of latitude for surveys in which unit area varies with latitude), SSR is the residual sum of squares of the model with fixed effects, $e(u)$ is the ecoregion of unit $u$ and $\overline{\ln Y}_{e(u)}$ is the mean of that ecoregion. The upper-limit model uses as explanatory variables the shares of classes with positive shares in at least 20 units (omitting one class because shares sum to 1). The units of each ecoregion were split into five, the model with fixed effects was fitted to four, and the remaining one was predicted (with the intercept from the mean of the training side). The model with covariates only was cross-validated with the same splits, and the difference is $\Delta R_{\max}^2$. Results were averaged over 20 different splits. Because the upper limit is a cross-validated value, the effect size of a predictor fitted in sample can exceed 1.

### Supplementary Note 14. Fitting the three taxa other than birds

This note gives the details of applying the level and models used for birds to mammals, reptiles and amphibians, and the reason for setting the minimum number of recording days per cell to five.

**Level and z.** The same $C^*$ (0.547) as for birds was used. For taxa other than birds, most cells with at least five recording days already reach a higher coverage than this value, and what largely changes the number of usable cells is not the target value but the minimum number of recording days. $z$ was taken directly from the richness estimate of each taxon (0.95 for mammals, 0.94 for reptiles and 0.97 for amphibians).

**Minimum number of recording days.** I fitted two versions, with minimum numbers of five and three days. The three-day version increased the share of changed cells from 2020 to 2024 with map values by 3.8–9.7 percentage points but moved the retention of anthropogenic classes by 12–17% in the ecoregion median (absolute log ratio), so I chose five days. The five-day version converged in 97.6%, 98.6% and 99.7% of ecoregions (mammals, reptiles and amphibians).

**Check of shrinkage.** The two conditions of Supplementary Note 3 were met for all three taxa (share satisfying the inequality 0.928–0.943; posterior standard deviation in ecoregions with few observations 22–30% of that of the independent estimates).

### Supplementary references

The references cited in the Supplementary Information are numbered here separately from those of the main text and Methods.


1. Lumbierres, M. et al. Translating habitat class to land cover to map area of habitat of terrestrial vertebrates. *Conserv. Biol.* **36**, e13851 (2022).
2. Royle, J. A. & Nichols, J. D. Estimating abundance from repeated presence–absence data or point counts. *Ecology* **84**, 777–790 (2003).
3. Schipper, A. M. et al. Projecting terrestrial biodiversity intactness with GLOBIO 4. *Glob. Change Biol.* **26**, 760–771 (2020).
4. Hoskins, A. J. et al. BILBI: supporting global biodiversity assessment through high-resolution macroecological modelling. *Environ. Model. Softw.* **132**, 104806 (2020).
5. Scherer, L. et al. Biodiversity impact assessment considering land use intensities and fragmentation. *Environ. Sci. Technol.* **57**, 19612–19623 (2023).
6. Gassert, F., Mazzarello, J. & Hyde, S. *Global 100m Projections of Biodiversity Intactness for the Years 2017–2025 (v1.1)* (Impact Observatory & Vizzuality, 2026); https://source.coop/vizzuality/biodiversity-intactness-100m-v1-1

7. De Palma, A. et al. Annual changes in the Biodiversity Intactness Index in tropical and subtropical forest biomes, 2001–2012. *Sci. Rep.* **11**, 20249 (2021).
8. Eyres, A. et al. LIFE: a metric for mapping the impact of land-cover change on global extinctions. *Phil. Trans. R. Soc. B* **380**, 20230327 (2025).
9. Mair, L. et al. A metric for spatially explicit contributions to science-based species targets. *Nat. Ecol. Evol.* **5**, 836–844 (2021).
10. Schipper, A. M. et al. Impact factors for quantifying country-level terrestrial biodiversity intactness footprints (IBIF). *Sci. Data* **12**, 1660 (2025).
11. Ware, C. et al. *Global Biodiversity Indicator Data for BHI, BERI, PARC-representativeness, PARC-connectedness and Ecosystem Condition (2000–2024)* (CSIRO, 2026); https://doi.org/10.25919/3aka-y730
12. Scherer, L. et al. *Biodiversity Impact Assessment Considering Land Use Intensities and Fragmentation, v1.0.1* (Zenodo, 2026); https://doi.org/10.5281/zenodo.20617889
13. Olson, D. M. et al. Terrestrial ecoregions of the world: a new map of life on Earth. *BioScience* **51**, 933–938 (2001).
14. Dinerstein, E. et al. An ecoregion-based approach to protecting half the terrestrial realm. *BioScience* **67**, 534–545 (2017).
15. De Palma, A. et al. *The Biodiversity Intactness Index Developed by The Natural History Museum, London, v2.1.1* (Natural History Museum, 2024); https://doi.org/10.5519/k33reyb6
16. Eyres, A., Dales, M. W. & Madhavapeddy, A. *LIFE: a Metric for Mapping the Impact of Land-cover Change on Global Extinctions, v1.01* (Zenodo, 2024); https://doi.org/10.5281/zenodo.14945383
17. Lumbierres, M. et al. Area of Habitat maps for the world's terrestrial birds and mammals. *Sci. Data* **9**, 749 (2022).
18. IUCN. *Species Richness from AoH Rasters, Birds (Based on IUCN Red List Version 2021-3)* (IUCN, 2023); https://www.iucnredlist.org/resources/spatial-data/other-spatial-downloads
19. Brooks, T. M. et al. Measuring terrestrial Area of Habitat (AOH) and its utility for the IUCN Red List. *Trends Ecol. Evol.* **34**, 977–986 (2019).
20. IUCN. *The IUCN Red List of Threatened Species, Version 2024-2* (IUCN, 2024); https://www.iucnredlist.org
21. BirdLife International & Handbook of the Birds of the World. *Bird Species Distribution Maps of the World, Version 2024.2* (BirdLife International, 2024); https://datazone.birdlife.org/contact-us/request-our-data
22. Danielson, J. J. & Gesch, D. B. *Global Multi-resolution Terrain Elevation Data 2010 (GMTED2010)*. U.S. Geological Survey Open-File Report 2011-1073 (USGS, 2011); https://doi.org/10.3133/ofr20111073
23. HBW & BirdLife International. *Handbook of the Birds of the World and BirdLife International Digital Checklist of the Birds of the World, Version 9* (BirdLife International, 2024); https://datazone.birdlife.org/about-our-science/taxonomy
24. Dornelas, M. et al. Assemblage time series reveal biodiversity change but not systematic loss. *Science* **344**, 296–299 (2014).
25. Chandler, R. E. & Bate, S. Inference for clustered data using the independence loglikelihood. *Biometrika* **94**, 167–183 (2007).
26. GBIF.org. GBIF occurrence download (birds; CC0 and CC BY 4.0). *GBIF* https://doi.org/10.15468/dl.pwdmqt (2026).
27. GBIF.org. GBIF occurrence download (mammals; CC0 and CC BY 4.0). *GBIF* https://doi.org/10.15468/dl.jgsvdv (2026).
28. GBIF.org. GBIF occurrence download (reptiles; CC0 and CC BY 4.0). *GBIF* https://doi.org/10.15468/dl.9sqgwd (2026).
29. GBIF.org. GBIF occurrence download (amphibians; CC0 and CC BY 4.0). *GBIF* https://doi.org/10.15468/dl.s272tx (2026).

30. GBIF Secretariat. *GBIF Backbone Taxonomy* (GBIF, 2023); https://doi.org/10.15468/39omei
31. Brooks, M. et al. The African Bird Atlas Project: a description of the project and BirdMap data-collection protocol. *Ostrich* **93**, 223–232 (2022).
32. Chaskda, A. et al. *Nigeria Bird Atlas (Full Protocol Submissions), Version 1.8* (A.P. Leventis Ornithological Research Institute, 2022); https://doi.org/10.15468/hsvds6
33. Danmallam, B. A. et al. *Nigerian Bird Atlas Project (Adhoc Records), Version 1.46* (A.P. Leventis Ornithological Research Institute, 2026); https://doi.org/10.15468/fcp87c
34. Pagad, S., Genovesi, P., Carnevali, L., Schigel, D. & McGeoch, M. A. Introducing the Global Register of Introduced and Invasive Species. *Sci. Data* **5**, 170202 (2018).
35. GBIF.org. Derived dataset (382 GRIIS country and territory checklists). *GBIF* (2026); DOI assigned on publication.
36. Sulla-Menashe, D., Gray, J. M., Abercrombie, S. P. & Friedl, M. A. Hierarchical mapping of annual global land cover 2001 to present: the MODIS Collection 6 Land Cover product. *Remote Sens. Environ.* **222**, 183–194 (2019).
37. Friedl, M. & Sulla-Menashe, D. *MODIS/Terra+Aqua Land Cover Type Yearly L3 Global 500m SIN Grid V061* (NASA EOSDIS Land Processes DAAC, 2022); https://doi.org/10.5067/MODIS/MCD12Q1.061
38. Gorelick, N. et al. Google Earth Engine: planetary-scale geospatial analysis for everyone. *Remote Sens. Environ.* **202**, 18–27 (2017).
39. Chao, A. & Jost, L. Coverage-based rarefaction and extrapolation: standardizing samples by completeness rather than size. *Ecology* **93**, 2533–2547 (2012).
40. Pereira, H. M. & Daily, G. C. Modeling biodiversity dynamics in countryside landscapes. *Ecology* **87**, 1877–1885 (2006).
41. Mao, C. X. & Colwell, R. K. Estimation of species richness: mixture models, the role of rare species, and inferential challenges. *Ecology* **86**, 1143–1153 (2005).
42. Chao, A., Chiu, C.-H. & Jost, L. Unifying species diversity, phylogenetic diversity, functional diversity, and related similarity and differentiation measures through Hill numbers. *Annu. Rev. Ecol. Evol. Syst.* **45**, 297–324 (2014).
43. Kristensen, K., Nielsen, A., Berg, C. W., Skaug, H. & Bell, B. M. TMB: automatic differentiation and Laplace approximation. *J. Stat. Softw.* **70**, 1–21 (2016).
44. Kristensen, K. RTMB: 'R' bindings for 'TMB'. R package version 2.0 https://doi.org/10.32614/CRAN.package.RTMB (2026).
45. R Core Team. *R: A Language and Environment for Statistical Computing, Version 4.6.1* (R Foundation for Statistical Computing, 2026); https://www.R-project.org/
46. Smyth, G. et al. statmod: statistical modeling. R package version 1.5.2 https://doi.org/10.32614/CRAN.package.statmod (2026).
47. Colwell, R. K. et al. Models and estimators linking individual-based and sample-based rarefaction, extrapolation and comparison of assemblages. *J. Plant Ecol.* **5**, 3–21 (2012).
48. Cameron, A. C. & Miller, D. L. A practitioner's guide to cluster-robust inference. *J. Hum. Resour.* **50**, 317–372 (2015).
49. Chao, A. et al. Rarefaction and extrapolation with Hill numbers: a framework for sampling and estimation in species diversity studies. *Ecol. Monogr.* **84**, 45–67 (2014).
50. GBIF.org. GBIF occurrence download (Birdata of BirdLife Australia; 2 ha, 20-minute surveys). *GBIF* https://doi.org/10.15468/dl.2qhj7a (2026).
51. Ziolkowski, D., Lutmerding, M., Skalos, S. M., English, W. B. & Hudson, M.-A. R. 2026 release – North American Breeding Bird Survey dataset (1966–2025). *U.S. Geological Survey* https://doi.org/10.5066/P144YU3S (2026).
52. Loyn, R. H. The 20 minute search – a simple method for counting forest birds. *Corella* **10**, 105–108 (1986).
53. FitzPatrick Institute of African Ornithology. Southern African Bird Atlas Project 2. *Africa Bird Atlas Project* https://api.birdmap.africa (University of Cape Town).

54. USGS Patuxent Wildlife Research Center. *Breeding Bird Survey Route Locations for Lower 48 States, 1966–1998* (National Atlas of the United States, 1999).
